## Supporting Information for "Natural variation in the *Caenorhabditis elegans* egg-laying circuit modulates an intergenerational fitness trade-off"

**This PDF file includes:**
Figures S1 to S7
Table S1 to S7
Description of separate file Data S1 (raw data in Excel format)

**Additional supporting information:**
Data S1: Raw data (separate Excel file)

### Supporting Figures

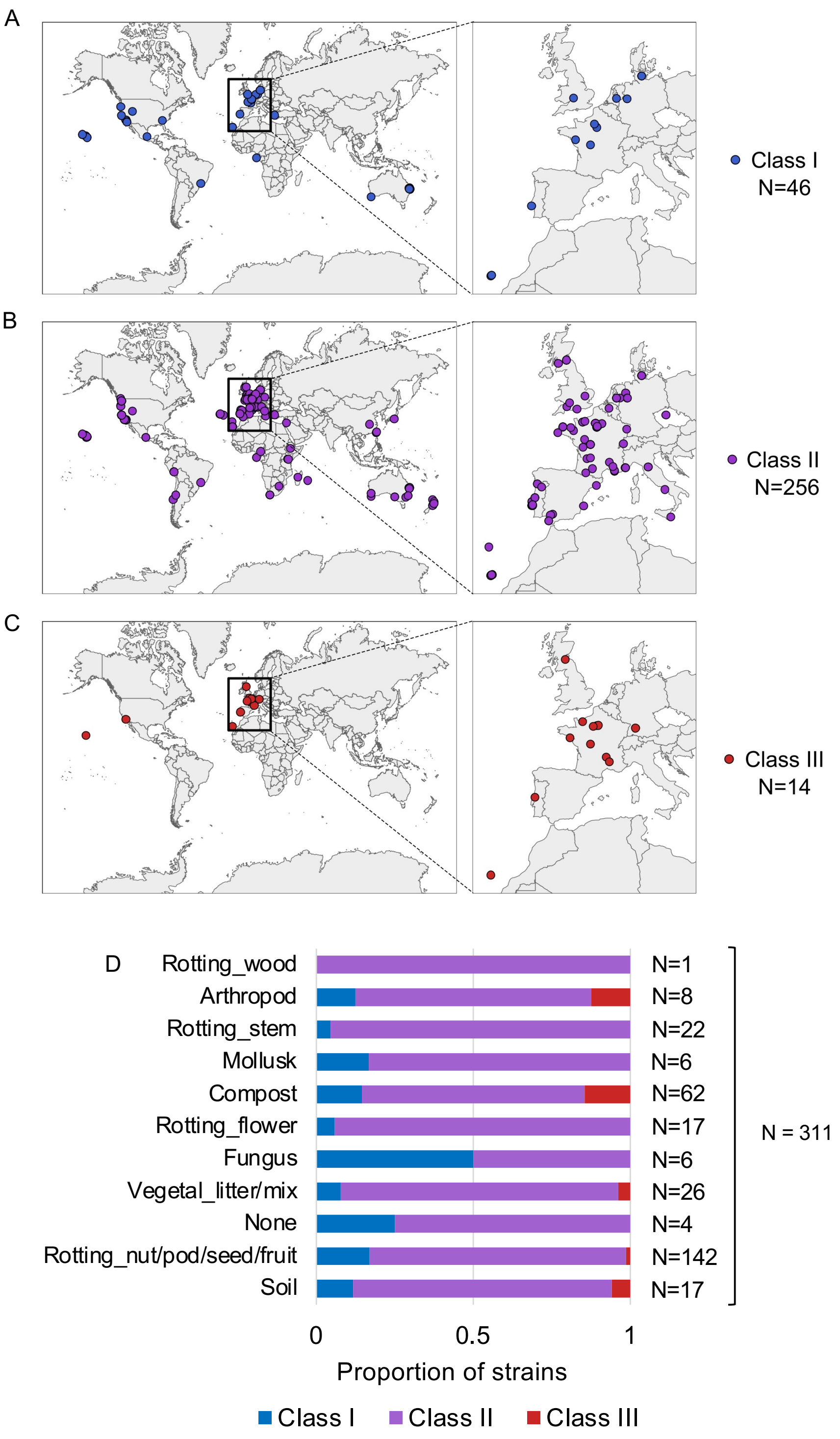

**Figure S1 Natural variation of *C. elegans* egg number *in utero***
**A** Geographic distribution of Class I isolates: < 10 eggs *in utero*, low retention (N = 34). **B** Geographic distribution of Class II isolates: 10-25 eggs *in utero*, canonical retention (N = 230). **C** Geographic distribution of Class III isolates: > 25 eggs *in utero*, high retention (N = 14). **D** Frequency of strains with different egg retention phenotypes (Class I to III) in each substrate category. Strain information was obtained from the CeNDR website: https://www.elegansvariation.org. N = 311 (for five strains there was no substrate information available).

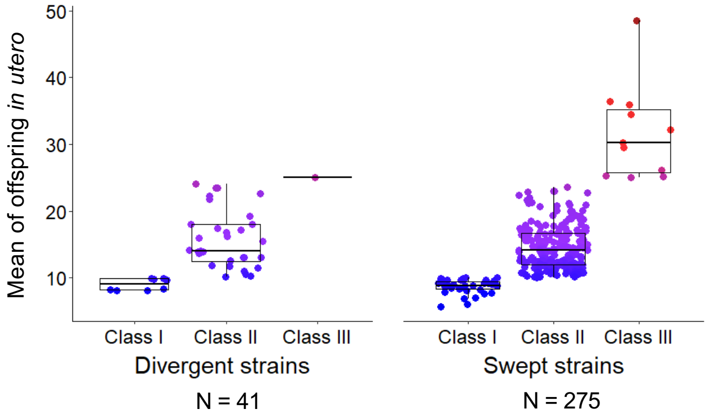

**Figure S2 Egg number *in utero* in swept versus divergent *C. elegans* strains**
Strains, scored for egg number *in utero* at L4+48h (Fig. 1A), were categorized as swept (N=275) or as divergent strains (N=41) following previous classifications (Gilbert et al., 2022). In brief, strains are categorized as swept if any of chromosomes I, IV, V, or X contained greater than or equal to 30% of the same haplotype; strains not among the swept strains are classified as divergent (Andersen et al., 2012; Gilbert et al., 2022; Lee et al., 2021). 13 out of the 14 Class III strains exhibited a swept haplotype.

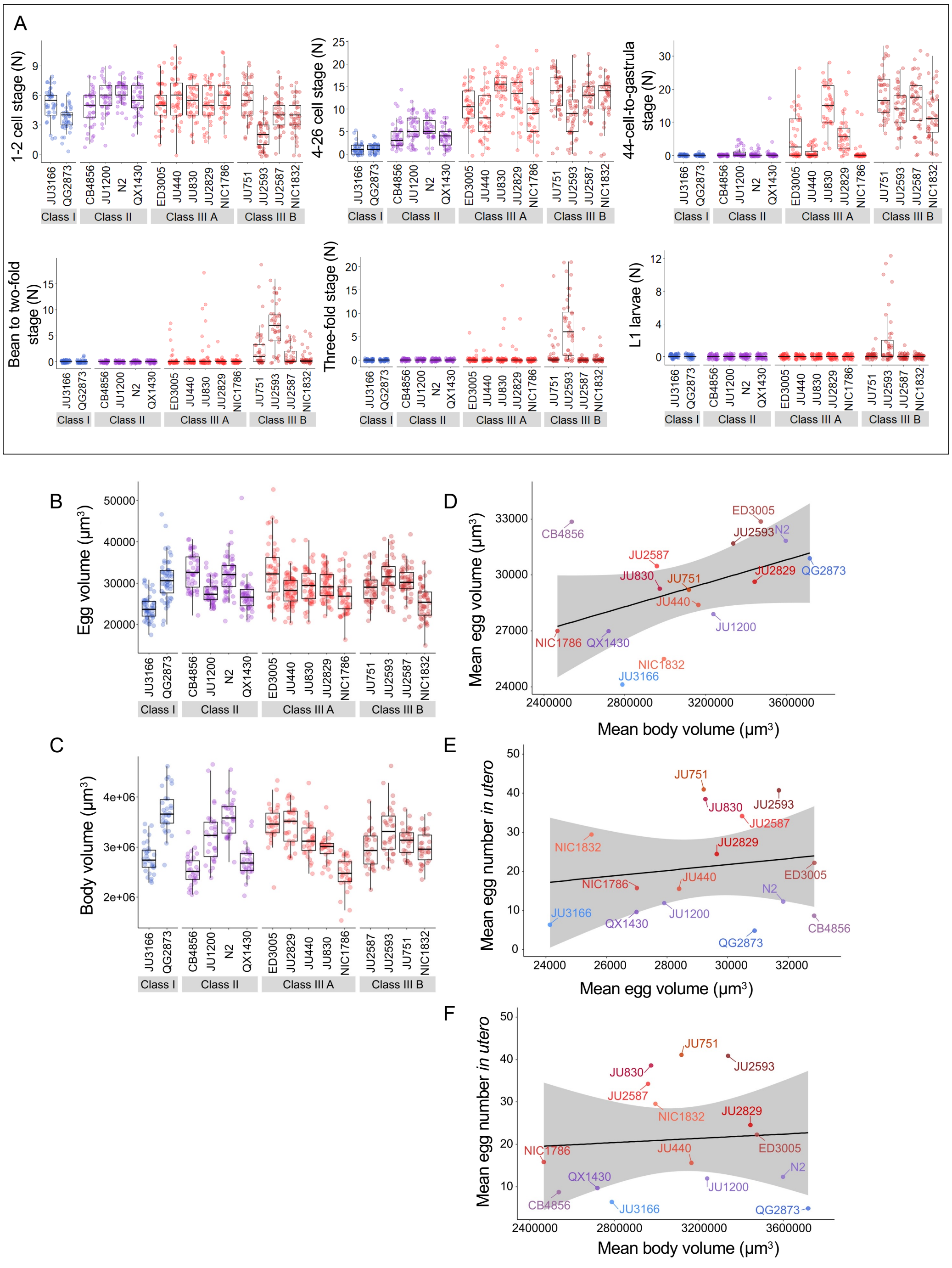

**Figure S3 Temporal progression of egg retention and internal hatching**

**A** Age distribution of embryos within eggs retained *in utero* of hermaphrodites (L4 + 30h) in the 15 focal strains, including the N2-derived strain QX1430 (Andersen et al., 2015). Embryonic stages were divided into five age groups according to the following characteristics using Nomarski microscopy (Hall et al., 2007): 1-2 cell stage, 4-26 cell stage, 44 cell to gastrula stage, bean to two-fold stage, three-fold stage, L1 larva. N = 36-40 individuals were scored per strain; data obtained from same individuals scored for number of eggs *in utero* (Fig. 2A). **B** Egg size (volume) in the 15 focal strains with variable egg retention, measured on laid eggs of mixed-age adult populations. Strains differed significantly in egg size (Kruskal-Wallis Test, $\chi^2$ = 223.97, df = 14, P < 0.0001). N = 50-69 eggs per strain. **C** Body size (volume) measured in the 15 focal strains; selfing hermaphrodites at first fertilization (1-2 eggs *in utero*). Strains differed significantly in body size (Kruskal-Wallis Test, $\chi^2$ = 227.45, df = 14, P < 0.0001). N = 47-51 individuals per strain. **D** Marginally significant positive correlation between mean egg size and mean body size (volume; at first fertilization, 1-2 eggs *in utero*) across the 15 focal strains with divergent egg retention ($\rho_{Spearman}$ = 0.50, P = 0.06). **E** No correlation between mean egg size and mean egg retention (at L4 + 30h) across the 15 focal strains with divergent egg retention ($\rho_{Spearman}$ = 0.01, P = 0.98). **F** No correlation between mean body size (volume; at first fertilization, 1-2 eggs *in utero*) and mean egg retention (at L4 + 30h) across the 15 focal strains with divergent egg retention ($\rho_{Spearman}$ = 0.1, P = 0.92).

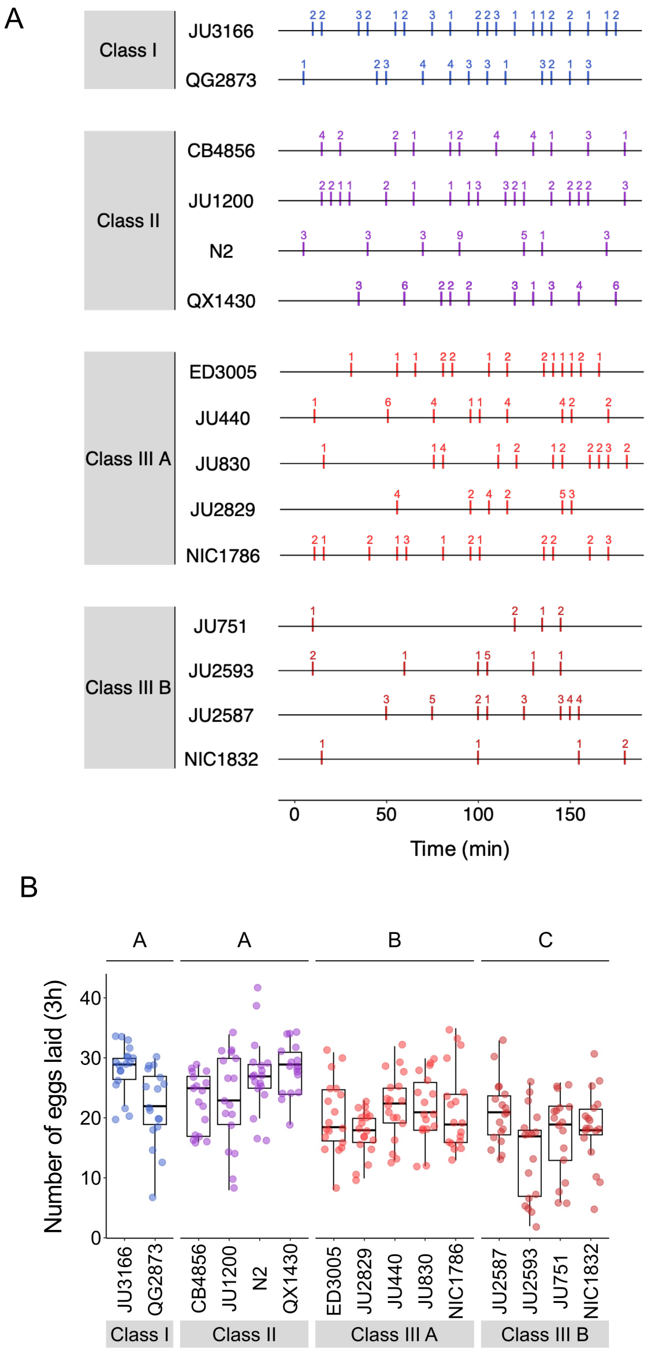

**Figure S4 Natural variation in egg-laying behaviour**
**A** Detailed representation of Fig. 3C: Representative raster plots of temporal patterns of egg-laying behaviour during a three-hour interval, with each horizontal line representing a single individual; vertical bars indicate five-minute intervals during which one or more eggs were laid. The exact number of eggs laid in a given interval is indicated above bars. **B** The total number of eggs laid during the three hours of observation differed significantly between strains and Classes. Two-Way ANOVA, fixed effect *Class:* $F_{3,248} = 23.07$, $P < 0.0001$, fixed effect *Strain(nested in Class)*: $F_{11,248} = 3.73$, $P < 0.0001$. Estimates of *Class* effects labelled with the same letter are not significantly different from each other (Tukey's honestly significant difference, $P < 0.05$). Data from same experiment as shown in Fig. 3C to 3F.

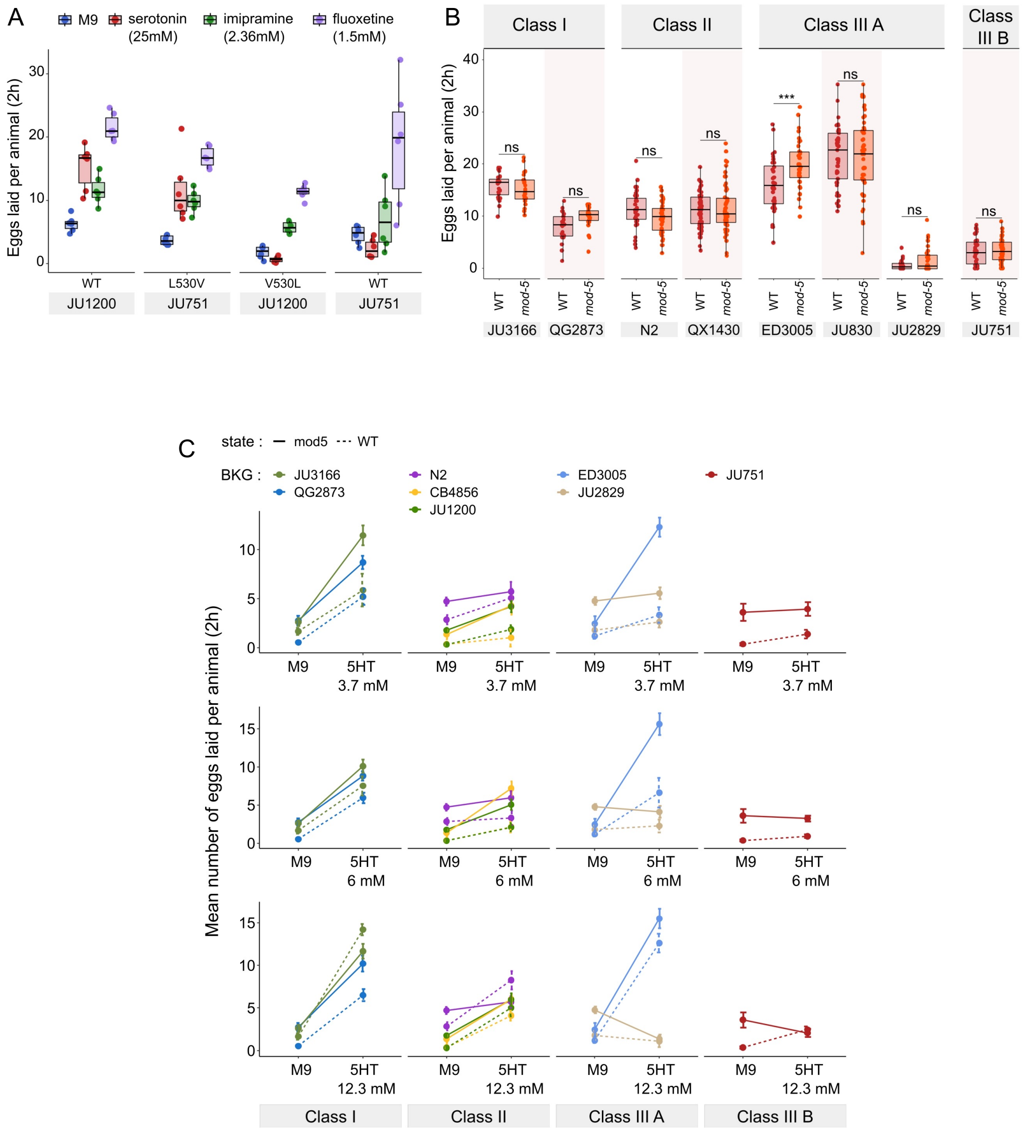

**Figure S5 Natural variation in *C. elegans* egg-laying in response to neuromodulatory inputs**
**A** Effects of exogenous serotonin, fluoxetine and imipramine on egg laying activity in strains with strongly divergent egg retention due to variation in a single amino acid residue of KCNL-1. Strains JU1200$_{WT}$ (canonical egg retention), JU1200$_{KCNL\text{-}1\ V530L}$ (CRISPR-engineered, strong egg retention), JU751$_{WT}$ (strong egg retention) and JU751$_{KCNL\text{-}1\ L530V}$ (CRISPR-engineered, low egg retention). Adult hermaphrodites (L4 + 30h) were placed into M9 buffer without food (control) or M9 with serotonin (25mM). In contrast to serotonin, imipramine and fluoxetine stimulated egg-laying activity in all four strains. For each strain, 6 replicates (each containing 5.50 ± 0.92 individuals on average) were scored for each treatment and control condition. For clarity, Fig. 4C only shows the serotonin versus control conditions of this experiment. **B** Natural variation in serotonin sensitivity of *mod-5(lf)* on egg laying (adult hermaphrodites, L4 + 30h). Align Rank Transform ANOVA, fixed effect *mod-5:* $F_{1,582}$ = 0.29, $P$ = 0.59, fixed effect *Background*: $F_{7,582}$ =277.73, $P$ < 0.0001; interaction *mod-5* x *Background*: $F_{7,582}$ = 3.57, P < 0.0001). *mod-5(lf)* did not increase egg-laying when exposed to a high dose of exogenous serotonin (25mM), except for a slight increase observed in ED3005 (Tukey's honestly significant difference, ***$P$ < 0.0001; ns: not significant). For each strain, 24-60 replicates (each containing 3.08 ± 0.14 individuals on average) were scored. For complete results of statistical analyses, see Table S6. **C** Alternative representation of data shown in Fig. 5C: Natural variation in egg laying in response to a gradient of low exogeneous serotonin concentrations in wild type and *mod-5(n822)* animals. Adult hermaphrodites (L4 + 30h) were placed into M9 buffer without food containing four different concentrations of serotonin (0.0mM, 3.7mM, 6.0mM, 12.3mM). Strains differed strongly in sensitivity to specific concentrations of exogenous serotonin and these effects of genetic background were further contingent on the presence of *mod-5(lf)* as indicated by the significant 3-way interaction term *genetic background* x *mod-5 allele* x *serotonin treatment* (Table 3). For complete results of statistical analyses, see Table 3. For each concentration, 8-10 replicates (each containing 3.15 ± 0.22 individuals on average) were scored per strain.

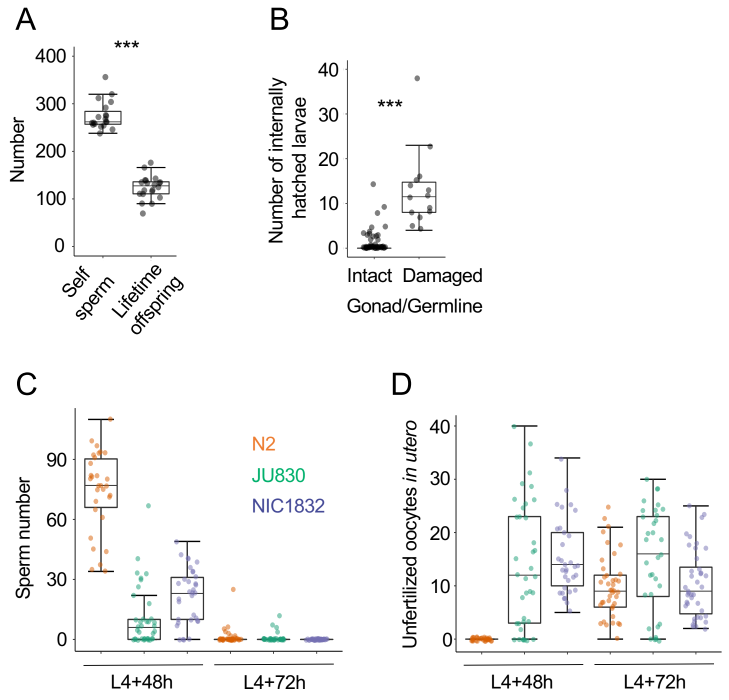

**Figure S6 Evaluating potential costs and benefits of variation in egg retention**
**A** The Class IIIB strain JU2593 shows a strong discordance between total self-sperm number (N=20 individuals) and lifetime offspring production (N=19 individuals (ANOVA, $F_{1,37} = 27.66$, $P < 0.0001$). Self-sperm was counted in DAPI-stained worms that had just reached reproductive maturity (0-5 eggs in the uterus). **B** Internal larval hatching increases the incidence of gonad/germline damage in the Class IIIB strain JU2593 (hermaphrodites at midL4 + 40h and midL4 + 48h were fixed and stained with DAPI to visualize germlines, N=105). The number of larvae *in utero* was higher in individuals with damaged gonads versus individuals with intact gonads (Kruskal-Wallis Test, $\chi^2 = 41.74$, df = 1, P < 0.0001). **C and D** Uterine accumulation of unfertilized oocytes precedes self-sperm depletion in the Class III strains JU830 and NIC1832 (but not in the Class II strain N2). For each strain, hermaphrodites (at L4 + 48h and L4 + 72h) were fixed and stained with DAPI to visualize and count remaining **(C)** self-sperm and **(D)** unfertilized oocytes in the uterus. At L4 + 48h, JU830 and NIC1832 have generated large numbers of unfertilized oocytes despite the presence of self-sperm (N2: N=30, JU830: N=37, NIC1832: N=33). At L4 + 72h, all strains have depleted their self-sperm and have unfertilized oocytes *in utero* (N2: N=40, JU830: N=34, NIC1832: N=40).

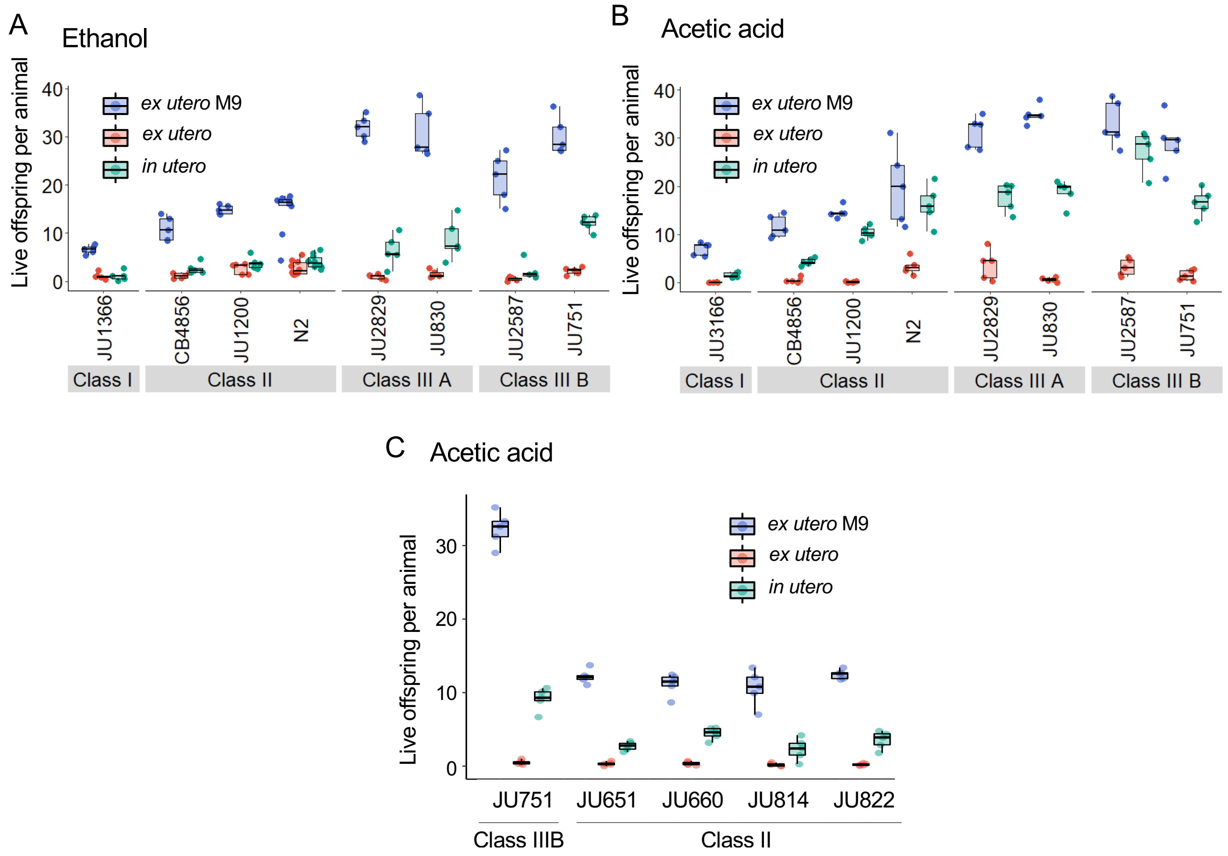

**Figure S7 Evaluating potential costs and benefits of variation in egg retention**
**A** Supplementary data for Fig. 8A: Differences in survival of eggs developing *ex utero* versus *in utero* when exposed to a high concentration of ethanol (96%, 10 minute-exposure). A subset of the 15 focal strains with divergent egg retention was selected to compare embryonic survival of eggs *ex utero* (extracted by dissection) versus *in utero* (eggs retained in mothers) exposed to ethanol. In addition to data shown in Fig. 8A, this figure includes control data for survival of eggs *ex utero* exposed to control conditions (M9 buffer). N=5-10 replicates (each containing 10 mothers) per genotype and treatment (hermaphrodites at L4 + 30h). **B** Supplementary data for Fig. 8B: Differences in survival of eggs developing *ex utero* versus *in utero* when exposed to a high concentration of acetic acid (10M, 15 minute-exposure). A subset of the 15 focal strains with divergent egg retention was selected to compare embryonic survival of eggs *ex utero* (extracted by dissection) versus *in utero* (eggs retained in mothers) exposed to acetic acid. In addition to data shown in Fig. 8B, this figure includes control data for survival of eggs *ex utero* exposed to control conditions (M9 buffer). N=5 replicates (each containing 10 mothers) per genotype and treatment (hermaphrodites at L4 + 30h). **C** Supplementary data for Fig. 8C: Differences in survival of eggs developing *ex utero* versus *in utero* when exposed to a high concentration of acetic acid (10M, 15 minute-exposure). Comparison of the strain JU751 (Class IIIB, high retention) to four strains (Class II, canonical retention) isolated from the same locality. In addition to data shown in Fig. 8C, this figure includes control data for survival of eggs *ex utero* exposed to control conditions (M9 buffer). N=5 replicates (each containing 10 mothers) per genotype and treatment (hermaphrodites at L4 + 30h).

### Supporting Tables

**Table S1.** List of strains used.

| Strain | Genotype | Source/Reference | Description |
|---|---|---|---|
| MCP355 | *mod-5(n822)*, strain N2 | This study | Single-nucleotide replacement by CRISPR-Cas9 |
| MCP374 | *mod-5(n822)*, strain ED3005 | This study | Single-nucleotide replacement by CRISPR-Cas9 |
| MCP378 | *mod-5(n822)*, strain JU2829 | This study | Single-nucleotide replacement by CRISPR-Cas9 |
| MCP376 | *mod-5(n822)*, strain QX1430 | This study | Single-nucleotide replacement by CRISPR-Cas9 |
| MCP369 | *mod-5(n822)*, strain JU830 | This study | Single-nucleotide replacement by CRISPR-Cas9 |
| MCP370 | *mod-5(n822)*, strain JU751 | This study | Single-nucleotide replacement by CRISPR-Cas9 |
| MCP387 | *mod-5(n822)*, strain CB4856 | This study | Single-nucleotide replacement by CRISPR-Cas9 |
| MCP381 | *mod-5(n822)*, strain JU1200 | This study | Single-nucleotide replacement by CRISPR-Cas9 |
| MCP383 | *mod-5(n822)*, strain JU3166 | This study | Single-nucleotide replacement by CRISPR-Cas9 |
| MCP380 | *mod-5(n822)*, strain QG2873 | This study | Single-nucleotide replacement by CRISPR-Cas9 |
| NIC1627 | *kcnl-1(cgb*1005[L530V]), strain JU751 | Vigne et al. 2021 | Single-nucleotide replacement by CRISPR-Cas9 |
| NIC1642 | *kcnl-1(cgb1*002[V530L]), strain JU1200 | Vigne et al. 2021 | Single-nucleotide replacement by CRISPR-Cas9 |
| PD4790 | mls12 (myo- 2::GFP) | *Caenorhabditis* Genetics Center (https://cgc.umn.edu/) | GFP-reporter, expression in pharynx |
| QX1430 | ttTi12715; qgIR1 | Andersen et al. 2015 | N2-derived strain with natural *npr-1 allele* (*qqR1*) |
| JU1200 | Wild strain | *CeNDR* (https://elegansvariation.org/) | |
| JU2593 | Wild strain | *CeNDR* (https://elegansvariation.org/) | |
| JU2587 | Wild strain | *CeNDR* (https://elegansvariation.org/) | |
| JU2829 | Wild strain | *CeNDR* (https://elegansvariation.org/) | |
| JU2581 | Wild strain | *CeNDR* (https://elegansvariation.org/) | |
| JU751 | Wild strain | *CeNDR* (https://elegansvariation.org/) | |
| JU3128 | Wild strain | *CeNDR* (https://elegansvariation.org/) | |
| XZ1516 | Wild strain | *CeNDR* (https://elegansvariation.org/) | |
| NIC266 | Wild strain | *CeNDR* (https://elegansvariation.org/) | |
| ECA372 | Wild strain | *CeNDR* (https://elegansvariation.org/) | |
| ECA191 | Wild strain | *CeNDR* (https://elegansvariation.org/) | |
| JU2001 | Wild strain | *CeNDR* (https://elegansvariation.org/) | |
| ECA396 | Wild strain | *CeNDR* (https://elegansvariation.org/) | |
| ED3073 | Wild strain | *CeNDR* (https://elegansvariation.org/) | |
| ECA189 | Wild strain | *CeNDR* (https://elegansvariation.org/) | |
| ECA369 | Wild strain | *CeNDR* (https://elegansvariation.org/) | |
| QX1211 | Wild strain | *CeNDR* (https://elegansvariation.org/) | |
| ECA363 | Wild strain | *CeNDR* (https://elegansvariation.org/) | |
| CX11276 | Wild strain | *CeNDR* (https://elegansvariation.org/) | |
| KR314 | Wild strain | *CeNDR* (https://elegansvariation.org/) | |
| ED3011 | Wild strain | *CeNDR* (https://elegansvariation.org/) | |
| CX11285 | Wild strain | *CeNDR* (https://elegansvariation.org/) | |
| QX1212 | Wild strain | *CeNDR* (https://elegansvariation.org/) | |
| ED3005 | Wild strain | *CeNDR* (https://elegansvariation.org/) | |
| JU830 | Wild strain | *CeNDR* (https://elegansvariation.org/) | |
| DL226 | Wild strain | *CeNDR* (https://elegansvariation.org/) | |
| CB4852 | Wild strain | *CeNDR* (https://elegansvariation.org/) | |
| JU1568 | Wild strain | *CeNDR* (https://elegansvariation.org/) | |
| MY18 | Wild strain | *CeNDR* (https://elegansvariation.org/) | |
| CB4853 | Wild strain | *CeNDR* (https://elegansvariation.org/) | |
| CB4854 | Wild strain | *CeNDR* (https://elegansvariation.org/) | |
| AB1 | Wild strain | *CeNDR* (https://elegansvariation.org/) | |
| XZ1514 | Wild strain | *CeNDR* (https://elegansvariation.org/) | |
| CB4857 | Wild strain | *CeNDR* (https://elegansvariation.org/) | |
| CB4858 | Wild strain | *CeNDR* (https://elegansvariation.org/) | |
| QX1791 | Wild strain | *CeNDR* (https://elegansvariation.org/) | |
| CX11262 | Wild strain | *CeNDR* (https://elegansvariation.org/) | |
| CX11271 | Wild strain | *CeNDR* (https://elegansvariation.org/) | |
| CB4932 | Wild strain | *CeNDR* (https://elegansvariation.org/) | |
| CX11292 | Wild strain | *CeNDR* (https://elegansvariation.org/) | |
| CX11264 | Wild strain | *CeNDR* (https://elegansvariation.org/) | |
| CX11307 | Wild strain | *CeNDR* (https://elegansvariation.org/) | |
| CX11314 | Wild strain | *CeNDR* (https://elegansvariation.org/) | |
| CX11315 | Wild strain | *CeNDR* (https://elegansvariation.org/) | |
| ED3049 | Wild strain | *CeNDR* (https://elegansvariation.org/) | |
| ED3077 | Wild strain | *CeNDR* (https://elegansvariation.org/) | |
| ED3052 | Wild strain | *CeNDR* (https://elegansvariation.org/) | |
| ED3048 | Wild strain | *CeNDR* (https://elegansvariation.org/) | |
| ED3046 | Wild strain | *CeNDR* (https://elegansvariation.org/) | |
| ED3017 | Wild strain | *CeNDR* (https://elegansvariation.org/) | |
| DL238 | Wild strain | *CeNDR* (https://elegansvariation.org/) | |
| DL200 | Wild strain | *CeNDR* (https://elegansvariation.org/) | |
| ED3012 | Wild strain | *CeNDR* (https://elegansvariation.org/) | |
| EG4724 | Wild strain | *CeNDR* (https://elegansvariation.org/) | |
| ED3040 | Wild strain | *CeNDR* (https://elegansvariation.org/) | |
| EG4347 | Wild strain | *CeNDR* (https://elegansvariation.org/) | |
| EG4725 | Wild strain | *CeNDR* (https://elegansvariation.org/) | |

| | | |
|---|---|---|
| EG4946 | Wild strain | *CeNDR* (https://elegansvariation.org/) |
| JT11398 | Wild strain | *CeNDR* (https://elegansvariation.org/) |
| JU1212 | Wild strain | *CeNDR* (https://elegansvariation.org/) |
| JU1213 | Wild strain | *CeNDR* (https://elegansvariation.org/) |
| JU1242 | Wild strain | *CeNDR* (https://elegansvariation.org/) |
| JU1246 | Wild strain | *CeNDR* (https://elegansvariation.org/) |
| JU1395 | Wild strain | *CeNDR* (https://elegansvariation.org/) |
| JU1409 | Wild strain | *CeNDR* (https://elegansvariation.org/) |
| JU1440 | Wild strain | *CeNDR* (https://elegansvariation.org/) |
| JU1491 | Wild strain | *CeNDR* (https://elegansvariation.org/) |
| JU1530 | Wild strain | *CeNDR* (https://elegansvariation.org/) |
| JU1581 | Wild strain | *CeNDR* (https://elegansvariation.org/) |
| JU1586 | Wild strain | *CeNDR* (https://elegansvariation.org/) |
| JU310 | Wild strain | *CeNDR* (https://elegansvariation.org/) |
| JU346 | Wild strain | *CeNDR* (https://elegansvariation.org/) |
| JU311 | Wild strain | *CeNDR* (https://elegansvariation.org/) |
| JU323 | Wild strain | *CeNDR* (https://elegansvariation.org/) |
| JU1400 | Wild strain | *CeNDR* (https://elegansvariation.org/) |
| JU792 | Wild strain | *CeNDR* (https://elegansvariation.org/) |
| JU397 | Wild strain | *CeNDR* (https://elegansvariation.org/) |
| JU406 | Wild strain | *CeNDR* (https://elegansvariation.org/) |
| JU440 | Wild strain | *CeNDR* (https://elegansvariation.org/) |
| JU393 | Wild strain | *CeNDR* (https://elegansvariation.org/) |
| JU394 | Wild strain | *CeNDR* (https://elegansvariation.org/) |
| MY1 | Wild strain | *CeNDR* (https://elegansvariation.org/) |
| JU774 | Wild strain | *CeNDR* (https://elegansvariation.org/) |
| LKC34 | Wild strain | *CeNDR* (https://elegansvariation.org/) |
| JU778 | Wild strain | *CeNDR* (https://elegansvariation.org/) |
| JU782 | Wild strain | *CeNDR* (https://elegansvariation.org/) |
| MY10 | Wild strain | *CeNDR* (https://elegansvariation.org/) |
| JU775 | Wild strain | *CeNDR* (https://elegansvariation.org/) |
| PB303 | Wild strain | *CeNDR* (https://elegansvariation.org/) |
| MY23 | Wild strain | *CeNDR* (https://elegansvariation.org/) |
| WN2002 | Wild strain | *CeNDR* (https://elegansvariation.org/) |
| PS2025 | Wild strain | *CeNDR* (https://elegansvariation.org/) |
| CX11254 | Wild strain | *CeNDR* (https://elegansvariation.org/) |
| RC301 | Wild strain | *CeNDR* (https://elegansvariation.org/) |
| EG4349 | Wild strain | *CeNDR* (https://elegansvariation.org/) |
| QX1233 | Wild strain | *CeNDR* (https://elegansvariation.org/) |
| JU1088 | Wild strain | *CeNDR* (https://elegansvariation.org/) |
| NIC207 | Wild strain | *CeNDR* (https://elegansvariation.org/) |
| JU1172 | Wild strain | *CeNDR* (https://elegansvariation.org/) |
| NIC166 | Wild strain | *CeNDR* (https://elegansvariation.org/) |
| JU367 | Wild strain | *CeNDR* (https://elegansvariation.org/) |
| MY16 | Wild strain | *CeNDR* (https://elegansvariation.org/) |
| CB4856 | Wild strain | *CeNDR* (https://elegansvariation.org/) |
| JU360 | Wild strain | *CeNDR* (https://elegansvariation.org/) |
| PX179 | Wild strain | *CeNDR* (https://elegansvariation.org/) |
| N2 | Wild strain | *CeNDR* (https://elegansvariation.org/) |
| BRC20067 | Wild strain | *CeNDR* (https://elegansvariation.org/) |
| NIC195 | Wild strain | *CeNDR* (https://elegansvariation.org/) |
| NIC199 | Wild strain | *CeNDR* (https://elegansvariation.org/) |
| NIC2 | Wild strain | *CeNDR* (https://elegansvariation.org/) |
| JU561 | Wild strain | *CeNDR* (https://elegansvariation.org/) |
| NIC1 | Wild strain | *CeNDR* (https://elegansvariation.org/) |
| NIC231 | Wild strain | *CeNDR* (https://elegansvariation.org/) |
| NIC236 | Wild strain | *CeNDR* (https://elegansvariation.org/) |
| NIC242 | Wild strain | *CeNDR* (https://elegansvariation.org/) |
| NIC251 | Wild strain | *CeNDR* (https://elegansvariation.org/) |
| NIC252 | Wild strain | *CeNDR* (https://elegansvariation.org/) |
| NIC255 | Wild strain | *CeNDR* (https://elegansvariation.org/) |
| NIC256 | Wild strain | *CeNDR* (https://elegansvariation.org/) |
| NIC260 | Wild strain | *CeNDR* (https://elegansvariation.org/) |
| NIC261 | Wild strain | *CeNDR* (https://elegansvariation.org/) |
| NIC262 | Wild strain | *CeNDR* (https://elegansvariation.org/) |
| NIC265 | Wild strain | *CeNDR* (https://elegansvariation.org/) |
| NIC267 | Wild strain | *CeNDR* (https://elegansvariation.org/) |
| NIC268 | Wild strain | *CeNDR* (https://elegansvariation.org/) |
| NIC269 | Wild strain | *CeNDR* (https://elegansvariation.org/) |
| NIC271 | Wild strain | *CeNDR* (https://elegansvariation.org/) |
| NIC274 | Wild strain | *CeNDR* (https://elegansvariation.org/) |
| NIC275 | Wild strain | *CeNDR* (https://elegansvariation.org/) |
| NIC276 | Wild strain | *CeNDR* (https://elegansvariation.org/) |
| NIC277 | Wild strain | *CeNDR* (https://elegansvariation.org/) |
| NIC3 | Wild strain | *CeNDR* (https://elegansvariation.org/) |
| GXW1 | Wild strain | *CeNDR* (https://elegansvariation.org/) |

| | | |
|---|---|---|
| JU2007 | Wild strain | *CeNDR* (https://elegansvariation.org/) |
| JU2316 | Wild strain | *CeNDR* (https://elegansvariation.org/) |
| NIC259 | Wild strain | *CeNDR* (https://elegansvariation.org/) |
| NIC272 | Wild strain | *CeNDR* (https://elegansvariation.org/) |
| QG556 | Wild strain | *CeNDR* (https://elegansvariation.org/) |
| QW947 | Wild strain | *CeNDR* (https://elegansvariation.org/) |
| JU2464 | Wild strain | *CeNDR* (https://elegansvariation.org/) |
| QX1794 | Wild strain | *CeNDR* (https://elegansvariation.org/) |
| JU2522 | Wild strain | *CeNDR* (https://elegansvariation.org/) |
| WN2001 | Wild strain | *CeNDR* (https://elegansvariation.org/) |
| QX1792 | Wild strain | *CeNDR* (https://elegansvariation.org/) |
| JU2513 | Wild strain | *CeNDR* (https://elegansvariation.org/) |
| QG536 | Wild strain | *CeNDR* (https://elegansvariation.org/) |
| QX1793 | Wild strain | *CeNDR* (https://elegansvariation.org/) |
| PB306 | Wild strain | *CeNDR* (https://elegansvariation.org/) |
| JU1666 | Wild strain | *CeNDR* (https://elegansvariation.org/) |
| JU3137 | Wild strain | *CeNDR* (https://elegansvariation.org/) |
| WN2033 | Wild strain | *CeNDR* (https://elegansvariation.org/) |
| MY518 | Wild strain | *CeNDR* (https://elegansvariation.org/) |
| MY2741 | Wild strain | *CeNDR* (https://elegansvariation.org/) |
| MY795 | Wild strain | *CeNDR* (https://elegansvariation.org/) |
| JU2131 | Wild strain | *CeNDR* (https://elegansvariation.org/) |
| NIC1698 | Wild strain | *CeNDR* (https://elegansvariation.org/) |
| QG2075 | Wild strain | *CeNDR* (https://elegansvariation.org/) |
| ECA36 | Wild strain | *CeNDR* (https://elegansvariation.org/) |
| NIC258 | Wild strain | *CeNDR* (https://elegansvariation.org/) |
| QG557 | Wild strain | *CeNDR* (https://elegansvariation.org/) |
| JU2466 | Wild strain | *CeNDR* (https://elegansvariation.org/) |
| JU2519 | Wild strain | *CeNDR* (https://elegansvariation.org/) |
| JU2526 | Wild strain | *CeNDR* (https://elegansvariation.org/) |
| JU2811 | Wild strain | *CeNDR* (https://elegansvariation.org/) |
| JU2879 | Wild strain | *CeNDR* (https://elegansvariation.org/) |
| CB4851 | Wild strain | *CeNDR* (https://elegansvariation.org/) |
| ECA594 | Wild strain | *CeNDR* (https://elegansvariation.org/) |
| JU3169 | Wild strain | *CeNDR* (https://elegansvariation.org/) |
| ECA744 | Wild strain | *CeNDR* (https://elegansvariation.org/) |
| ECA640 | Wild strain | *CeNDR* (https://elegansvariation.org/) |
| JU3167 | Wild strain | *CeNDR* (https://elegansvariation.org/) |
| ECA592 | Wild strain | *CeNDR* (https://elegansvariation.org/) |
| ECA593 | Wild strain | *CeNDR* (https://elegansvariation.org/) |
| ECA741 | Wild strain | *CeNDR* (https://elegansvariation.org/) |
| ECA740 | Wild strain | *CeNDR* (https://elegansvariation.org/) |
| ECA738 | Wild strain | *CeNDR* (https://elegansvariation.org/) |
| ECA724 | Wild strain | *CeNDR* (https://elegansvariation.org/) |
| ECA706 | Wild strain | *CeNDR* (https://elegansvariation.org/) |
| ECA722 | Wild strain | *CeNDR* (https://elegansvariation.org/) |
| ECA723 | Wild strain | *CeNDR* (https://elegansvariation.org/) |
| ECA347 | Wild strain | *CeNDR* (https://elegansvariation.org/) |
| JU3166 | Wild strain | *CeNDR* (https://elegansvariation.org/) |
| ECA745 | Wild strain | *CeNDR* (https://elegansvariation.org/) |
| ECA743 | Wild strain | *CeNDR* (https://elegansvariation.org/) |
| JU3226 | Wild strain | *CeNDR* (https://elegansvariation.org/) |
| ECA928 | Wild strain | *CeNDR* (https://elegansvariation.org/) |
| ECA930 | Wild strain | *CeNDR* (https://elegansvariation.org/) |
| QG2825 | Wild strain | *CeNDR* (https://elegansvariation.org/) |
| QG2828 | Wild strain | *CeNDR* (https://elegansvariation.org/) |
| QG2811 | Wild strain | *CeNDR* (https://elegansvariation.org/) |
| JU3282 | Wild strain | *CeNDR* (https://elegansvariation.org/) |
| QG2841 | Wild strain | *CeNDR* (https://elegansvariation.org/) |
| XZ1734 | Wild strain | *CeNDR* (https://elegansvariation.org/) |
| ECA746 | Wild strain | *CeNDR* (https://elegansvariation.org/) |
| ECA923 | Wild strain | *CeNDR* (https://elegansvariation.org/) |
| QG2832 | Wild strain | *CeNDR* (https://elegansvariation.org/) |
| QG2837 | Wild strain | *CeNDR* (https://elegansvariation.org/) |
| QG2846 | Wild strain | *CeNDR* (https://elegansvariation.org/) |
| ECA742 | Wild strain | *CeNDR* (https://elegansvariation.org/) |
| QG2836 | Wild strain | *CeNDR* (https://elegansvariation.org/) |
| QG2873 | Wild strain | *CeNDR* (https://elegansvariation.org/) |
| XZ1756 | Wild strain | *CeNDR* (https://elegansvariation.org/) |
| XZ1735 | Wild strain | *CeNDR* (https://elegansvariation.org/) |
| XZ2019 | Wild strain | *CeNDR* (https://elegansvariation.org/) |
| QG2823 | Wild strain | *CeNDR* (https://elegansvariation.org/) |
| QG2843 | Wild strain | *CeNDR* (https://elegansvariation.org/) |
| QG2857 | Wild strain | *CeNDR* (https://elegansvariation.org/) |
| WN2064 | Wild strain | *CeNDR* (https://elegansvariation.org/) |
| QG2855 | Wild strain | *CeNDR* (https://elegansvariation.org/) |

| | | |
|---|---|---|
| JU1896 | Wild strain | *CeNDR* (https://elegansvariation.org/) |
| JU847 | Wild strain | *CeNDR* (https://elegansvariation.org/) |
| JU258 | Wild strain | *CeNDR* (https://elegansvariation.org/) |
| ECA701 | Wild strain | *CeNDR* (https://elegansvariation.org/) |
| XZ1672 | Wild strain | *CeNDR* (https://elegansvariation.org/) |
| QG2850 | Wild strain | *CeNDR* (https://elegansvariation.org/) |
| QG2818 | Wild strain | *CeNDR* (https://elegansvariation.org/) |
| JU3224 | Wild strain | *CeNDR* (https://elegansvariation.org/) |
| QG2875 | Wild strain | *CeNDR* (https://elegansvariation.org/) |
| JU3228 | Wild strain | *CeNDR* (https://elegansvariation.org/) |
| QG2827 | Wild strain | *CeNDR* (https://elegansvariation.org/) |
| QG2854 | Wild strain | *CeNDR* (https://elegansvariation.org/) |
| ECA710 | Wild strain | *CeNDR* (https://elegansvariation.org/) |
| QG2874 | Wild strain | *CeNDR* (https://elegansvariation.org/) |
| QG2835 | Wild strain | *CeNDR* (https://elegansvariation.org/) |
| QG2838 | Wild strain | *CeNDR* (https://elegansvariation.org/) |
| QG2932 | Wild strain | *CeNDR* (https://elegansvariation.org/) |
| JU2907 | Wild strain | *CeNDR* (https://elegansvariation.org/) |
| MY2147 | Wild strain | *CeNDR* (https://elegansvariation.org/) |
| MY2585 | Wild strain | *CeNDR* (https://elegansvariation.org/) |
| JU2619 | Wild strain | *CeNDR* (https://elegansvariation.org/) |
| JU2838 | Wild strain | *CeNDR* (https://elegansvariation.org/) |
| JU2586 | Wild strain | *CeNDR* (https://elegansvariation.org/) |
| JU2825 | Wild strain | *CeNDR* (https://elegansvariation.org/) |
| JU2841 | Wild strain | *CeNDR* (https://elegansvariation.org/) |
| JU2800 | Wild strain | *CeNDR* (https://elegansvariation.org/) |
| JU1543 | Wild strain | *CeNDR* (https://elegansvariation.org/) |
| JU2862 | Wild strain | *CeNDR* (https://elegansvariation.org/) |
| JU2853 | Wild strain | *CeNDR* (https://elegansvariation.org/) |
| JU3144 | Wild strain | *CeNDR* (https://elegansvariation.org/) |
| JU2234 | Wild strain | *CeNDR* (https://elegansvariation.org/) |
| JU3132 | Wild strain | *CeNDR* (https://elegansvariation.org/) |
| JU2610 | Wild strain | *CeNDR* (https://elegansvariation.org/) |
| MY2453 | Wild strain | *CeNDR* (https://elegansvariation.org/) |
| NIC1107 | Wild strain | *CeNDR* (https://elegansvariation.org/) |
| MY2573 | Wild strain | *CeNDR* (https://elegansvariation.org/) |
| MY679 | Wild strain | *CeNDR* (https://elegansvariation.org/) |
| NIC513 | Wild strain | *CeNDR* (https://elegansvariation.org/) |
| XZ1513 | Wild strain | *CeNDR* (https://elegansvariation.org/) |
| NIC522 | Wild strain | *CeNDR* (https://elegansvariation.org/) |
| NIC523 | Wild strain | *CeNDR* (https://elegansvariation.org/) |
| MY920 | Wild strain | *CeNDR* (https://elegansvariation.org/) |
| NIC514 | Wild strain | *CeNDR* (https://elegansvariation.org/) |
| MY2693 | Wild strain | *CeNDR* (https://elegansvariation.org/) |
| MY2530 | Wild strain | *CeNDR* (https://elegansvariation.org/) |
| MY2212 | Wild strain | *CeNDR* (https://elegansvariation.org/) |
| MY2535 | Wild strain | *CeNDR* (https://elegansvariation.org/) |
| JU3140 | Wild strain | *CeNDR* (https://elegansvariation.org/) |
| JU3135 | Wild strain | *CeNDR* (https://elegansvariation.org/) |
| JU1249 | Wild strain | *CeNDR* (https://elegansvariation.org/) |
| JU1793 | Wild strain | *CeNDR* (https://elegansvariation.org/) |
| JU2878 | Wild strain | *CeNDR* (https://elegansvariation.org/) |
| JU2534 | Wild strain | *CeNDR* (https://elegansvariation.org/) |
| JU2016 | Wild strain | *CeNDR* (https://elegansvariation.org/) |
| JU2141 | Wild strain | *CeNDR* (https://elegansvariation.org/) |
| BRC20263 | Wild strain | *CeNDR* (https://elegansvariation.org/) |
| JU1792 | Wild strain | *CeNDR* (https://elegansvariation.org/) |
| JU2600 | Wild strain | *CeNDR* (https://elegansvariation.org/) |
| JU2565 | Wild strain | *CeNDR* (https://elegansvariation.org/) |
| JU1934 | Wild strain | *CeNDR* (https://elegansvariation.org/) |
| JU2906 | Wild strain | *CeNDR* (https://elegansvariation.org/) |
| ECA348 | Wild strain | *CeNDR* (https://elegansvariation.org/) |
| JU1808 | Wild strain | *CeNDR* (https://elegansvariation.org/) |
| NIC515 | Wild strain | *CeNDR* (https://elegansvariation.org/) |
| JU2257 | Wild strain | *CeNDR* (https://elegansvariation.org/) |
| JU2250 | Wild strain | *CeNDR* (https://elegansvariation.org/) |
| JU2478 | Wild strain | *CeNDR* (https://elegansvariation.org/) |
| JU2017 | Wild strain | *CeNDR* (https://elegansvariation.org/) |
| NIC511 | Wild strain | *CeNDR* (https://elegansvariation.org/) |
| JU2576 | Wild strain | *CeNDR* (https://elegansvariation.org/) |
| MY2713 | Wild strain | *CeNDR* (https://elegansvariation.org/) |
| JU2572 | Wild strain | *CeNDR* (https://elegansvariation.org/) |
| JU2575 | Wild strain | *CeNDR* (https://elegansvariation.org/) |
| XZ1515 | Wild strain | *CeNDR* (https://elegansvariation.org/) |
| MY772 | Wild strain | *CeNDR* (https://elegansvariation.org/) |
| JU2570 | Wild strain | *CeNDR* (https://elegansvariation.org/) |

| | | |
|---|---|---|
| NIC1049 | Wild strain | *CeNDR* (https://elegansvariation.org/) |
| JU2106 | Wild strain | *CeNDR* (https://elegansvariation.org/) |
| ECA349 | Wild strain | *CeNDR* (https://elegansvariation.org/) |
| JU2592 | Wild strain | *CeNDR* (https://elegansvariation.org/) |
| WN2063 | Wild strain | *CeNDR* (https://elegansvariation.org/) |
| WN2050 | Wild strain | *CeNDR* (https://elegansvariation.org/) |
| WN2066 | Wild strain | *CeNDR* (https://elegansvariation.org/) |
| NIC501 | Wild strain | *CeNDR* (https://elegansvariation.org/) |
| NIC1785 | Wild strain | *CeNDR* (https://elegansvariation.org/) |
| NIC1789 | Wild strain | *CeNDR* (https://elegansvariation.org/) |
| NIC1786 | Wild strain | *CeNDR* (https://elegansvariation.org/) |
| NIC1780 | Wild strain | *CeNDR* (https://elegansvariation.org/) |
| NIC1782 | Wild strain | *CeNDR* (https://elegansvariation.org/) |
| NIC1790 | Wild strain | *CeNDR* (https://elegansvariation.org/) |
| NIC1791 | Wild strain | *CeNDR* (https://elegansvariation.org/) |
| NIC1783 | Wild strain | *CeNDR* (https://elegansvariation.org/) |
| NIC1779 | Wild strain | *CeNDR* (https://elegansvariation.org/) |
| NIC1788 | Wild strain | *CeNDR* (https://elegansvariation.org/) |
| NIC1787 | Wild strain | *CeNDR* (https://elegansvariation.org/) |
| NIC1792 | Wild strain | *CeNDR* (https://elegansvariation.org/) |
| NIC1802 | Wild strain | *CeNDR* (https://elegansvariation.org/) |
| NIC1796 | Wild strain | *CeNDR* (https://elegansvariation.org/) |
| NIC1798 | Wild strain | *CeNDR* (https://elegansvariation.org/) |
| NIC1794 | Wild strain | *CeNDR* (https://elegansvariation.org/) |
| NIC1799 | Wild strain | *CeNDR* (https://elegansvariation.org/) |
| NIC1801 | Wild strain | *CeNDR* (https://elegansvariation.org/) |
| NIC1805 | Wild strain | *CeNDR* (https://elegansvariation.org/) |
| NIC1812 | Wild strain | *CeNDR* (https://elegansvariation.org/) |
| NIC1808 | Wild strain | *CeNDR* (https://elegansvariation.org/) |
| NIC1832 | Wild strain | *CeNDR* (https://elegansvariation.org/) |
| NIC1809 | Wild strain | *CeNDR* (https://elegansvariation.org/) |

**Table S2 (accompanies Fig. 4B).** Results for statistical analyses testing for the effects of and interactions between *Strain* and *Treatment* (control versus serotonin) on egg laying (Align Rank Transform ANOVA).

| Source | Df | Sum of squares | F value | Pr(>F) | Sig |
|---|---|---|---|---|---|
| Treatment | 1 | 3136965,42 | 432,62 | 3,45e-65 | *** |
| Strain | 14 | 3719884,79 | 42,94 | 4,44e-70 | *** |
| Treatment x Strain | 14 | 3373489,93 | 34,40 | 1,95e-59 | *** |

**Contrasts**

| contrast | estimate | SE | Df | t-ratio | p-value |
|---|---|---|---|---|---|
| M9,CB4856 - M9,ED3005 | -25,63 | 24,69 | 390 | -1,04 | 1,00E+00 |
| M9,CB4856 - M9,JU1200 | -61,08 | 22,80 | 390 | -2,68 | 6,46E-01 |
| M9,CB4856 - M9,JU2587 | -117,54 | 24,69 | 390 | -4,76 | 1,05E-03 |
| M9,CB4856 - M9,JU2593 | 37,38 | 24,69 | 390 | 1,51 | 1,00E+00 |
| M9,CB4856 - M9,JU2829 | -171,88 | 24,69 | 390 | -6,96 | 6,23E-09 |
| M9,CB4856 - M9,JU3166 | -172,33 | 24,69 | 390 | -6,98 | 5,54E-09 |
| M9,CB4856 - M9,JU440 | -90,67 | 24,69 | 390 | -3,67 | 7,02E-02 |
| M9,CB4856 - M9,JU751 | -106,29 | 24,69 | 390 | -4,30 | 7,27E-03 |
| M9,CB4856 - M9,JU830 | -97,58 | 24,69 | 390 | -3,95 | 2,75E-02 |
| M9,CB4856 - M9,N2 | -115,10 | 21,38 | 390 | -5,38 | 5,25E-05 |
| M9,CB4856 - M9,NIC1786 | 34,61 | 25,25 | 390 | 1,37 | 1,00E+00 |
| M9,CB4856 - M9,NIC1832 | -38,08 | 24,69 | 390 | -1,54 | 1,00E+00 |
| M9,CB4856 - M9,QG2873 | -96,00 | 24,69 | 390 | -3,89 | 3,44E-02 |
| M9,CB4856 - M9,QX1430 | -134,58 | 21,38 | 390 | -6,29 | 3,59E-07 |
| M9,CB4856 - serotonin,CB4856 | -228,08 | 24,69 | 390 | -9,24 | <0,0001 |
| M9,CB4856 - serotonin,ED3005 | -323,21 | 24,69 | 390 | -13,09 | <0,0001 |
| M9,CB4856 - serotonin,JU1200 | -242,08 | 22,54 | 390 | -10,74 | <0,0001 |
| M9,CB4856 - serotonin,JU2587 | -29,87 | 24,69 | 390 | -1,21 | 1,00E+00 |
| M9,CB4856 - serotonin,JU2593 | -18,40 | 23,42 | 390 | -0,79 | 1,00E+00 |
| M9,CB4856 - serotonin,JU2829 | 11,83 | 24,69 | 390 | 0,48 | 1,00E+00 |
| M9,CB4856 - serotonin,JU3166 | -311,71 | 24,69 | 390 | -12,62 | <0,0001 |
| M9,CB4856 - serotonin,JU440 | -287,92 | 24,69 | 390 | -11,66 | <0,0001 |
| M9,CB4856 - serotonin,JU751 | -27,79 | 24,69 | 390 | -1,13 | 1,00E+00 |
| M9,CB4856 - serotonin,JU830 | -287,25 | 24,69 | 390 | -11,63 | <0,0001 |
| M9,CB4856 - serotonin,N2 | -269,44 | 21,38 | 390 | -12,60 | <0,0001 |
| M9,CB4856 - serotonin,NIC1786 | -152,08 | 24,69 | 390 | -6,16 | 7,80E-07 |
| M9,CB4856 - serotonin,NIC1832 | -5,71 | 25,25 | 390 | -0,23 | 1,00E+00 |
| M9,CB4856 - serotonin,QG2873 | -245,67 | 24,69 | 390 | -9,95 | <0,0001 |
| M9,CB4856 - serotonin,QX1430 | -225,52 | 21,38 | 390 | -10,55 | <0,0001 |
| M9,ED3005 - M9,JU1200 | -35,45 | 22,80 | 390 | -1,55 | 1,00E+00 |
| M9,ED3005 - M9,JU2587 | -91,92 | 24,69 | 390 | -3,72 | 5,97E-02 |
| M9,ED3005 - M9,JU2593 | 63,00 | 24,69 | 390 | 2,55 | 7,42E-01 |
| M9,ED3005 - M9,JU2829 | -146,25 | 24,69 | 390 | -5,92 | 2,96E-06 |
| M9,ED3005 - M9,JU3166 | -146,71 | 24,69 | 390 | -5,94 | 2,67E-06 |
| M9,ED3005 - M9,JU440 | -65,04 | 24,69 | 390 | -2,63 | 6,80E-01 |
| M9,ED3005 - M9,JU751 | -80,67 | 24,69 | 390 | -3,27 | 2,18E-01 |
| M9,ED3005 - M9,JU830 | -71,96 | 24,69 | 390 | -2,91 | 4,55E-01 |
| M9,ED3005 - M9,N2 | -89,48 | 21,38 | 390 | -4,18 | 1,16E-02 |
| M9,ED3005 - M9,NIC1786 | 60,23 | 25,25 | 390 | 2,39 | 8,49E-01 |

| | | | | | |
|---|---|---|---|---|---|
| M9,ED3005 - M9,NIC1832 | -12,46 | 24,69 | 390 | -0,50 | 1,00E+00 |
| M9,ED3005 - M9,QG2873 | -70,38 | 24,69 | 390 | -2,85 | 5,06E-01 |
| M9,ED3005 - M9,QX1430 | -108,96 | 21,38 | 390 | -5,10 | 2,19E-04 |
| M9,ED3005 - serotonin,CB4856 | -202,46 | 24,69 | 390 | -8,20 | <0,0001 |
| M9,ED3005 - serotonin,ED3005 | -297,58 | 24,69 | 390 | -12,05 | <0,0001 |
| M9,ED3005 - serotonin,JU1200 | -216,46 | 22,54 | 390 | -9,60 | <0,0001 |
| M9,ED3005 - serotonin,JU2587 | -4,25 | 24,69 | 390 | -0,17 | 1,00E+00 |
| M9,ED3005 - serotonin,JU2593 | 7,23 | 23,42 | 390 | 0,31 | 1,00E+00 |
| M9,ED3005 - serotonin,JU2829 | 37,46 | 24,69 | 390 | 1,52 | 1,00E+00 |
| M9,ED3005 - serotonin,JU3166 | -286,08 | 24,69 | 390 | -11,59 | <0,0001 |
| M9,ED3005 - serotonin,JU440 | -262,29 | 24,69 | 390 | -10,62 | <0,0001 |
| M9,ED3005 - serotonin,JU751 | -2,17 | 24,69 | 390 | -0,09 | 1,00E+00 |
| M9,ED3005 - serotonin,JU830 | -261,62 | 24,69 | 390 | -10,60 | <0,0001 |
| M9,ED3005 - serotonin,N2 | -243,81 | 21,38 | 390 | -11,40 | <0,0001 |
| M9,ED3005 - serotonin,NIC1786 | -126,46 | 24,69 | 390 | -5,12 | 1,93E-04 |
| M9,ED3005 - serotonin,NIC1832 | 19,91 | 25,25 | 390 | 0,79 | 1,00E+00 |
| M9,ED3005 - serotonin,QG2873 | -220,04 | 24,69 | 390 | -8,91 | <0,0001 |
| M9,ED3005 - SEROTONIN,QX1430 | -199,90 | 21,38 | 390 | -9,35 | <0,0001 |
| M9,JU1200 - M9,JU2587 | -56,46 | 22,80 | 390 | -2,48 | 7,94E-01 |
| M9,JU1200 - M9,JU2593 | 98,45 | 22,80 | 390 | 4,32 | 6,91E-03 |
| M9,JU1200 - M9,JU2829 | -110,80 | 22,80 | 390 | -4,86 | 6,68E-04 |
| M9,JU1200 - M9,JU3166 | -111,25 | 22,80 | 390 | -4,88 | 6,09E-04 |
| M9,JU1200 - M9,JU440 | -29,59 | 22,80 | 390 | -1,30 | 1,00E+00 |
| M9,JU1200 - M9,JU751 | -45,21 | 22,80 | 390 | -1,98 | 9,80E-01 |
| M9,JU1200 - M9,JU830 | -36,50 | 22,80 | 390 | -1,60 | 9,99E-01 |
| M9,JU1200 - M9,N2 | -54,03 | 19,17 | 390 | -2,82 | 5,33E-01 |
| M9,JU1200 - M9,NIC1786 | 95,68 | 23,40 | 390 | 4,09 | 1,68E-02 |
| M9,JU1200 - M9,NIC1832 | 23,00 | 22,80 | 390 | 1,01 | 1,00E+00 |
| M9,JU1200 - M9,QG2873 | -34,92 | 22,80 | 390 | -1,53 | 1,00E+00 |
| M9,JU1200 - M9,QX1430 | -73,50 | 19,17 | 390 | -3,83 | 4,14E-02 |
| M9,JU1200 - serotonin,CB4856 | -167,00 | 22,80 | 390 | -7,32 | 5,91E-10 |
| M9,JU1200 - serotonin,ED3005 | -262,13 | 22,80 | 390 | -11,49 | <0,0001 |
| M9,JU1200 - serotonin,JU1200 | -181,00 | 20,46 | 390 | -8,85 | <0,0001 |
| M9,JU1200 - serotonin,JU2587 | 31,20 | 22,80 | 390 | 1,37 | 1,00E+00 |
| M9,JU1200 - serotonin,JU2593 | 42,68 | 21,43 | 390 | 1,99 | 9,79E-01 |
| M9,JU1200 - serotonin,JU2829 | 72,91 | 22,80 | 390 | 3,20 | 2,57E-01 |
| M9,JU1200 - serotonin,JU3166 | -250,63 | 22,80 | 390 | -10,99 | <0,0001 |
| M9,JU1200 - serotonin,JU440 | -226,84 | 22,80 | 390 | -9,95 | <0,0001 |
| M9,JU1200 - serotonin,JU751 | 33,29 | 22,80 | 390 | 1,46 | 1,00E+00 |
| M9,JU1200 - serotonin,JU830 | -226,17 | 22,80 | 390 | -9,92 | <0,0001 |
| M9,JU1200 - serotonin,N2 | -208,36 | 19,17 | 390 | -10,87 | <0,0001 |
| M9,JU1200 - serotonin,NIC1786 | -91,00 | 22,80 | 390 | -3,99 | 2,40E-02 |
| M9,JU1200 - serotonin,NIC1832 | 55,37 | 23,40 | 390 | 2,37 | 8,59E-01 |
| M9,JU1200 - serotonin,QG2873 | -184,59 | 22,80 | 390 | -8,09 | <0,0001 |
| M9,JU1200 - serotonin,QX1430 | -164,44 | 19,17 | 390 | -8,58 | <0,0001 |
| M9,JU2587 - M9,JU2593 | 154,92 | 24,69 | 390 | 6,27 | 4,02E-07 |
| M9,JU2587 - M9,JU2829 | -54,33 | 24,69 | 390 | -2,20 | 9,31E-01 |
| M9,JU2587 - M9,JU3166 | -54,79 | 24,69 | 390 | -2,22 | 9,24E-01 |
| M9,JU2587 - M9,JU440 | 26,88 | 24,69 | 390 | 1,09 | 1,00E+00 |
| M9,JU2587 - M9,JU751 | 11,25 | 24,69 | 390 | 0,46 | 1,00E+00 |

| | | | | | |
|---|---|---|---|---|---|
| M9,JU2587 - M9,JU830 | 19,96 | 24,69 | 390 | 0,81 | 1,00E+00 |
| M9,JU2587 - M9,N2 | 2,44 | 21,38 | 390 | 0,11 | 1,00E+00 |
| M9,JU2587 - M9,NIC1786 | 152,15 | 25,25 | 390 | 6,03 | 1,66E-06 |
| M9,JU2587 - M9,NIC1832 | 79,46 | 24,69 | 390 | 3,22 | 2,45E-01 |
| M9,JU2587 - M9,QG2873 | 21,54 | 24,69 | 390 | 0,87 | 1,00E+00 |
| M9,JU2587 - M9,QX1430 | -17,04 | 21,38 | 390 | -0,80 | 1,00E+00 |
| M9,JU2587 - serotonin,CB4856 | -110,54 | 24,69 | 390 | -4,48 | 3,59E-03 |
| M9,JU2587 - serotonin,ED3005 | -205,67 | 24,69 | 390 | -8,33 | <0,0001 |
| M9,JU2587 - serotonin,JU1200 | -124,54 | 22,54 | 390 | -5,53 | 2,52E-05 |
| M9,JU2587 - serotonin,JU2587 | 87,67 | 24,69 | 390 | 3,55 | 1,01E-01 |
| M9,JU2587 - serotonin,JU2593 | 99,14 | 23,42 | 390 | 4,23 | 9,67E-03 |
| M9,JU2587 - serotonin,JU2829 | 129,38 | 24,69 | 390 | 5,24 | 1,08E-04 |
| M9,JU2587 - serotonin,JU3166 | -194,17 | 24,69 | 390 | -7,86 | 4,24E-13 |
| M9,JU2587 - serotonin,JU440 | -170,37 | 24,69 | 390 | -6,90 | 9,13E-09 |
| M9,JU2587 - serotonin,JU751 | 89,75 | 24,69 | 390 | 3,63 | 7,87E-02 |
| M9,JU2587 - serotonin,JU830 | -169,71 | 24,69 | 390 | -6,87 | 1,08E-08 |
| M9,JU2587 - serotonin,N2 | -151,90 | 21,38 | 390 | -7,10 | 2,51E-09 |
| M9,JU2587 - serotonin,NIC1786 | -34,54 | 24,69 | 390 | -1,40 | 1,00E+00 |
| M9,JU2587 - serotonin,NIC1832 | 111,83 | 25,25 | 390 | 4,43 | 4,37E-03 |
| M9,JU2587 - serotonin,QG2873 | -128,12 | 24,69 | 390 | -5,19 | 1,39E-04 |
| M9,JU2587 - serotonin,QX1430 | -107,98 | 21,38 | 390 | -5,05 | 2,72E-04 |
| M9,JU2593 - M9,JU2829 | -209,25 | 24,69 | 390 | -8,47 | <0,0001 |
| M9,JU2593 - M9,JU3166 | -209,71 | 24,69 | 390 | -8,49 | <0,0001 |
| M9,JU2593 - M9,JU440 | -128,04 | 24,69 | 390 | -5,19 | 1,41E-04 |
| M9,JU2593 - M9,JU751 | -143,67 | 24,69 | 390 | -5,82 | 5,26E-06 |
| M9,JU2593 - M9,JU830 | -134,96 | 24,69 | 390 | -5,47 | 3,43E-05 |
| M9,JU2593 - M9,N2 | -152,48 | 21,38 | 390 | -7,13 | 2,10E-09 |
| M9,JU2593 - M9,NIC1786 | -2,77 | 25,25 | 390 | -0,11 | 1,00E+00 |
| M9,JU2593 - M9,NIC1832 | -75,46 | 24,69 | 390 | -3,06 | 3,49E-01 |
| M9,JU2593 - M9,QG2873 | -133,38 | 24,69 | 390 | -5,40 | 4,77E-05 |
| M9,JU2593 - M9,QX1430 | -171,96 | 21,38 | 390 | -8,04 | <0,0001 |
| M9,JU2593 - serotonin,CB4856 | -265,46 | 24,69 | 390 | -10,75 | <0,0001 |
| M9,JU2593 - serotonin,ED3005 | -360,58 | 24,69 | 390 | -14,60 | <0,0001 |
| M9,JU2593 - serotonin,JU1200 | -279,46 | 22,54 | 390 | -12,40 | <0,0001 |
| M9,JU2593 - serotonin,JU2587 | -67,25 | 24,69 | 390 | -2,72 | 6,09E-01 |
| M9,JU2593 - serotonin,JU2593 | -55,78 | 23,42 | 390 | -2,38 | 8,51E-01 |
| M9,JU2593 - serotonin,JU2829 | -25,54 | 24,69 | 390 | -1,03 | 1,00E+00 |
| M9,JU2593 - serotonin,JU3166 | -349,08 | 24,69 | 390 | -14,14 | <0,0001 |
| M9,JU2593 - serotonin,JU440 | -325,29 | 24,69 | 390 | -13,17 | <0,0001 |
| M9,JU2593 - serotonin,JU751 | -65,17 | 24,69 | 390 | -2,64 | 6,77E-01 |
| M9,JU2593 - serotonin,JU830 | -324,63 | 24,69 | 390 | -13,15 | <0,0001 |
| M9,JU2593 - serotonin,N2 | -306,81 | 21,38 | 390 | -14,35 | <0,0001 |
| M9,JU2593 - serotonin,NIC1786 | -189,46 | 24,69 | 390 | -7,67 | 4,34E-11 |
| M9,JU2593 - serotonin,NIC1832 | -43,09 | 25,25 | 390 | -1,71 | 9,98E-01 |
| M9,JU2593 - serotonin,QG2873 | -283,04 | 24,69 | 390 | -11,46 | <0,0001 |
| M9,JU2593 - serotonin,QX1430 | -262,90 | 21,38 | 390 | -12,29 | <0,0001 |
| M9,JU2829 - M9,JU3166 | -0,46 | 24,69 | 390 | -0,02 | 1,00E+00 |
| M9,JU2829 - M9,JU440 | 81,21 | 24,69 | 390 | 3,29 | 2,06E-01 |
| M9,JU2829 - M9,JU751 | 65,58 | 24,69 | 390 | 2,66 | 6,63E-01 |
| M9,JU2829 - M9,JU830 | 74,29 | 24,69 | 390 | 3,01 | 3,83E-01 |

| | | | | | |
|---|---|---|---|---|---|
| M9,JU2829 - M9,N2 | 56,77 | 21,38 | 390 | 2,65 | 6,64E-01 |
| M9,JU2829 - M9,NIC1786 | 206,48 | 25,25 | 390 | 8,18 | <0,0001 |
| M9,JU2829 - M9,NIC1832 | 133,79 | 24,69 | 390 | 5,42 | 4,37E-05 |
| M9,JU2829 - M9,QG2873 | 75,88 | 24,69 | 390 | 3,07 | 3,37E-01 |
| M9,JU2829 - M9,QX1430 | 37,29 | 21,38 | 390 | 1,74 | 9,97E-01 |
| M9,JU2829 - serotonin,CB4856 | -56,21 | 24,69 | 390 | -2,28 | 9,02E-01 |
| M9,JU2829 - serotonin,ED3005 | -151,33 | 24,69 | 390 | -6,13 | 9,29E-07 |
| M9,JU2829 - serotonin,JU1200 | -70,21 | 22,54 | 390 | -3,11 | 3,08E-01 |
| M9,JU2829 - serotonin,JU2587 | 142,00 | 24,69 | 390 | 5,75 | 7,59E-06 |
| M9,JU2829 - serotonin,JU2593 | 153,48 | 23,42 | 390 | 6,55 | 7,77E-08 |
| M9,JU2829 - serotonin,JU2829 | 183,71 | 24,69 | 390 | 7,44 | 2,66E-10 |
| M9,JU2829 - serotonin,JU3166 | -139,83 | 24,69 | 390 | -5,66 | 1,22E-05 |
| M9,JU2829 - serotonin,JU440 | -116,04 | 24,69 | 390 | -4,70 | 1,37E-03 |
| M9,JU2829 - serotonin,JU751 | 144,08 | 24,69 | 390 | 5,84 | 4,80E-06 |
| M9,JU2829 - serotonin,JU830 | -115,37 | 24,69 | 390 | -4,67 | 1,55E-03 |
| M9,JU2829 - serotonin,N2 | -97,56 | 21,38 | 390 | -4,56 | 2,50E-03 |
| M9,JU2829 - serotonin,NIC1786 | 19,79 | 24,69 | 390 | 0,80 | 1,00E+00 |
| M9,JU2829 - serotonin,NIC1832 | 166,16 | 25,25 | 390 | 6,58 | 6,49E-08 |
| M9,JU2829 - serotonin,QG2873 | -73,79 | 24,69 | 390 | -2,99 | 3,98E-01 |
| M9,JU2829 - serotonin,QX1430 | -53,65 | 21,38 | 390 | -2,51 | 7,72E-01 |
| M9,JU3166 - M9,JU440 | 81,67 | 24,69 | 390 | 3,31 | 1,97E-01 |
| M9,JU3166 - M9,JU751 | 66,04 | 24,69 | 390 | 2,67 | 6,49E-01 |
| M9,JU3166 - M9,JU830 | 74,75 | 24,69 | 390 | 3,03 | 3,69E-01 |
| M9,JU3166 - M9,N2 | 57,23 | 21,38 | 390 | 2,68 | 6,47E-01 |
| M9,JU3166 - M9,NIC1786 | 206,94 | 25,25 | 390 | 8,20 | <0,0001 |
| M9,JU3166 - M9,NIC1832 | 134,25 | 24,69 | 390 | 5,44 | 3,98E-05 |
| M9,JU3166 - M9,QG2873 | 76,33 | 24,69 | 390 | 3,09 | 3,24E-01 |
| M9,JU3166 - M9,QX1430 | 37,75 | 21,38 | 390 | 1,77 | 9,96E-01 |
| M9,JU3166 - serotonin,CB4856 | -55,75 | 24,69 | 390 | -2,26 | 9,10E-01 |
| M9,JU3166 - serotonin,ED3005 | -150,88 | 24,69 | 390 | -6,11 | 1,03E-06 |
| M9,JU3166 - serotonin,JU1200 | -69,75 | 22,54 | 390 | -3,09 | 3,22E-01 |
| M9,JU3166 - serotonin,JU2587 | 142,46 | 24,69 | 390 | 5,77 | 6,86E-06 |
| M9,JU3166 - serotonin,JU2593 | 153,93 | 23,42 | 390 | 6,57 | 6,91E-08 |
| M9,JU3166 - serotonin,JU2829 | 184,17 | 24,69 | 390 | 7,46 | 2,33E-10 |
| M9,JU3166 - serotonin,JU3166 | -139,37 | 24,69 | 390 | -5,64 | 1,34E-05 |
| M9,JU3166 - serotonin,JU440 | -115,58 | 24,69 | 390 | -4,68 | 1,49E-03 |
| M9,JU3166 - serotonin,JU751 | 144,54 | 24,69 | 390 | 5,85 | 4,33E-06 |
| M9,JU3166 - serotonin,JU830 | -114,92 | 24,69 | 390 | -4,65 | 1,68E-03 |
| M9,JU3166 - serotonin,N2 | -97,10 | 21,38 | 390 | -4,54 | 2,74E-03 |
| M9,JU3166 - serotonin,NIC1786 | 20,25 | 24,69 | 390 | 0,82 | 1,00E+00 |
| M9,JU3166 - serotonin,NIC1832 | 166,62 | 25,25 | 390 | 6,60 | 5,82E-08 |
| M9,JU3166 - serotonin,QG2873 | -73,33 | 24,69 | 390 | -2,97 | 4,12E-01 |
| M9,JU3166 - serotonin,QX1430 | -53,19 | 21,38 | 390 | -2,49 | 7,87E-01 |
| M9,JU440 - M9,JU751 | -15,63 | 24,69 | 390 | -0,63 | 1,00E+00 |
| M9,JU440 - M9,JU830 | -6,92 | 24,69 | 390 | -0,28 | 1,00E+00 |
| M9,JU440 - M9,N2 | -24,44 | 21,38 | 390 | -1,14 | 1,00E+00 |
| M9,JU440 - M9,NIC1786 | 125,27 | 25,25 | 390 | 4,96 | 4,13E-04 |
| M9,JU440 - M9,NIC1832 | 52,58 | 24,69 | 390 | 2,13 | 9,52E-01 |
| M9,JU440 - M9,QG2873 | -5,33 | 24,69 | 390 | -0,22 | 1,00E+00 |
| M9,JU440 - M9,QX1430 | -43,92 | 21,38 | 390 | -2,05 | 9,68E-01 |

| | | | | | |
|---|---|---|---|---|---|
| M9,JU440 - serotonin,CB4856 | -137,42 | 24,69 | 390 | -5,57 | 2,04E-05 |
| M9,JU440 - serotonin,ED3005 | -232,54 | 24,69 | 390 | -9,42 | <0,0001 |
| M9,JU440 - serotonin,JU1200 | -151,42 | 22,54 | 390 | -6,72 | 2,84E-08 |
| M9,JU440 - serotonin,JU2587 | 60,79 | 24,69 | 390 | 2,46 | 8,03E-01 |
| M9,JU440 - serotonin,JU2593 | 72,27 | 23,42 | 390 | 3,09 | 3,28E-01 |
| M9,JU440 - serotonin,JU2829 | 102,50 | 24,69 | 390 | 4,15 | 1,32E-02 |
| M9,JU440 - serotonin,JU3166 | -221,04 | 24,69 | 390 | -8,95 | <0,0001 |
| M9,JU440 - serotonin,JU440 | -197,25 | 24,69 | 390 | -7,99 | <0,0001 |
| M9,JU440 - serotonin,JU751 | 62,88 | 24,69 | 390 | 2,55 | 7,46E-01 |
| M9,JU440 - serotonin,JU830 | -196,58 | 24,69 | 390 | -7,96 | <0,0001 |
| M9,JU440 - serotonin,N2 | -178,77 | 21,38 | 390 | -8,36 | <0,0001 |
| M9,JU440 - serotonin,NIC1786 | -61,42 | 24,69 | 390 | -2,49 | 7,87E-01 |
| M9,JU440 - serotonin,NIC1832 | 84,95 | 25,25 | 390 | 3,36 | 1,70E-01 |
| M9,JU440 - serotonin,QG2873 | -155,00 | 24,69 | 390 | -6,28 | 3,95E-07 |
| M9,JU440 - serotonin,QX1430 | -134,85 | 21,38 | 390 | -6,31 | 3,33E-07 |
| M9,JU751 - M9,JU830 | 8,71 | 24,69 | 390 | 0,35 | 1,00E+00 |
| M9,JU751 - M9,N2 | -8,81 | 21,38 | 390 | -0,41 | 1,00E+00 |
| M9,JU751 - M9,NIC1786 | 140,90 | 25,25 | 390 | 5,58 | 1,88E-05 |
| M9,JU751 - M9,NIC1832 | 68,21 | 24,69 | 390 | 2,76 | 5,78E-01 |
| M9,JU751 - M9,QG2873 | 10,29 | 24,69 | 390 | 0,42 | 1,00E+00 |
| M9,JU751 - M9,QX1430 | -28,29 | 21,38 | 390 | -1,32 | 1,00E+00 |
| M9,JU751 - serotonin,CB4856 | -121,79 | 24,69 | 390 | -4,93 | 4,74E-04 |
| M9,JU751 - serotonin,ED3005 | -216,92 | 24,69 | 390 | -8,79 | <0,0001 |
| M9,JU751 - serotonin,JU1200 | -135,79 | 22,54 | 390 | -6,02 | 1,68E-06 |
| M9,JU751 - serotonin,JU2587 | 76,42 | 24,69 | 390 | 3,09 | 3,22E-01 |
| M9,JU751 - serotonin,JU2593 | 87,89 | 23,42 | 390 | 3,75 | 5,43E-02 |
| M9,JU751 - serotonin,JU2829 | 118,13 | 24,69 | 390 | 4,78 | 9,39E-04 |
| M9,JU751 - serotonin,JU3166 | -205,42 | 24,69 | 390 | -8,32 | <0,0001 |
| M9,JU751 - serotonin,JU440 | -181,62 | 24,69 | 390 | -7,36 | 4,75E-10 |
| M9,JU751 - serotonin,JU751 | 78,50 | 24,69 | 390 | 3,18 | 2,68E-01 |
| M9,JU751 - serotonin,JU830 | -180,96 | 24,69 | 390 | -7,33 | 5,70E-10 |
| M9,JU751 - serotonin,N2 | -163,15 | 21,38 | 390 | -7,63 | 6,36E-11 |
| M9,JU751 - serotonin,NIC1786 | -45,79 | 24,69 | 390 | -1,85 | 9,92E-01 |
| M9,JU751 - serotonin,NIC1832 | 100,58 | 25,25 | 390 | 3,98 | 2,46E-02 |
| M9,JU751 - serotonin,QG2873 | -139,37 | 24,69 | 390 | -5,64 | 1,34E-05 |
| M9,JU751 - serotonin,QX1430 | -119,23 | 21,38 | 390 | -5,58 | 1,93E-05 |
| M9,JU830 - M9,N2 | -17,52 | 21,38 | 390 | -0,82 | 1,00E+00 |
| M9,JU830 - M9,NIC1786 | 132,19 | 25,25 | 390 | 5,24 | 1,10E-04 |
| M9,JU830 - M9,NIC1832 | 59,50 | 24,69 | 390 | 2,41 | 8,35E-01 |
| M9,JU830 - M9,QG2873 | 1,58 | 24,69 | 390 | 0,06 | 1,00E+00 |
| M9,JU830 - M9,QX1430 | -37,00 | 21,38 | 390 | -1,73 | 9,97E-01 |
| M9,JU830 - serotonin,CB4856 | -130,50 | 24,69 | 390 | -5,29 | 8,59E-05 |
| M9,JU830 - serotonin,ED3005 | -225,63 | 24,69 | 390 | -9,14 | <0,0001 |
| M9,JU830 - serotonin,JU1200 | -144,50 | 22,54 | 390 | -6,41 | 1,80E-07 |
| M9,JU830 - serotonin,JU2587 | 67,71 | 24,69 | 390 | 2,74 | 5,94E-01 |
| M9,JU830 - serotonin,JU2593 | 79,18 | 23,42 | 390 | 3,38 | 1,63E-01 |
| M9,JU830 - serotonin,JU2829 | 109,42 | 24,69 | 390 | 4,43 | 4,34E-03 |
| M9,JU830 - serotonin,JU3166 | -214,13 | 24,69 | 390 | -8,67 | <0,0001 |
| M9,JU830 - serotonin,JU440 | -190,33 | 24,69 | 390 | -7,71 | 3,08E-11 |
| M9,JU830 - serotonin,JU751 | 69,79 | 24,69 | 390 | 2,83 | 5,26E-01 |

| | | | | | |
|---|---|---|---|---|---|
| M9,JU830 - serotonin,JU830 | -189,67 | 24,69 | 390 | -7,68 | 4,01E-11 |
| M9,JU830 - serotonin,N2 | -171,85 | 21,38 | 390 | -8,04 | <0,0001 |
| M9,JU830 - serotonin,NIC1786 | -54,50 | 24,69 | 390 | -2,21 | 9,28E-01 |
| M9,JU830 - serotonin,NIC1832 | 91,87 | 25,25 | 390 | 3,64 | 7,77E-02 |
| M9,JU830 - serotonin,QG2873 | -148,08 | 24,69 | 390 | -6,00 | 1,95E-06 |
| M9,JU830 - serotonin,QX1430 | -127,94 | 21,38 | 390 | -5,98 | 2,12E-06 |
| M9,N2 - M9,NIC1786 | 149,71 | 22,02 | 390 | 6,80 | 1,72E-08 |
| M9,N2 - M9,NIC1832 | 77,02 | 21,38 | 390 | 3,60 | 8,70E-02 |
| M9,N2 - M9,QG2873 | 19,10 | 21,38 | 390 | 0,89 | 1,00E+00 |
| M9,N2 - M9,QX1430 | -19,48 | 17,46 | 390 | -1,12 | 1,00E+00 |
| M9,N2 - serotonin,CB4856 | -112,98 | 21,38 | 390 | -5,28 | 8,67E-05 |
| M9,N2 - serotonin,ED3005 | -208,10 | 21,38 | 390 | -9,73 | <0,0001 |
| M9,N2 - serotonin,JU1200 | -126,98 | 18,86 | 390 | -6,73 | 2,58E-08 |
| M9,N2 - serotonin,JU2587 | 85,23 | 21,38 | 390 | 3,99 | 2,44E-02 |
| M9,N2 - serotonin,JU2593 | 96,70 | 19,91 | 390 | 4,86 | 6,71E-04 |
| M9,N2 - serotonin,JU2829 | 126,94 | 21,38 | 390 | 5,94 | 2,75E-06 |
| M9,N2 - serotonin,JU3166 | -196,60 | 21,38 | 390 | -9,19 | <0,0001 |
| M9,N2 - serotonin,JU440 | -172,81 | 21,38 | 390 | -8,08 | <0,0001 |
| M9,N2 - serotonin,JU751 | 87,31 | 21,38 | 390 | 4,08 | 1,71E-02 |
| M9,N2 - serotonin,JU830 | -172,15 | 21,38 | 390 | -8,05 | <0,0001 |
| M9,N2 - serotonin,N2 | -154,33 | 17,46 | 390 | -8,84 | <0,0001 |
| M9,N2 - serotonin,NIC1786 | -36,98 | 21,38 | 390 | -1,73 | 9,97E-01 |
| M9,N2 - serotonin,NIC1832 | 109,39 | 22,02 | 390 | 4,97 | 4,03E-04 |
| M9,N2 - serotonin,QG2873 | -130,56 | 21,38 | 390 | -6,11 | 1,06E-06 |
| M9,N2 - serotonin,QX1430 | -110,42 | 17,46 | 390 | -6,32 | 3,00E-07 |
| M9,NIC1786 - M9,NIC1832 | -72,69 | 25,25 | 390 | -2,88 | 4,83E-01 |
| M9,NIC1786 - M9,QG2873 | -130,61 | 25,25 | 390 | -5,17 | 1,50E-04 |
| M9,NIC1786 - M9,QX1430 | -169,19 | 22,02 | 390 | -7,68 | 3,96E-11 |
| M9,NIC1786 - serotonin,CB4856 | -262,69 | 25,25 | 390 | -10,40 | <0,0001 |
| M9,NIC1786 - serotonin,ED3005 | -357,81 | 25,25 | 390 | -14,17 | <0,0001 |
| M9,NIC1786 - serotonin,JU1200 | -276,69 | 23,15 | 390 | -11,95 | <0,0001 |
| M9,NIC1786 - serotonin,JU2587 | -64,48 | 25,25 | 390 | -2,55 | 7,41E-01 |
| M9,NIC1786 - serotonin,JU2593 | -53,01 | 24,01 | 390 | -2,21 | 9,28E-01 |
| M9,NIC1786 - serotonin,JU2829 | -22,77 | 25,25 | 390 | -0,90 | 1,00E+00 |
| M9,NIC1786 - serotonin,JU3166 | -346,31 | 25,25 | 390 | -13,72 | <0,0001 |
| M9,NIC1786 - serotonin,JU440 | -322,52 | 25,25 | 390 | -12,77 | <0,0001 |
| M9,NIC1786 - serotonin,JU751 | -62,40 | 25,25 | 390 | -2,47 | 7,97E-01 |
| M9,NIC1786 - serotonin,JU830 | -321,86 | 25,25 | 390 | -12,75 | <0,0001 |
| M9,NIC1786 - serotonin,N2 | -304,04 | 22,02 | 390 | -13,81 | <0,0001 |
| M9,NIC1786 - serotonin,NIC1786 | -186,69 | 25,25 | 390 | -7,39 | 3,64E-10 |
| M9,NIC1786 - serotonin,NIC1832 | -40,32 | 25,79 | 390 | -1,56 | 9,99E-01 |
| M9,NIC1786 - serotonin,QG2873 | -280,27 | 25,25 | 390 | -11,10 | <0,0001 |
| M9,NIC1786 - serotonin,QX1430 | -260,13 | 22,02 | 390 | -11,81 | <0,0001 |
| M9,NIC1832 - M9,QG2873 | -57,92 | 24,69 | 390 | -2,35 | 8,70E-01 |
| M9,NIC1832 - M9,QX1430 | -96,50 | 21,38 | 390 | -4,51 | 3,08E-03 |
| M9,NIC1832 - serotonin,CB4856 | -190,00 | 24,69 | 390 | -7,69 | 3,52E-11 |
| M9,NIC1832 - serotonin,ED3005 | -285,13 | 24,69 | 390 | -11,55 | <0,0001 |
| M9,NIC1832 - serotonin,JU1200 | -204,00 | 22,54 | 390 | -9,05 | <0,0001 |
| M9,NIC1832 - serotonin,JU2587 | 8,21 | 24,69 | 390 | 0,33 | 1,00E+00 |
| M9,NIC1832 - serotonin,JU2593 | 19,68 | 23,42 | 390 | 0,84 | 1,00E+00 |

| | | | | | |
|---|---|---|---|---|---|
| M9,NIC1832 - serotonin,JU2829 | 49,92 | 24,69 | 390 | 2,02 | 9,74E-01 |
| M9,NIC1832 - serotonin,JU3166 | -273,63 | 24,69 | 390 | -11,08 | <0,0001 |
| M9,NIC1832 - serotonin,JU440 | -249,83 | 24,69 | 390 | -10,12 | <0,0001 |
| M9,NIC1832 - serotonin,JU751 | 10,29 | 24,69 | 390 | 0,42 | 1,00E+00 |
| M9,NIC1832 - serotonin,JU830 | -249,17 | 24,69 | 390 | -10,09 | <0,0001 |
| M9,NIC1832 - serotonin,N2 | -231,35 | 21,38 | 390 | -10,82 | <0,0001 |
| M9,NIC1832 - serotonin,NIC1786 | -114,00 | 24,69 | 390 | -4,62 | 1,97E-03 |
| M9,NIC1832 - serotonin,NIC1832 | 32,37 | 25,25 | 390 | 1,28 | 1,00E+00 |
| M9,NIC1832 - serotonin,QG2873 | -207,58 | 24,69 | 390 | -8,41 | <0,0001 |
| M9,NIC1832 - serotonin,QX1430 | -187,44 | 21,38 | 390 | -8,77 | <0,0001 |
| M9,QG2873 - M9,QX1430 | -38,58 | 21,38 | 390 | -1,80 | 9,95E-01 |
| M9,QG2873 - serotonin,CB4856 | -132,08 | 24,69 | 390 | -5,35 | 6,22E-05 |
| M9,QG2873 - serotonin,ED3005 | -227,21 | 24,69 | 390 | -9,20 | <0,0001 |
| M9,QG2873 - serotonin,JU1200 | -146,08 | 22,54 | 390 | -6,48 | 1,19E-07 |
| M9,QG2873 - serotonin,JU2587 | 66,13 | 24,69 | 390 | 2,68 | 6,46E-01 |
| M9,QG2873 - serotonin,JU2593 | 77,60 | 23,42 | 390 | 3,31 | 1,94E-01 |
| M9,QG2873 - serotonin,JU2829 | 107,83 | 24,69 | 390 | 4,37 | 5,65E-03 |
| M9,QG2873 - serotonin,JU3166 | -215,71 | 24,69 | 390 | -8,74 | <0,0001 |
| M9,QG2873 - serotonin,JU440 | -191,92 | 24,69 | 390 | -7,77 | 1,43E-11 |
| M9,QG2873 - serotonin,JU751 | 68,21 | 24,69 | 390 | 2,76 | 5,78E-01 |
| M9,QG2873 - serotonin,JU830 | -191,25 | 24,69 | 390 | -7,75 | 2,04E-11 |
| M9,QG2873 - serotonin,N2 | -173,44 | 21,38 | 390 | -8,11 | <0,0001 |
| M9,QG2873 - serotonin,NIC1786 | -56,08 | 24,69 | 390 | -2,27 | 9,04E-01 |
| M9,QG2873 - serotonin,NIC1832 | 90,29 | 25,25 | 390 | 3,58 | 9,40E-02 |
| M9,QG2873 - serotonin,QG2873 | -149,67 | 24,69 | 390 | -6,06 | 1,36E-06 |
| M9,QG2873 - serotonin,QX1430 | -129,52 | 21,38 | 390 | -6,06 | 1,40E-06 |
| M9,QX1430 - serotonin,CB4856 | -93,50 | 21,38 | 390 | -4,37 | 5,53E-03 |
| M9,QX1430 - serotonin,ED3005 | -188,63 | 21,38 | 390 | -8,82 | <0,0001 |
| M9,QX1430 - serotonin,JU1200 | -107,50 | 18,86 | 390 | -5,70 | 9,97E-06 |
| M9,QX1430 - serotonin,JU2587 | 104,71 | 21,38 | 390 | 4,90 | 5,60E-04 |
| M9,QX1430 - serotonin,JU2593 | 116,18 | 19,91 | 390 | 5,84 | 4,77E-06 |
| M9,QX1430 - serotonin,JU2829 | 146,42 | 21,38 | 390 | 6,85 | 1,27E-08 |
| M9,QX1430 - serotonin,JU3166 | -177,13 | 21,38 | 390 | -8,28 | <0,0001 |
| M9,QX1430 - serotonin,JU440 | -153,33 | 21,38 | 390 | -7,17 | 1,62E-09 |
| M9,QX1430 - serotonin,JU751 | 106,79 | 21,38 | 390 | 4,99 | 3,55E-04 |
| M9,QX1430 - serotonin,JU830 | -152,67 | 21,38 | 390 | -7,14 | 1,99E-09 |
| M9,QX1430 - serotonin,N2 | -134,85 | 17,46 | 390 | -7,72 | 2,62E-11 |
| M9,QX1430 - serotonin,NIC1786 | -17,50 | 21,38 | 390 | -0,82 | 1,00E+00 |
| M9,QX1430 - serotonin,NIC1832 | 128,87 | 22,02 | 390 | 5,85 | 4,38E-06 |
| M9,QX1430 - serotonin,QG2873 | -111,08 | 21,38 | 390 | -5,19 | 1,35E-04 |
| M9,QX1430 - serotonin,QX1430 | -90,94 | 17,46 | 390 | -5,21 | 1,26E-04 |
| serotonin,CB4856 - serotonin,ED3005 | -95,13 | 24,69 | 390 | -3,85 | 3,89E-02 |
| serotonin,CB4856 - serotonin,JU1200 | -14,00 | 22,54 | 390 | -0,62 | 1,00E+00 |
| serotonin,CB4856 - serotonin,JU2587 | 198,21 | 24,69 | 390 | 8,03 | <0,0001 |
| serotonin,CB4856 - serotonin,JU2593 | 209,68 | 23,42 | 390 | 8,95 | <0,0001 |
| serotonin,CB4856 - serotonin,JU2829 | 239,92 | 24,69 | 390 | 9,72 | <0,0001 |
| serotonin,CB4856 - serotonin,JU3166 | -83,63 | 24,69 | 390 | -3,39 | 1,60E-01 |
| serotonin,CB4856 - serotonin,JU440 | -59,83 | 24,69 | 390 | -2,42 | 8,27E-01 |
| serotonin,CB4856 - serotonin,JU751 | 200,29 | 24,69 | 390 | 8,11 | <0,0001 |
| serotonin,CB4856 - serotonin,JU830 | -59,17 | 24,69 | 390 | -2,40 | 8,43E-01 |

| | | | | | |
|---|---|---|---|---|---|
| serotonin,CB4856 - serotonin,N2 | -41,35 | 21,38 | 390 | -1,93 | 9,85E-01 |
| serotonin,CB4856 - serotonin,NIC1786 | 76,00 | 24,69 | 390 | 3,08 | 3,33E-01 |
| serotonin,CB4856 - serotonin,NIC1832 | 222,37 | 25,25 | 390 | 8,81 | <0,0001 |
| serotonin,CB4856 - serotonin,QG2873 | -17,58 | 24,69 | 390 | -0,71 | 1,00E+00 |
| serotonin,CB4856 - serotonin,QX1430 | 2,56 | 21,38 | 390 | 0,12 | 1,00E+00 |
| serotonin,ED3005 - serotonin,JU1200 | 81,12 | 22,54 | 390 | 3,60 | 8,78E-02 |
| serotonin,ED3005 - serotonin,JU2587 | 293,33 | 24,69 | 390 | 11,88 | <0,0001 |
| serotonin,ED3005 - serotonin,JU2593 | 304,81 | 23,42 | 390 | 13,01 | <0,0001 |
| serotonin,ED3005 - serotonin,JU2829 | 335,04 | 24,69 | 390 | 13,57 | <0,0001 |
| serotonin,ED3005 - serotonin,JU3166 | 11,50 | 24,69 | 390 | 0,47 | 1,00E+00 |
| serotonin,ED3005 - serotonin,JU440 | 35,29 | 24,69 | 390 | 1,43 | 1,00E+00 |
| serotonin,ED3005 - serotonin,JU751 | 295,42 | 24,69 | 390 | 11,96 | <0,0001 |
| serotonin,ED3005 - serotonin,JU830 | 35,96 | 24,69 | 390 | 1,46 | 1,00E+00 |
| serotonin,ED3005 - serotonin,N2 | 53,77 | 21,38 | 390 | 2,51 | 7,68E-01 |
| serotonin,ED3005 - serotonin,NIC1786 | 171,13 | 24,69 | 390 | 6,93 | 7,54E-09 |
| serotonin,ED3005 - serotonin,NIC1832 | 317,50 | 25,25 | 390 | 12,58 | <0,0001 |
| serotonin,ED3005 - serotonin,QG2873 | 77,54 | 24,69 | 390 | 3,14 | 2,92E-01 |
| serotonin,ED3005 - serotonin,QX1430 | 97,69 | 21,38 | 390 | 4,57 | 2,43E-03 |
| serotonin,JU1200 - serotonin,JU2587 | 212,21 | 22,54 | 390 | 9,41 | <0,0001 |
| serotonin,JU1200 - serotonin,JU2593 | 223,68 | 21,14 | 390 | 10,58 | <0,0001 |
| serotonin,JU1200 - serotonin,JU2829 | 253,92 | 22,54 | 390 | 11,26 | <0,0001 |
| serotonin,JU1200 - serotonin,JU3166 | -69,62 | 22,54 | 390 | -3,09 | 3,26E-01 |
| serotonin,JU1200 - serotonin,JU440 | -45,83 | 22,54 | 390 | -2,03 | 9,72E-01 |
| serotonin,JU1200 - serotonin,JU751 | 214,29 | 22,54 | 390 | 9,51 | <0,0001 |
| serotonin,JU1200 - serotonin,JU830 | -45,17 | 22,54 | 390 | -2,00 | 9,77E-01 |
| serotonin,JU1200 - serotonin,N2 | -27,35 | 18,86 | 390 | -1,45 | 1,00E+00 |
| serotonin,JU1200 - serotonin,NIC1786 | 90,00 | 22,54 | 390 | 3,99 | 2,38E-02 |
| serotonin,JU1200 - serotonin,NIC1832 | 236,37 | 23,15 | 390 | 10,21 | <0,0001 |
| serotonin,JU1200 - serotonin,QG2873 | -3,58 | 22,54 | 390 | -0,16 | 1,00E+00 |
| serotonin,JU1200 - serotonin,QX1430 | 16,56 | 18,86 | 390 | 0,88 | 1,00E+00 |
| serotonin,JU2587 - serotonin,JU2593 | 11,47 | 23,42 | 390 | 0,49 | 1,00E+00 |
| serotonin,JU2587 - serotonin,JU2829 | 41,71 | 24,69 | 390 | 1,69 | 9,98E-01 |
| serotonin,JU2587 - serotonin,JU3166 | -281,83 | 24,69 | 390 | -11,41 | <0,0001 |
| serotonin,JU2587 - serotonin,JU440 | -258,04 | 24,69 | 390 | -10,45 | <0,0001 |
| serotonin,JU2587 - serotonin,JU751 | 2,08 | 24,69 | 390 | 0,08 | 1,00E+00 |
| serotonin,JU2587 - serotonin,JU830 | -257,38 | 24,69 | 390 | -10,42 | <0,0001 |
| serotonin,JU2587 - serotonin,N2 | -239,56 | 21,38 | 390 | -11,20 | <0,0001 |
| serotonin,JU2587 - serotonin,NIC1786 | -122,21 | 24,69 | 390 | -4,95 | 4,38E-04 |
| serotonin,JU2587 - serotonin,NIC1832 | 24,16 | 25,25 | 390 | 0,96 | 1,00E+00 |
| serotonin,JU2587 - serotonin,QG2873 | -215,79 | 24,69 | 390 | -8,74 | <0,0001 |
| serotonin,JU2587 - serotonin,QX1430 | -195,65 | 21,38 | 390 | -9,15 | <0,0001 |
| serotonin,JU2593 - serotonin,JU2829 | 30,23 | 23,42 | 390 | 1,29 | 1,00E+00 |
| serotonin,JU2593 - serotonin,JU3166 | -293,31 | 23,42 | 390 | -12,52 | <0,0001 |
| serotonin,JU2593 - serotonin,JU440 | -269,52 | 23,42 | 390 | -11,51 | <0,0001 |
| serotonin,JU2593 - serotonin,JU751 | -9,39 | 23,42 | 390 | -0,40 | 1,00E+00 |
| serotonin,JU2593 - serotonin,JU830 | -268,85 | 23,42 | 390 | -11,48 | <0,0001 |
| serotonin,JU2593 - serotonin,N2 | -251,04 | 19,91 | 390 | -12,61 | <0,0001 |
| serotonin,JU2593 - serotonin,NIC1786 | -133,68 | 23,42 | 390 | -5,71 | 9,62E-06 |
| serotonin,JU2593 - serotonin,NIC1832 | 12,69 | 24,01 | 390 | 0,53 | 1,00E+00 |
| serotonin,JU2593 - serotonin,QG2873 | -227,27 | 23,42 | 390 | -9,70 | <0,0001 |

| | | | | | |
|---|---|---|---|---|---|
| serotonin,JU2593 - serotonin,QX1430 | -207,12 | 19,91 | 390 | -10,40 | <0,0001 |
| serotonin,JU2829 - serotonin,JU3166 | -323,54 | 24,69 | 390 | -13,10 | <0,0001 |
| serotonin,JU2829 - serotonin,JU440 | -299,75 | 24,69 | 390 | -12,14 | <0,0001 |
| serotonin,JU2829 - serotonin,JU751 | -39,62 | 24,69 | 390 | -1,60 | 9,99E-01 |
| serotonin,JU2829 - serotonin,JU830 | -299,08 | 24,69 | 390 | -12,11 | <0,0001 |
| serotonin,JU2829 - serotonin,N2 | -281,27 | 21,38 | 390 | -13,15 | <0,0001 |
| serotonin,JU2829 - serotonin,NIC1786 | -163,92 | 24,69 | 390 | -6,64 | 4,60E-08 |
| serotonin,JU2829 - serotonin,NIC1832 | -17,55 | 25,25 | 390 | -0,69 | 1,00E+00 |
| serotonin,JU2829 - serotonin,QG2873 | -257,50 | 24,69 | 390 | -10,43 | <0,0001 |
| serotonin,JU2829 - serotonin,QX1430 | -237,35 | 21,38 | 390 | -11,10 | <0,0001 |
| serotonin,JU3166 - serotonin,JU440 | 23,79 | 24,69 | 390 | 0,96 | 1,00E+00 |
| serotonin,JU3166 - serotonin,JU751 | 283,92 | 24,69 | 390 | 11,50 | <0,0001 |
| serotonin,JU3166 - serotonin,JU830 | 24,46 | 24,69 | 390 | 0,99 | 1,00E+00 |
| serotonin,JU3166 - serotonin,N2 | 42,27 | 21,38 | 390 | 1,98 | 9,81E-01 |
| serotonin,JU3166 - serotonin,NIC1786 | 159,63 | 24,69 | 390 | 6,46 | 1,31E-07 |
| serotonin,JU3166 - serotonin,NIC1832 | 306,00 | 25,25 | 390 | 12,12 | <0,0001 |
| serotonin,JU3166 - serotonin,QG2873 | 66,04 | 24,69 | 390 | 2,67 | 6,49E-01 |
| serotonin,JU3166 - serotonin,QX1430 | 86,19 | 21,38 | 390 | 4,03 | 2,07E-02 |
| serotonin,JU440 - serotonin,JU751 | 260,12 | 24,69 | 390 | 10,53 | <0,0001 |
| serotonin,JU440 - serotonin,JU830 | 0,67 | 24,69 | 390 | 0,03 | 1,00E+00 |
| serotonin,JU440 - serotonin,N2 | 18,48 | 21,38 | 390 | 0,86 | 1,00E+00 |
| serotonin,JU440 - serotonin,NIC1786 | 135,83 | 24,69 | 390 | 5,50 | 2,85E-05 |
| serotonin,JU440 - serotonin,NIC1832 | 282,20 | 25,25 | 390 | 11,18 | <0,0001 |
| serotonin,JU440 - serotonin,QG2873 | 42,25 | 24,69 | 390 | 1,71 | 9,98E-01 |
| serotonin,JU440 - serotonin,QX1430 | 62,40 | 21,38 | 390 | 2,92 | 4,52E-01 |
| serotonin,JU751 - serotonin,JU830 | -259,46 | 24,69 | 390 | -10,51 | <0,0001 |
| serotonin,JU751 - serotonin,N2 | -241,65 | 21,38 | 390 | -11,30 | <0,0001 |
| serotonin,JU751 - serotonin,NIC1786 | -124,29 | 24,69 | 390 | -5,03 | 2,94E-04 |
| serotonin,JU751 - serotonin,NIC1832 | 22,08 | 25,25 | 390 | 0,87 | 1,00E+00 |
| serotonin,JU751 - serotonin,QG2873 | -217,88 | 24,69 | 390 | -8,82 | <0,0001 |
| serotonin,JU751 - serotonin,QX1430 | -197,73 | 21,38 | 390 | -9,25 | <0,0001 |
| serotonin,JU830 - serotonin,N2 | 17,81 | 21,38 | 390 | 0,83 | 1,00E+00 |
| serotonin,JU830 - serotonin,NIC1786 | 135,17 | 24,69 | 390 | 5,47 | 3,28E-05 |
| serotonin,JU830 - serotonin,NIC1832 | 281,54 | 25,25 | 390 | 11,15 | <0,0001 |
| serotonin,JU830 - serotonin,QG2873 | 41,58 | 24,69 | 390 | 1,68 | 9,98E-01 |
| serotonin,JU830 - serotonin,QX1430 | 61,73 | 21,38 | 390 | 2,89 | 4,77E-01 |
| serotonin,N2 - serotonin,NIC1786 | 117,35 | 21,38 | 390 | 5,49 | 3,06E-05 |
| serotonin,N2 - serotonin,NIC1832 | 263,73 | 22,02 | 390 | 11,98 | <0,0001 |
| serotonin,N2 - serotonin,QG2873 | 23,77 | 21,38 | 390 | 1,11 | 1,00E+00 |
| serotonin,N2 - serotonin,QX1430 | 43,92 | 17,46 | 390 | 2,52 | 7,68E-01 |
| serotonin,NIC1786 - serotonin,NIC1832 | 146,37 | 25,25 | 390 | 5,80 | 5,89E-06 |
| serotonin,NIC1786 - serotonin,QG2873 | -93,58 | 24,69 | 390 | -3,79 | 4,79E-02 |
| serotonin,NIC1786 - serotonin,QX1430 | -73,44 | 21,38 | 390 | -3,43 | 1,41E-01 |
| serotonin,NIC1832 - serotonin,QG2873 | -239,95 | 25,25 | 390 | -9,50 | <0,0001 |
| serotonin,NIC1832 - serotonin,QX1430 | -219,81 | 22,02 | 390 | -9,98 | <0,0001 |
| serotonin,QG2873 - serotonin,QX1430 | 20,15 | 21,38 | 390 | 0,94 | 1,00E+00 |

**Table S3 (accompanies Fig. 4C).** Results for statistical analyses testing for the effects of and interactions between *Strain* and *Treatment* (control versus imipramine) on egg laying (Align Rank Transform ANOVA).

| Term | Df | Sum of squares | F value | Pr(>F) | Sig |
|---|---|---|---|---|---|
| Treatment | 1 | 840834,17 | 562,00 | 9,46e-63 | *** |
| Strain | 14 | 799593,01 | 23,86 | 8,28e-37 | *** |
| Treatment x Strain | 14 | 462389,08 | 8,52 | 1,17e-14 | *** |

**Contrasts**

| contrast | estimate | SE | Df | t-ratio | p-value |
|---|---|---|---|---|---|
| imipramine,CB4856 - imipramine,ED3005 | -102,50 | 13,85 | 222 | -7,40 | 1,20E-09 |
| imipramine,CB4856 - imipramine,JU1200 | -85,67 | 13,85 | 222 | -6,19 | 1,26E-06 |
| imipramine,CB4856 - imipramine,JU2587 | -53,50 | 13,85 | 222 | -3,86 | 4,07E-02 |
| imipramine,CB4856 - imipramine,JU2593 | 0,33 | 13,85 | 222 | 0,02 | 1,00E+00 |
| imipramine,CB4856 - imipramine,JU2829 | -18,67 | 13,85 | 222 | -1,35 | 1,00E+00 |
| imipramine,CB4856 - imipramine,JU3166 | -60,38 | 11,99 | 222 | -5,03 | 3,90E-04 |
| imipramine,CB4856 - imipramine,JU440 | -24,83 | 13,85 | 222 | -1,79 | 9,95E-01 |
| imipramine,CB4856 - imipramine,JU751 | -60,33 | 13,85 | 222 | -4,36 | 6,85E-03 |
| imipramine,CB4856 - imipramine,JU830 | -83,25 | 13,85 | 222 | -6,01 | 3,18E-06 |
| imipramine,CB4856 - imipramine,N2 | -81,44 | 11,31 | 222 | -7,20 | 3,95E-09 |
| imipramine,CB4856 - imipramine,NIC1786 | -102,33 | 13,85 | 222 | -7,39 | 1,29E-09 |
| imipramine,CB4856 - imipramine,NIC1832 | -18,50 | 13,85 | 222 | -1,34 | 1,00E+00 |
| imipramine,CB4856 - imipramine,QG2873 | -35,13 | 11,99 | 222 | -2,93 | 4,47E-01 |
| imipramine,CB4856 - imipramine,QX1430 | -85,39 | 11,31 | 222 | -7,55 | 4,84E-10 |
| imipramine,CB4856 - M9,CB4856 | 101,75 | 13,85 | 222 | 7,35 | 1,67E-09 |
| imipramine,CB4856 - M9,ED3005 | 2,42 | 13,85 | 222 | 0,17 | 1,00E+00 |
| imipramine,CB4856 - M9,JU1200 | 25,75 | 13,85 | 222 | 1,86 | 9,91E-01 |
| imipramine,CB4856 - M9,JU2587 | 61,58 | 13,85 | 222 | 4,45 | 4,80E-03 |
| imipramine,CB4856 - M9,JU2593 | 104,58 | 13,85 | 222 | 7,55 | 4,83E-10 |
| imipramine,CB4856 - M9,JU2829 | 63,83 | 13,85 | 222 | 4,61 | 2,48E-03 |
| imipramine,CB4856 - M9,JU3166 | 28,54 | 11,99 | 222 | 2,38 | 8,49E-01 |
| imipramine,CB4856 - M9,JU440 | 111,00 | 13,85 | 222 | 8,01 | 2,75E-11 |
| imipramine,CB4856 - M9,JU751 | 81,00 | 13,85 | 222 | 5,85 | 7,42E-06 |
| imipramine,CB4856 - M9,JU830 | 88,42 | 13,85 | 222 | 6,38 | 4,27E-07 |
| imipramine,CB4856 - M9,N2 | 59,11 | 11,31 | 222 | 5,23 | 1,59E-04 |
| imipramine,CB4856 - M9,NIC1786 | 114,50 | 13,85 | 222 | 8,27 | 5,79E-12 |
| imipramine,CB4856 - M9,NIC1832 | 107,67 | 13,85 | 222 | 7,77 | 1,23E-10 |
| imipramine,CB4856 - M9,QG2873 | 63,29 | 11,99 | 222 | 5,28 | 1,26E-04 |
| imipramine,CB4856 - M9,QX1430 | 40,75 | 11,31 | 222 | 3,60 | 9,12E-02 |
| imipramine,ED3005 - imipramine,JU1200 | 16,83 | 13,85 | 222 | 1,22 | 1,00E+00 |
| imipramine,ED3005 - imipramine,JU2587 | 49,00 | 13,85 | 222 | 3,54 | 1,10E-01 |
| imipramine,ED3005 - imipramine,JU2593 | 102,83 | 13,85 | 222 | 7,42 | 1,04E-09 |
| imipramine,ED3005 - imipramine,JU2829 | 83,83 | 13,85 | 222 | 6,05 | 2,55E-06 |
| imipramine,ED3005 - imipramine,JU3166 | 42,13 | 11,99 | 222 | 3,51 | 1,18E-01 |
| imipramine,ED3005 - imipramine,JU440 | 77,67 | 13,85 | 222 | 5,61 | 2,52E-05 |
| imipramine,ED3005 - imipramine,JU751 | 42,17 | 13,85 | 222 | 3,04 | 3,61E-01 |
| imipramine,ED3005 - imipramine,JU830 | 19,25 | 13,85 | 222 | 1,39 | 1,00E+00 |
| imipramine,ED3005 - imipramine,N2 | 21,06 | 11,31 | 222 | 1,86 | 9,91E-01 |

| | | | | | |
|---|---|---|---|---|---|
| imipramine,ED3005 - imipramine,NIC1786 | 0,17 | 13,85 | 222 | 0,01 | 1,00E+00 |
| imipramine,ED3005 - imipramine,NIC1832 | 84,00 | 13,85 | 222 | 6,06 | 2,39E-06 |
| imipramine,ED3005 - imipramine,QG2873 | 67,38 | 11,99 | 222 | 5,62 | 2,40E-05 |
| imipramine,ED3005 - imipramine,QX1430 | 17,11 | 11,31 | 222 | 1,51 | 1,00E+00 |
| imipramine,ED3005 - M9,CB4856 | 204,25 | 13,85 | 222 | 14,75 | 2,29E-14 |
| imipramine,ED3005 - M9,ED3005 | 104,92 | 13,85 | 222 | 7,57 | 4,17E-10 |
| imipramine,ED3005 - M9,JU1200 | 128,25 | 13,85 | 222 | 9,26 | 3,21E-13 |
| imipramine,ED3005 - M9,JU2587 | 164,08 | 13,85 | 222 | 11,85 | 5,35E-14 |
| imipramine,ED3005 - M9,JU2593 | 207,08 | 13,85 | 222 | 14,95 | 2,29E-14 |
| imipramine,ED3005 - M9,JU2829 | 166,33 | 13,85 | 222 | 12,01 | 4,04E-14 |
| imipramine,ED3005 - M9,JU3166 | 131,04 | 11,99 | 222 | 10,92 | 2,02E-13 |
| imipramine,ED3005 - M9,JU440 | 213,50 | 13,85 | 222 | 15,41 | 2,29E-14 |
| imipramine,ED3005 - M9,JU751 | 183,50 | 13,85 | 222 | 13,25 | 2,30E-14 |
| imipramine,ED3005 - M9,JU830 | 190,92 | 13,85 | 222 | 13,78 | 2,29E-14 |
| imipramine,ED3005 - M9,N2 | 161,61 | 11,31 | 222 | 14,29 | 2,29E-14 |
| imipramine,ED3005 - M9,NIC1786 | 217,00 | 13,85 | 222 | 15,67 | 2,29E-14 |
| imipramine,ED3005 - M9,NIC1832 | 210,17 | 13,85 | 222 | 15,17 | 2,29E-14 |
| imipramine,ED3005 - M9,QG2873 | 165,79 | 11,99 | 222 | 13,82 | 2,29E-14 |
| imipramine,ED3005 - M9,QX1430 | 143,25 | 11,31 | 222 | 12,67 | 2,43E-14 |
| imipramine,JU1200 - imipramine,JU2587 | 32,17 | 13,85 | 222 | 2,32 | 8,79E-01 |
| imipramine,JU1200 - imipramine,JU2593 | 86,00 | 13,85 | 222 | 6,21 | 1,10E-06 |
| imipramine,JU1200 - imipramine,JU2829 | 67,00 | 13,85 | 222 | 4,84 | 9,34E-04 |
| imipramine,JU1200 - imipramine,JU3166 | 25,29 | 11,99 | 222 | 2,11 | 9,55E-01 |
| imipramine,JU1200 - imipramine,JU440 | 60,83 | 13,85 | 222 | 4,39 | 5,95E-03 |
| imipramine,JU1200 - imipramine,JU751 | 25,33 | 13,85 | 222 | 1,83 | 9,93E-01 |
| imipramine,JU1200 - imipramine,JU830 | 2,42 | 13,85 | 222 | 0,17 | 1,00E+00 |
| imipramine,JU1200 - imipramine,N2 | 4,22 | 11,31 | 222 | 0,37 | 1,00E+00 |
| imipramine,JU1200 - imipramine,NIC1786 | -16,67 | 13,85 | 222 | -1,20 | 1,00E+00 |
| imipramine,JU1200 - imipramine,NIC1832 | 67,17 | 13,85 | 222 | 4,85 | 8,86E-04 |
| imipramine,JU1200 - imipramine,QG2873 | 50,54 | 11,99 | 222 | 4,21 | 1,18E-02 |
| imipramine,JU1200 - imipramine,QX1430 | 0,28 | 11,31 | 222 | 0,02 | 1,00E+00 |
| imipramine,JU1200 - M9,CB4856 | 187,42 | 13,85 | 222 | 13,53 | 2,30E-14 |
| imipramine,JU1200 - M9,ED3005 | 88,08 | 13,85 | 222 | 6,36 | 4,87E-07 |
| imipramine,JU1200 - M9,JU1200 | 111,42 | 13,85 | 222 | 8,04 | 2,28E-11 |
| imipramine,JU1200 - M9,JU2587 | 147,25 | 13,85 | 222 | 10,63 | 2,59E-13 |
| imipramine,JU1200 - M9,JU2593 | 190,25 | 13,85 | 222 | 13,74 | 2,29E-14 |
| imipramine,JU1200 - M9,JU2829 | 149,50 | 13,85 | 222 | 10,79 | 2,37E-13 |
| imipramine,JU1200 - M9,JU3166 | 114,21 | 11,99 | 222 | 9,52 | 3,23E-13 |
| imipramine,JU1200 - M9,JU440 | 196,67 | 13,85 | 222 | 14,20 | 2,29E-14 |
| imipramine,JU1200 - M9,JU751 | 166,67 | 13,85 | 222 | 12,03 | 3,95E-14 |
| imipramine,JU1200 - M9,JU830 | 174,08 | 13,85 | 222 | 12,57 | 2,49E-14 |
| imipramine,JU1200 - M9,N2 | 144,78 | 11,31 | 222 | 12,80 | 2,40E-14 |
| imipramine,JU1200 - M9,NIC1786 | 200,17 | 13,85 | 222 | 14,45 | 2,29E-14 |
| imipramine,JU1200 - M9,NIC1832 | 193,33 | 13,85 | 222 | 13,96 | 2,29E-14 |
| imipramine,JU1200 - M9,QG2873 | 148,96 | 11,99 | 222 | 12,42 | 2,76E-14 |
| imipramine,JU1200 - M9,QX1430 | 126,42 | 11,31 | 222 | 11,18 | 1,54E-13 |
| imipramine,JU2587 - imipramine,JU2593 | 53,83 | 13,85 | 222 | 3,89 | 3,75E-02 |
| imipramine,JU2587 - imipramine,JU2829 | 34,83 | 13,85 | 222 | 2,51 | 7,66E-01 |
| imipramine,JU2587 - imipramine,JU3166 | -6,87 | 11,99 | 222 | -0,57 | 1,00E+00 |
| imipramine,JU2587 - imipramine,JU440 | 28,67 | 13,85 | 222 | 2,07 | 9,64E-01 |

| | | | | | |
|---|---|---|---|---|---|
| imipramine,JU2587 - imipramine,JU751 | -6,83 | 13,85 | 222 | -0,49 | 1,00E+00 |
| imipramine,JU2587 - imipramine,JU830 | -29,75 | 13,85 | 222 | -2,15 | 9,45E-01 |
| imipramine,JU2587 - imipramine,N2 | -27,94 | 11,31 | 222 | -2,47 | 7,95E-01 |
| imipramine,JU2587 - imipramine,NIC1786 | -48,83 | 13,85 | 222 | -3,53 | 1,14E-01 |
| imipramine,JU2587 - imipramine,NIC1832 | 35,00 | 13,85 | 222 | 2,53 | 7,58E-01 |
| imipramine,JU2587 - imipramine,QG2873 | 18,38 | 11,99 | 222 | 1,53 | 1,00E+00 |
| imipramine,JU2587 - imipramine,QX1430 | -31,89 | 11,31 | 222 | -2,82 | 5,33E-01 |
| imipramine,JU2587 - M9,CB4856 | 155,25 | 13,85 | 222 | 11,21 | 1,50E-13 |
| imipramine,JU2587 - M9,ED3005 | 55,92 | 13,85 | 222 | 4,04 | 2,24E-02 |
| imipramine,JU2587 - M9,JU1200 | 79,25 | 13,85 | 222 | 5,72 | 1,42E-05 |
| imipramine,JU2587 - M9,JU2587 | 115,08 | 13,85 | 222 | 8,31 | 4,49E-12 |
| imipramine,JU2587 - M9,JU2593 | 158,08 | 13,85 | 222 | 11,41 | 1,10E-13 |
| imipramine,JU2587 - M9,JU2829 | 117,33 | 13,85 | 222 | 8,47 | 1,78E-12 |
| imipramine,JU2587 - M9,JU3166 | 82,04 | 11,99 | 222 | 6,84 | 3,28E-08 |
| imipramine,JU2587 - M9,JU440 | 164,50 | 13,85 | 222 | 11,88 | 5,01E-14 |
| imipramine,JU2587 - M9,JU751 | 134,50 | 13,85 | 222 | 9,71 | 2,95E-13 |
| imipramine,JU2587 - M9,JU830 | 141,92 | 13,85 | 222 | 10,25 | 3,02E-13 |
| imipramine,JU2587 - M9,N2 | 112,61 | 11,31 | 222 | 9,96 | 2,93E-13 |
| imipramine,JU2587 - M9,NIC1786 | 168,00 | 13,85 | 222 | 12,13 | 3,66E-14 |
| imipramine,JU2587 - M9,NIC1832 | 161,17 | 13,85 | 222 | 11,64 | 7,49E-14 |
| imipramine,JU2587 - M9,QG2873 | 116,79 | 11,99 | 222 | 9,74 | 3,02E-13 |
| imipramine,JU2587 - M9,QX1430 | 94,25 | 11,31 | 222 | 8,33 | 3,87E-12 |
| imipramine,JU2593 - imipramine,JU2829 | -19,00 | 13,85 | 222 | -1,37 | 1,00E+00 |
| imipramine,JU2593 - imipramine,JU3166 | -60,71 | 11,99 | 222 | -5,06 | 3,43E-04 |
| imipramine,JU2593 - imipramine,JU440 | -25,17 | 13,85 | 222 | -1,82 | 9,94E-01 |
| imipramine,JU2593 - imipramine,JU751 | -60,67 | 13,85 | 222 | -4,38 | 6,24E-03 |
| imipramine,JU2593 - imipramine,JU830 | -83,58 | 13,85 | 222 | -6,03 | 2,80E-06 |
| imipramine,JU2593 - imipramine,N2 | -81,78 | 11,31 | 222 | -7,23 | 3,32E-09 |
| imipramine,JU2593 - imipramine,NIC1786 | -102,67 | 13,85 | 222 | -7,41 | 1,12E-09 |
| imipramine,JU2593 - imipramine,NIC1832 | -18,83 | 13,85 | 222 | -1,36 | 1,00E+00 |
| imipramine,JU2593 - imipramine,QG2873 | -35,46 | 11,99 | 222 | -2,96 | 4,26E-01 |
| imipramine,JU2593 - imipramine,QX1430 | -85,72 | 11,31 | 222 | -7,58 | 4,04E-10 |
| imipramine,JU2593 - M9,CB4856 | 101,42 | 13,85 | 222 | 7,32 | 1,93E-09 |
| imipramine,JU2593 - M9,ED3005 | 2,08 | 13,85 | 222 | 0,15 | 1,00E+00 |
| imipramine,JU2593 - M9,JU1200 | 25,42 | 13,85 | 222 | 1,84 | 9,93E-01 |
| imipramine,JU2593 - M9,JU2587 | 61,25 | 13,85 | 222 | 4,42 | 5,28E-03 |
| imipramine,JU2593 - M9,JU2593 | 104,25 | 13,85 | 222 | 7,53 | 5,59E-10 |
| imipramine,JU2593 - M9,JU2829 | 63,50 | 13,85 | 222 | 4,58 | 2,74E-03 |
| imipramine,JU2593 - M9,JU3166 | 28,21 | 11,99 | 222 | 2,35 | 8,64E-01 |
| imipramine,JU2593 - M9,JU440 | 110,67 | 13,85 | 222 | 7,99 | 3,20E-11 |
| imipramine,JU2593 - M9,JU751 | 80,67 | 13,85 | 222 | 5,82 | 8,39E-06 |
| imipramine,JU2593 - M9,JU830 | 88,08 | 13,85 | 222 | 6,36 | 4,87E-07 |
| imipramine,JU2593 - M9,N2 | 58,78 | 11,31 | 222 | 5,20 | 1,83E-04 |
| imipramine,JU2593 - M9,NIC1786 | 114,17 | 13,85 | 222 | 8,24 | 6,70E-12 |
| imipramine,JU2593 - M9,NIC1832 | 107,33 | 13,85 | 222 | 7,75 | 1,43E-10 |
| imipramine,JU2593 - M9,QG2873 | 62,96 | 11,99 | 222 | 5,25 | 1,44E-04 |
| imipramine,JU2593 - M9,QX1430 | 40,42 | 11,31 | 222 | 3,57 | 9,94E-02 |
| imipramine,JU2829 - imipramine,JU3166 | -41,71 | 11,99 | 222 | -3,48 | 1,30E-01 |
| imipramine,JU2829 - imipramine,JU440 | -6,17 | 13,85 | 222 | -0,45 | 1,00E+00 |
| imipramine,JU2829 - imipramine,JU751 | -41,67 | 13,85 | 222 | -3,01 | 3,87E-01 |

| | | | | | |
|---|---|---|---|---|---|
| imipramine,JU2829 - imipramine,JU830 | -64,58 | 13,85 | 222 | -4,66 | 1,97E-03 |
| imipramine,JU2829 - imipramine,N2 | -62,78 | 11,31 | 222 | -5,55 | 3,33E-05 |
| imipramine,JU2829 - imipramine,NIC1786 | -83,67 | 13,85 | 222 | -6,04 | 2,71E-06 |
| imipramine,JU2829 - imipramine,NIC1832 | 0,17 | 13,85 | 222 | 0,01 | 1,00E+00 |
| imipramine,JU2829 - imipramine,QG2873 | -16,46 | 11,99 | 222 | -1,37 | 1,00E+00 |
| imipramine,JU2829 - imipramine,QX1430 | -66,72 | 11,31 | 222 | -5,90 | 5,67E-06 |
| imipramine,JU2829 - M9,CB4856 | 120,42 | 13,85 | 222 | 8,69 | 6,53E-13 |
| imipramine,JU2829 - M9,ED3005 | 21,08 | 13,85 | 222 | 1,52 | 1,00E+00 |
| imipramine,JU2829 - M9,JU1200 | 44,42 | 13,85 | 222 | 3,21 | 2,56E-01 |
| imipramine,JU2829 - M9,JU2587 | 80,25 | 13,85 | 222 | 5,79 | 9,79E-06 |
| imipramine,JU2829 - M9,JU2593 | 123,25 | 13,85 | 222 | 8,90 | 3,96E-13 |
| imipramine,JU2829 - M9,JU2829 | 82,50 | 13,85 | 222 | 5,96 | 4,22E-06 |
| imipramine,JU2829 - M9,JU3166 | 47,21 | 11,99 | 222 | 3,94 | 3,18E-02 |
| imipramine,JU2829 - M9,JU440 | 129,67 | 13,85 | 222 | 9,36 | 3,11E-13 |
| imipramine,JU2829 - M9,JU751 | 99,67 | 13,85 | 222 | 7,20 | 4,09E-09 |
| imipramine,JU2829 - M9,JU830 | 107,08 | 13,85 | 222 | 7,73 | 1,59E-10 |
| imipramine,JU2829 - M9,N2 | 77,78 | 11,31 | 222 | 6,88 | 2,64E-08 |
| imipramine,JU2829 - M9,NIC1786 | 133,17 | 13,85 | 222 | 9,61 | 3,13E-13 |
| imipramine,JU2829 - M9,NIC1832 | 126,33 | 13,85 | 222 | 9,12 | 3,31E-13 |
| imipramine,JU2829 - M9,QG2873 | 81,96 | 11,99 | 222 | 6,83 | 3,42E-08 |
| imipramine,JU2829 - M9,QX1430 | 59,42 | 11,31 | 222 | 5,25 | 1,40E-04 |
| imipramine,JU3166 - imipramine,JU440 | 35,54 | 11,99 | 222 | 2,96 | 4,21E-01 |
| imipramine,JU3166 - imipramine,JU751 | 0,04 | 11,99 | 222 | 0,00 | 1,00E+00 |
| imipramine,JU3166 - imipramine,JU830 | -22,87 | 11,99 | 222 | -1,91 | 9,87E-01 |
| imipramine,JU3166 - imipramine,N2 | -21,07 | 8,94 | 222 | -2,36 | 8,62E-01 |
| imipramine,JU3166 - imipramine,NIC1786 | -41,96 | 11,99 | 222 | -3,50 | 1,23E-01 |
| imipramine,JU3166 - imipramine,NIC1832 | 41,87 | 11,99 | 222 | 3,49 | 1,26E-01 |
| imipramine,JU3166 - imipramine,QG2873 | 25,25 | 9,79 | 222 | 2,58 | 7,22E-01 |
| imipramine,JU3166 - imipramine,QX1430 | -25,01 | 8,94 | 222 | -2,80 | 5,50E-01 |
| imipramine,JU3166 - M9,CB4856 | 162,12 | 11,99 | 222 | 13,52 | 2,30E-14 |
| imipramine,JU3166 - M9,ED3005 | 62,79 | 11,99 | 222 | 5,23 | 1,53E-04 |
| imipramine,JU3166 - M9,JU1200 | 86,13 | 11,99 | 222 | 7,18 | 4,49E-09 |
| imipramine,JU3166 - M9,JU2587 | 121,96 | 11,99 | 222 | 10,17 | 2,86E-13 |
| imipramine,JU3166 - M9,JU2593 | 164,96 | 11,99 | 222 | 13,75 | 2,29E-14 |
| imipramine,JU3166 - M9,JU2829 | 124,21 | 11,99 | 222 | 10,36 | 2,80E-13 |
| imipramine,JU3166 - M9,JU3166 | 88,92 | 9,79 | 222 | 9,08 | 3,35E-13 |
| imipramine,JU3166 - M9,JU440 | 171,37 | 11,99 | 222 | 14,29 | 2,29E-14 |
| imipramine,JU3166 - M9,JU751 | 141,37 | 11,99 | 222 | 11,79 | 6,06E-14 |
| imipramine,JU3166 - M9,JU830 | 148,79 | 11,99 | 222 | 12,40 | 2,78E-14 |
| imipramine,JU3166 - M9,N2 | 119,49 | 8,94 | 222 | 13,36 | 2,30E-14 |
| imipramine,JU3166 - M9,NIC1786 | 174,88 | 11,99 | 222 | 14,58 | 2,29E-14 |
| imipramine,JU3166 - M9,NIC1832 | 168,04 | 11,99 | 222 | 14,01 | 2,29E-14 |
| imipramine,JU3166 - M9,QG2873 | 123,67 | 9,79 | 222 | 12,63 | 2,45E-14 |
| imipramine,JU3166 - M9,QX1430 | 101,13 | 8,94 | 222 | 11,31 | 1,28E-13 |
| imipramine,JU440 - imipramine,JU751 | -35,50 | 13,85 | 222 | -2,56 | 7,32E-01 |
| imipramine,JU440 - imipramine,JU830 | -58,42 | 13,85 | 222 | -4,22 | 1,16E-02 |
| imipramine,JU440 - imipramine,N2 | -56,61 | 11,31 | 222 | -5,01 | 4,42E-04 |
| imipramine,JU440 - imipramine,NIC1786 | -77,50 | 13,85 | 222 | -5,60 | 2,67E-05 |
| imipramine,JU440 - imipramine,NIC1832 | 6,33 | 13,85 | 222 | 0,46 | 1,00E+00 |
| imipramine,JU440 - imipramine,QG2873 | -10,29 | 11,99 | 222 | -0,86 | 1,00E+00 |

| | | | | | |
|---|---|---|---|---|---|
| imipramine,JU440 - imipramine,QX1430 | -60,56 | 11,31 | 222 | -5,35 | 8,68E-05 |
| imipramine,JU440 - M9,CB4856 | 126,58 | 13,85 | 222 | 9,14 | 3,23E-13 |
| imipramine,JU440 - M9,ED3005 | 27,25 | 13,85 | 222 | 1,97 | 9,81E-01 |
| imipramine,JU440 - M9,JU1200 | 50,58 | 13,85 | 222 | 3,65 | 7,90E-02 |
| imipramine,JU440 - M9,JU2587 | 86,42 | 13,85 | 222 | 6,24 | 9,38E-07 |
| imipramine,JU440 - M9,JU2593 | 129,42 | 13,85 | 222 | 9,34 | 3,17E-13 |
| imipramine,JU440 - M9,JU2829 | 88,67 | 13,85 | 222 | 6,40 | 3,87E-07 |
| imipramine,JU440 - M9,JU3166 | 53,37 | 11,99 | 222 | 4,45 | 4,73E-03 |
| imipramine,JU440 - M9,JU440 | 135,83 | 13,85 | 222 | 9,81 | 3,11E-13 |
| imipramine,JU440 - M9,JU751 | 105,83 | 13,85 | 222 | 7,64 | 2,78E-10 |
| imipramine,JU440 - M9,JU830 | 113,25 | 13,85 | 222 | 8,18 | 1,00E-11 |
| imipramine,JU440 - M9,N2 | 83,94 | 11,31 | 222 | 7,42 | 1,05E-09 |
| imipramine,JU440 - M9,NIC1786 | 139,33 | 13,85 | 222 | 10,06 | 3,04E-13 |
| imipramine,JU440 - M9,NIC1832 | 132,50 | 13,85 | 222 | 9,57 | 3,07E-13 |
| imipramine,JU440 - M9,QG2873 | 88,12 | 11,99 | 222 | 7,35 | 1,66E-09 |
| imipramine,JU440 - M9,QX1430 | 65,58 | 11,31 | 222 | 5,80 | 9,53E-06 |
| imipramine,JU751 - imipramine,JU830 | -22,92 | 13,85 | 222 | -1,65 | 9,98E-01 |
| imipramine,JU751 - imipramine,N2 | -21,11 | 11,31 | 222 | -1,87 | 9,90E-01 |
| imipramine,JU751 - imipramine,NIC1786 | -42,00 | 13,85 | 222 | -3,03 | 3,70E-01 |
| imipramine,JU751 - imipramine,NIC1832 | 41,83 | 13,85 | 222 | 3,02 | 3,78E-01 |
| imipramine,JU751 - imipramine,QG2873 | 25,21 | 11,99 | 222 | 2,10 | 9,57E-01 |
| imipramine,JU751 - imipramine,QX1430 | -25,06 | 11,31 | 222 | -2,22 | 9,23E-01 |
| imipramine,JU751 - M9,CB4856 | 162,08 | 13,85 | 222 | 11,70 | 6,36E-14 |
| imipramine,JU751 - M9,ED3005 | 62,75 | 13,85 | 222 | 4,53 | 3,42E-03 |
| imipramine,JU751 - M9,JU1200 | 86,08 | 13,85 | 222 | 6,22 | 1,07E-06 |
| imipramine,JU751 - M9,JU2587 | 121,92 | 13,85 | 222 | 8,80 | 4,78E-13 |
| imipramine,JU751 - M9,JU2593 | 164,92 | 13,85 | 222 | 11,91 | 4,45E-14 |
| imipramine,JU751 - M9,JU2829 | 124,17 | 13,85 | 222 | 8,96 | 3,58E-13 |
| imipramine,JU751 - M9,JU3166 | 88,88 | 11,99 | 222 | 7,41 | 1,14E-09 |
| imipramine,JU751 - M9,JU440 | 171,33 | 13,85 | 222 | 12,37 | 2,79E-14 |
| imipramine,JU751 - M9,JU751 | 141,33 | 13,85 | 222 | 10,20 | 3,01E-13 |
| imipramine,JU751 - M9,JU830 | 148,75 | 13,85 | 222 | 10,74 | 2,31E-13 |
| imipramine,JU751 - M9,N2 | 119,44 | 11,31 | 222 | 10,56 | 2,66E-13 |
| imipramine,JU751 - M9,NIC1786 | 174,83 | 13,85 | 222 | 12,62 | 2,45E-14 |
| imipramine,JU751 - M9,NIC1832 | 168,00 | 13,85 | 222 | 12,13 | 3,66E-14 |
| imipramine,JU751 - M9,QG2873 | 123,63 | 11,99 | 222 | 10,31 | 2,86E-13 |
| imipramine,JU751 - M9,QX1430 | 101,08 | 11,31 | 222 | 8,94 | 3,74E-13 |
| imipramine,JU830 - imipramine,N2 | 1,81 | 11,31 | 222 | 0,16 | 1,00E+00 |
| imipramine,JU830 - imipramine,NIC1786 | -19,08 | 13,85 | 222 | -1,38 | 1,00E+00 |
| imipramine,JU830 - imipramine,NIC1832 | 64,75 | 13,85 | 222 | 4,67 | 1,88E-03 |
| imipramine,JU830 - imipramine,QG2873 | 48,12 | 11,99 | 222 | 4,01 | 2,45E-02 |
| imipramine,JU830 - imipramine,QX1430 | -2,14 | 11,31 | 222 | -0,19 | 1,00E+00 |
| imipramine,JU830 - M9,CB4856 | 185,00 | 13,85 | 222 | 13,36 | 2,30E-14 |
| imipramine,JU830 - M9,ED3005 | 85,67 | 13,85 | 222 | 6,19 | 1,26E-06 |
| imipramine,JU830 - M9,JU1200 | 109,00 | 13,85 | 222 | 7,87 | 6,76E-11 |
| imipramine,JU830 - M9,JU2587 | 144,83 | 13,85 | 222 | 10,46 | 2,90E-13 |
| imipramine,JU830 - M9,JU2593 | 187,83 | 13,85 | 222 | 13,56 | 2,29E-14 |
| imipramine,JU830 - M9,JU2829 | 147,08 | 13,85 | 222 | 10,62 | 2,66E-13 |
| imipramine,JU830 - M9,JU3166 | 111,79 | 11,99 | 222 | 9,32 | 3,02E-13 |
| imipramine,JU830 - M9,JU440 | 194,25 | 13,85 | 222 | 14,02 | 2,29E-14 |

| | | | | | |
|---|---|---|---|---|---|
| imipramine,JU830 - M9,JU751 | 164,25 | 13,85 | 222 | 11,86 | 5,25E-14 |
| imipramine,JU830 - M9,JU830 | 171,67 | 13,85 | 222 | 12,39 | 2,79E-14 |
| imipramine,JU830 - M9,N2 | 142,36 | 11,31 | 222 | 12,59 | 2,48E-14 |
| imipramine,JU830 - M9,NIC1786 | 197,75 | 13,85 | 222 | 14,28 | 2,29E-14 |
| imipramine,JU830 - M9,NIC1832 | 190,92 | 13,85 | 222 | 13,78 | 2,29E-14 |
| imipramine,JU830 - M9,QG2873 | 146,54 | 11,99 | 222 | 12,22 | 3,08E-14 |
| imipramine,JU830 - M9,QX1430 | 124,00 | 11,31 | 222 | 10,96 | 2,01E-13 |
| imipramine,N2 - imipramine,NIC1786 | -20,89 | 11,31 | 222 | -1,85 | 9,92E-01 |
| imipramine,N2 - imipramine,NIC1832 | 62,94 | 11,31 | 222 | 5,57 | 3,09E-05 |
| imipramine,N2 - imipramine,QG2873 | 46,32 | 8,94 | 222 | 5,18 | 1,98E-04 |
| imipramine,N2 - imipramine,QX1430 | -3,94 | 8,00 | 222 | -0,49 | 1,00E+00 |
| imipramine,N2 - M9,CB4856 | 183,19 | 11,31 | 222 | 16,20 | 2,29E-14 |
| imipramine,N2 - M9,ED3005 | 83,86 | 11,31 | 222 | 7,42 | 1,10E-09 |
| imipramine,N2 - M9,JU1200 | 107,19 | 11,31 | 222 | 9,48 | 2,99E-13 |
| imipramine,N2 - M9,JU2587 | 143,03 | 11,31 | 222 | 12,65 | 2,44E-14 |
| imipramine,N2 - M9,JU2593 | 186,03 | 11,31 | 222 | 16,45 | 2,29E-14 |
| imipramine,N2 - M9,JU2829 | 145,28 | 11,31 | 222 | 12,85 | 2,38E-14 |
| imipramine,N2 - M9,JU3166 | 109,99 | 8,94 | 222 | 12,30 | 2,92E-14 |
| imipramine,N2 - M9,JU440 | 192,44 | 11,31 | 222 | 17,02 | 2,29E-14 |
| imipramine,N2 - M9,JU751 | 162,44 | 11,31 | 222 | 14,36 | 2,29E-14 |
| imipramine,N2 - M9,JU830 | 169,86 | 11,31 | 222 | 15,02 | 2,29E-14 |
| imipramine,N2 - M9,N2 | 140,56 | 8,00 | 222 | 17,58 | 2,29E-14 |
| imipramine,N2 - M9,NIC1786 | 195,94 | 11,31 | 222 | 17,33 | 2,29E-14 |
| imipramine,N2 - M9,NIC1832 | 189,11 | 11,31 | 222 | 16,72 | 2,29E-14 |
| imipramine,N2 - M9,QG2873 | 144,74 | 8,94 | 222 | 16,19 | 2,29E-14 |
| imipramine,N2 - M9,QX1430 | 122,19 | 8,00 | 222 | 15,28 | 2,29E-14 |
| imipramine,NIC1786 - imipramine,NIC1832 | 83,83 | 13,85 | 222 | 6,05 | 2,55E-06 |
| imipramine,NIC1786 - imipramine,QG2873 | 67,21 | 11,99 | 222 | 5,60 | 2,57E-05 |
| imipramine,NIC1786 - imipramine,QX1430 | 16,94 | 11,31 | 222 | 1,50 | 1,00E+00 |
| imipramine,NIC1786 - M9,CB4856 | 204,08 | 13,85 | 222 | 14,73 | 2,29E-14 |
| imipramine,NIC1786 - M9,ED3005 | 104,75 | 13,85 | 222 | 7,56 | 4,49E-10 |
| imipramine,NIC1786 - M9,JU1200 | 128,08 | 13,85 | 222 | 9,25 | 3,25E-13 |
| imipramine,NIC1786 - M9,JU2587 | 163,92 | 13,85 | 222 | 11,83 | 5,54E-14 |
| imipramine,NIC1786 - M9,JU2593 | 206,92 | 13,85 | 222 | 14,94 | 2,29E-14 |
| imipramine,NIC1786 - M9,JU2829 | 166,17 | 13,85 | 222 | 12,00 | 4,11E-14 |
| imipramine,NIC1786 - M9,JU3166 | 130,87 | 11,99 | 222 | 10,91 | 2,10E-13 |
| imipramine,NIC1786 - M9,JU440 | 213,33 | 13,85 | 222 | 15,40 | 2,29E-14 |
| imipramine,NIC1786 - M9,JU751 | 183,33 | 13,85 | 222 | 13,24 | 2,31E-14 |
| imipramine,NIC1786 - M9,JU830 | 190,75 | 13,85 | 222 | 13,77 | 2,29E-14 |
| imipramine,NIC1786 - M9,N2 | 161,44 | 11,31 | 222 | 14,28 | 2,29E-14 |
| imipramine,NIC1786 - M9,NIC1786 | 216,83 | 13,85 | 222 | 15,66 | 2,29E-14 |
| imipramine,NIC1786 - M9,NIC1832 | 210,00 | 13,85 | 222 | 15,16 | 2,29E-14 |
| imipramine,NIC1786 - M9,QG2873 | 165,63 | 11,99 | 222 | 13,81 | 2,29E-14 |
| imipramine,NIC1786 - M9,QX1430 | 143,08 | 11,31 | 222 | 12,65 | 2,44E-14 |
| imipramine,NIC1832 - imipramine,QG2873 | -16,62 | 11,99 | 222 | -1,39 | 1,00E+00 |
| imipramine,NIC1832 - imipramine,QX1430 | -66,89 | 11,31 | 222 | -5,91 | 5,25E-06 |
| imipramine,NIC1832 - M9,CB4856 | 120,25 | 13,85 | 222 | 8,68 | 6,84E-13 |
| imipramine,NIC1832 - M9,ED3005 | 20,92 | 13,85 | 222 | 1,51 | 1,00E+00 |
| imipramine,NIC1832 - M9,JU1200 | 44,25 | 13,85 | 222 | 3,19 | 2,63E-01 |
| imipramine,NIC1832 - M9,JU2587 | 80,08 | 13,85 | 222 | 5,78 | 1,04E-05 |

| | | | | | |
|---|---|---|---|---|---|
| imipramine,NIC1832 - M9,JU2593 | 123,08 | 13,85 | 222 | 8,89 | 3,87E-13 |
| imipramine,NIC1832 - M9,JU2829 | 82,33 | 13,85 | 222 | 5,94 | 4,50E-06 |
| imipramine,NIC1832 - M9,JU3166 | 47,04 | 11,99 | 222 | 3,92 | 3,34E-02 |
| imipramine,NIC1832 - M9,JU440 | 129,50 | 13,85 | 222 | 9,35 | 3,15E-13 |
| imipramine,NIC1832 - M9,JU751 | 99,50 | 13,85 | 222 | 7,18 | 4,40E-09 |
| imipramine,NIC1832 - M9,JU830 | 106,92 | 13,85 | 222 | 7,72 | 1,72E-10 |
| imipramine,NIC1832 - M9,N2 | 77,61 | 11,31 | 222 | 6,86 | 2,87E-08 |
| imipramine,NIC1832 - M9,NIC1786 | 133,00 | 13,85 | 222 | 9,60 | 3,08E-13 |
| imipramine,NIC1832 - M9,NIC1832 | 126,17 | 13,85 | 222 | 9,11 | 3,36E-13 |
| imipramine,NIC1832 - M9,QG2873 | 81,79 | 11,99 | 222 | 6,82 | 3,70E-08 |
| imipramine,NIC1832 - M9,QX1430 | 59,25 | 11,31 | 222 | 5,24 | 1,50E-04 |
| imipramine,QG2873 - imipramine,QX1430 | -50,26 | 8,94 | 222 | -5,62 | 2,34E-05 |
| imipramine,QG2873 - M9,CB4856 | 136,88 | 11,99 | 222 | 11,41 | 1,05E-13 |
| imipramine,QG2873 - M9,ED3005 | 37,54 | 11,99 | 222 | 3,13 | 3,03E-01 |
| imipramine,QG2873 - M9,JU1200 | 60,88 | 11,99 | 222 | 5,08 | 3,22E-04 |
| imipramine,QG2873 - M9,JU2587 | 96,71 | 11,99 | 222 | 8,06 | 2,04E-11 |
| imipramine,QG2873 - M9,JU2593 | 139,71 | 11,99 | 222 | 11,65 | 7,34E-14 |
| imipramine,QG2873 - M9,JU2829 | 98,96 | 11,99 | 222 | 8,25 | 6,41E-12 |
| imipramine,QG2873 - M9,JU3166 | 63,67 | 9,79 | 222 | 6,50 | 2,23E-07 |
| imipramine,QG2873 - M9,JU440 | 146,12 | 11,99 | 222 | 12,18 | 3,13E-14 |
| imipramine,QG2873 - M9,JU751 | 116,13 | 11,99 | 222 | 9,68 | 3,02E-13 |
| imipramine,QG2873 - M9,JU830 | 123,54 | 11,99 | 222 | 10,30 | 2,91E-13 |
| imipramine,QG2873 - M9,N2 | 94,24 | 8,94 | 222 | 10,54 | 2,60E-13 |
| imipramine,QG2873 - M9,NIC1786 | 149,63 | 11,99 | 222 | 12,47 | 2,70E-14 |
| imipramine,QG2873 - M9,NIC1832 | 142,79 | 11,99 | 222 | 11,90 | 4,50E-14 |
| imipramine,QG2873 - M9,QG2873 | 98,42 | 9,79 | 222 | 10,05 | 3,05E-13 |
| imipramine,QG2873 - M9,QX1430 | 75,88 | 8,94 | 222 | 8,49 | 1,65E-12 |
| imipramine,QX1430 - M9,CB4856 | 187,14 | 11,31 | 222 | 16,55 | 2,29E-14 |
| imipramine,QX1430 - M9,ED3005 | 87,81 | 11,31 | 222 | 7,76 | 1,30E-10 |
| imipramine,QX1430 - M9,JU1200 | 111,14 | 11,31 | 222 | 9,83 | 3,05E-13 |
| imipramine,QX1430 - M9,JU2587 | 146,97 | 11,31 | 222 | 13,00 | 2,32E-14 |
| imipramine,QX1430 - M9,JU2593 | 189,97 | 11,31 | 222 | 16,80 | 2,29E-14 |
| imipramine,QX1430 - M9,JU2829 | 149,22 | 11,31 | 222 | 13,20 | 2,31E-14 |
| imipramine,QX1430 - M9,JU3166 | 113,93 | 8,94 | 222 | 12,74 | 2,41E-14 |
| imipramine,QX1430 - M9,JU440 | 196,39 | 11,31 | 222 | 17,37 | 2,29E-14 |
| imipramine,QX1430 - M9,JU751 | 166,39 | 11,31 | 222 | 14,71 | 2,29E-14 |
| imipramine,QX1430 - M9,JU830 | 173,81 | 11,31 | 222 | 15,37 | 2,29E-14 |
| imipramine,QX1430 - M9,N2 | 144,50 | 8,00 | 222 | 18,07 | 2,29E-14 |
| imipramine,QX1430 - M9,NIC1786 | 199,89 | 11,31 | 222 | 17,68 | 2,29E-14 |
| imipramine,QX1430 - M9,NIC1832 | 193,06 | 11,31 | 222 | 17,07 | 2,29E-14 |
| imipramine,QX1430 - M9,QG2873 | 148,68 | 8,94 | 222 | 16,63 | 2,29E-14 |
| imipramine,QX1430 - M9,QX1430 | 126,14 | 8,00 | 222 | 15,77 | 2,29E-14 |
| M9,CB4856 - M9,ED3005 | -99,33 | 13,85 | 222 | -7,17 | 4,72E-09 |
| M9,CB4856 - M9,JU1200 | -76,00 | 13,85 | 222 | -5,49 | 4,56E-05 |
| M9,CB4856 - M9,JU2587 | -40,17 | 13,85 | 222 | -2,90 | 4,69E-01 |
| M9,CB4856 - M9,JU2593 | 2,83 | 13,85 | 222 | 0,20 | 1,00E+00 |
| M9,CB4856 - M9,JU2829 | -37,92 | 13,85 | 222 | -2,74 | 5,99E-01 |
| M9,CB4856 - M9,JU3166 | -73,21 | 11,99 | 222 | -6,10 | 1,95E-06 |
| M9,CB4856 - M9,JU440 | 9,25 | 13,85 | 222 | 0,67 | 1,00E+00 |
| M9,CB4856 - M9,JU751 | -20,75 | 13,85 | 222 | -1,50 | 1,00E+00 |

| | | | | | |
|---|---|---|---|---|---|
| M9,CB4856 - M9,JU830 | -13,33 | 13,85 | 222 | -0,96 | 1,00E+00 |
| M9,CB4856 - M9,N2 | -42,64 | 11,31 | 222 | -3,77 | 5,48E-02 |
| M9,CB4856 - M9,NIC1786 | 12,75 | 13,85 | 222 | 0,92 | 1,00E+00 |
| M9,CB4856 - M9,NIC1832 | 5,92 | 13,85 | 222 | 0,43 | 1,00E+00 |
| M9,CB4856 - M9,QG2873 | -38,46 | 11,99 | 222 | -3,21 | 2,57E-01 |
| M9,CB4856 - M9,QX1430 | -61,00 | 11,31 | 222 | -5,39 | 7,18E-05 |
| M9,ED3005 - M9,JU1200 | 23,33 | 13,85 | 222 | 1,68 | 9,98E-01 |
| M9,ED3005 - M9,JU2587 | 59,17 | 13,85 | 222 | 4,27 | 9,48E-03 |
| M9,ED3005 - M9,JU2593 | 102,17 | 13,85 | 222 | 7,38 | 1,39E-09 |
| M9,ED3005 - M9,JU2829 | 61,42 | 13,85 | 222 | 4,43 | 5,04E-03 |
| M9,ED3005 - M9,JU3166 | 26,13 | 11,99 | 222 | 2,18 | 9,36E-01 |
| M9,ED3005 - M9,JU440 | 108,58 | 13,85 | 222 | 7,84 | 8,15E-11 |
| M9,ED3005 - M9,JU751 | 78,58 | 13,85 | 222 | 5,67 | 1,80E-05 |
| M9,ED3005 - M9,JU830 | 86,00 | 13,85 | 222 | 6,21 | 1,10E-06 |
| M9,ED3005 - M9,N2 | 56,69 | 11,31 | 222 | 5,01 | 4,27E-04 |
| M9,ED3005 - M9,NIC1786 | 112,08 | 13,85 | 222 | 8,09 | 1,69E-11 |
| M9,ED3005 - M9,NIC1832 | 105,25 | 13,85 | 222 | 7,60 | 3,60E-10 |
| M9,ED3005 - M9,QG2873 | 60,88 | 11,99 | 222 | 5,08 | 3,22E-04 |
| M9,ED3005 - M9,QX1430 | 38,33 | 11,31 | 222 | 3,39 | 1,65E-01 |
| M9,JU1200 - M9,JU2587 | 35,83 | 13,85 | 222 | 2,59 | 7,15E-01 |
| M9,JU1200 - M9,JU2593 | 78,83 | 13,85 | 222 | 5,69 | 1,65E-05 |
| M9,JU1200 - M9,JU2829 | 38,08 | 13,85 | 222 | 2,75 | 5,89E-01 |
| M9,JU1200 - M9,JU3166 | 2,79 | 11,99 | 222 | 0,23 | 1,00E+00 |
| M9,JU1200 - M9,JU440 | 85,25 | 13,85 | 222 | 6,15 | 1,48E-06 |
| M9,JU1200 - M9,JU751 | 55,25 | 13,85 | 222 | 3,99 | 2,65E-02 |
| M9,JU1200 - M9,JU830 | 62,67 | 13,85 | 222 | 4,52 | 3,50E-03 |
| M9,JU1200 - M9,N2 | 33,36 | 11,31 | 222 | 2,95 | 4,30E-01 |
| M9,JU1200 - M9,NIC1786 | 88,75 | 13,85 | 222 | 6,41 | 3,74E-07 |
| M9,JU1200 - M9,NIC1832 | 81,92 | 13,85 | 222 | 5,91 | 5,26E-06 |
| M9,JU1200 - M9,QG2873 | 37,54 | 11,99 | 222 | 3,13 | 3,03E-01 |
| M9,JU1200 - M9,QX1430 | 15,00 | 11,31 | 222 | 1,33 | 1,00E+00 |
| M9,JU2587 - M9,JU2593 | 43,00 | 13,85 | 222 | 3,10 | 3,20E-01 |
| M9,JU2587 - M9,JU2829 | 2,25 | 13,85 | 222 | 0,16 | 1,00E+00 |
| M9,JU2587 - M9,JU3166 | -33,04 | 11,99 | 222 | -2,75 | 5,85E-01 |
| M9,JU2587 - M9,JU440 | 49,42 | 13,85 | 222 | 3,57 | 1,01E-01 |
| M9,JU2587 - M9,JU751 | 19,42 | 13,85 | 222 | 1,40 | 1,00E+00 |
| M9,JU2587 - M9,JU830 | 26,83 | 13,85 | 222 | 1,94 | 9,84E-01 |
| M9,JU2587 - M9,N2 | -2,47 | 11,31 | 222 | -0,22 | 1,00E+00 |
| M9,JU2587 - M9,NIC1786 | 52,92 | 13,85 | 222 | 3,82 | 4,67E-02 |
| M9,JU2587 - M9,NIC1832 | 46,08 | 13,85 | 222 | 3,33 | 1,93E-01 |
| M9,JU2587 - M9,QG2873 | 1,71 | 11,99 | 222 | 0,14 | 1,00E+00 |
| M9,JU2587 - M9,QX1430 | -20,83 | 11,31 | 222 | -1,84 | 9,92E-01 |
| M9,JU2593 - M9,JU2829 | -40,75 | 13,85 | 222 | -2,94 | 4,36E-01 |
| M9,JU2593 - M9,JU3166 | -76,04 | 11,99 | 222 | -6,34 | 5,44E-07 |
| M9,JU2593 - M9,JU440 | 6,42 | 13,85 | 222 | 0,46 | 1,00E+00 |
| M9,JU2593 - M9,JU751 | -23,58 | 13,85 | 222 | -1,70 | 9,98E-01 |
| M9,JU2593 - M9,JU830 | -16,17 | 13,85 | 222 | -1,17 | 1,00E+00 |
| M9,JU2593 - M9,N2 | -45,47 | 11,31 | 222 | -4,02 | 2,37E-02 |
| M9,JU2593 - M9,NIC1786 | 9,92 | 13,85 | 222 | 0,72 | 1,00E+00 |
| M9,JU2593 - M9,NIC1832 | 3,08 | 13,85 | 222 | 0,22 | 1,00E+00 |

| | | | | | |
|---|---|---|---|---|---|
| M9,JU2593 - M9,QG2873 | -41,29 | 11,99 | 222 | -3,44 | 1,43E-01 |
| M9,JU2593 - M9,QX1430 | -63,83 | 11,31 | 222 | -5,64 | 2,09E-05 |
| M9,JU2829 - M9,JU3166 | -35,29 | 11,99 | 222 | -2,94 | 4,36E-01 |
| M9,JU2829 - M9,JU440 | 47,17 | 13,85 | 222 | 3,41 | 1,58E-01 |
| M9,JU2829 - M9,JU751 | 17,17 | 13,85 | 222 | 1,24 | 1,00E+00 |
| M9,JU2829 - M9,JU830 | 24,58 | 13,85 | 222 | 1,77 | 9,95E-01 |
| M9,JU2829 - M9,N2 | -4,72 | 11,31 | 222 | -0,42 | 1,00E+00 |
| M9,JU2829 - M9,NIC1786 | 50,67 | 13,85 | 222 | 3,66 | 7,76E-02 |
| M9,JU2829 - M9,NIC1832 | 43,83 | 13,85 | 222 | 3,16 | 2,81E-01 |
| M9,JU2829 - M9,QG2873 | -0,54 | 11,99 | 222 | -0,05 | 1,00E+00 |
| M9,JU2829 - M9,QX1430 | -23,08 | 11,31 | 222 | -2,04 | 9,69E-01 |
| M9,JU3166 - M9,JU440 | 82,46 | 11,99 | 222 | 6,87 | 2,69E-08 |
| M9,JU3166 - M9,JU751 | 52,46 | 11,99 | 222 | 4,37 | 6,41E-03 |
| M9,JU3166 - M9,JU830 | 59,87 | 11,99 | 222 | 4,99 | 4,71E-04 |
| M9,JU3166 - M9,N2 | 30,57 | 8,94 | 222 | 3,42 | 1,52E-01 |
| M9,JU3166 - M9,NIC1786 | 85,96 | 11,99 | 222 | 7,17 | 4,88E-09 |
| M9,JU3166 - M9,NIC1832 | 79,13 | 11,99 | 222 | 6,60 | 1,31E-07 |
| M9,JU3166 - M9,QG2873 | 34,75 | 9,79 | 222 | 3,55 | 1,07E-01 |
| M9,JU3166 - M9,QX1430 | 12,21 | 8,94 | 222 | 1,37 | 1,00E+00 |
| M9,JU440 - M9,JU751 | -30,00 | 13,85 | 222 | -2,17 | 9,39E-01 |
| M9,JU440 - M9,JU830 | -22,58 | 13,85 | 222 | -1,63 | 9,99E-01 |
| M9,JU440 - M9,N2 | -51,89 | 11,31 | 222 | -4,59 | 2,70E-03 |
| M9,JU440 - M9,NIC1786 | 3,50 | 13,85 | 222 | 0,25 | 1,00E+00 |
| M9,JU440 - M9,NIC1832 | -3,33 | 13,85 | 222 | -0,24 | 1,00E+00 |
| M9,JU440 - M9,QG2873 | -47,71 | 11,99 | 222 | -3,98 | 2,76E-02 |
| M9,JU440 - M9,QX1430 | -70,25 | 11,31 | 222 | -6,21 | 1,09E-06 |
| M9,JU751 - M9,JU830 | 7,42 | 13,85 | 222 | 0,54 | 1,00E+00 |
| M9,JU751 - M9,N2 | -21,89 | 11,31 | 222 | -1,94 | 9,84E-01 |
| M9,JU751 - M9,NIC1786 | 33,50 | 13,85 | 222 | 2,42 | 8,27E-01 |
| M9,JU751 - M9,NIC1832 | 26,67 | 13,85 | 222 | 1,93 | 9,85E-01 |
| M9,JU751 - M9,QG2873 | -17,71 | 11,99 | 222 | -1,48 | 1,00E+00 |
| M9,JU751 - M9,QX1430 | -40,25 | 11,31 | 222 | -3,56 | 1,04E-01 |
| M9,JU830 - M9,N2 | -29,31 | 11,31 | 222 | -2,59 | 7,12E-01 |
| M9,JU830 - M9,NIC1786 | 26,08 | 13,85 | 222 | 1,88 | 9,89E-01 |
| M9,JU830 - M9,NIC1832 | 19,25 | 13,85 | 222 | 1,39 | 1,00E+00 |
| M9,JU830 - M9,QG2873 | -25,12 | 11,99 | 222 | -2,09 | 9,58E-01 |
| M9,JU830 - M9,QX1430 | -47,67 | 11,31 | 222 | -4,21 | 1,17E-02 |
| M9,N2 - M9,NIC1786 | 55,39 | 11,31 | 222 | 4,90 | 7,16E-04 |
| M9,N2 - M9,NIC1832 | 48,56 | 11,31 | 222 | 4,29 | 8,72E-03 |
| M9,N2 - M9,QG2873 | 4,18 | 8,94 | 222 | 0,47 | 1,00E+00 |
| M9,N2 - M9,QX1430 | -18,36 | 8,00 | 222 | -2,30 | 8,91E-01 |
| M9,NIC1786 - M9,NIC1832 | -6,83 | 13,85 | 222 | -0,49 | 1,00E+00 |
| M9,NIC1786 - M9,QG2873 | -51,21 | 11,99 | 222 | -4,27 | 9,57E-03 |
| M9,NIC1786 - M9,QX1430 | -73,75 | 11,31 | 222 | -6,52 | 1,99E-07 |
| M9,NIC1832 - M9,QG2873 | -44,38 | 11,99 | 222 | -3,70 | 6,84E-02 |
| M9,NIC1832 - M9,QX1430 | -66,92 | 11,31 | 222 | -5,92 | 5,19E-06 |
| M9,QG2873 - M9,QX1430 | -22,54 | 8,94 | 222 | -2,52 | 7,62E-01 |

**Table S4 (accompanies Fig. 4C).** Results for statistical analyses testing for the effects of and interactions between *Strain* and *Treatment* (control versus fluoxetine) on egg laying (Align Rank Transform ANOVA).

| Term | Df | Sum of squares | F value | Pr(>F) |
|---|---|---|---|---|
| Treatment | 1 | 3911655,54 | 1005,00 | 1,63E-108 |
| Strain | 14 | 2953428,29 | 30,09 | 5,43E-53 |
| Treatment x Strain | 14 | 2118873,89 | 16,35 | 2,91E-31 |

**Contrasts**

| Contrast | Estimate | Se | Df | t-ratio | p-value |
|---|---|---|---|---|---|
| fluoxetine,CB4856 - fluoxetine,ED3005 | 66,71 | 18,10 | 378 | 3,69 | 6,72E-02 |
| fluoxetine,CB4856 - fluoxetine,JU1200 | -10,54 | 18,10 | 378 | -0,58 | 1,00E+00 |
| fluoxetine,CB4856 - fluoxetine,JU2587 | 51,37 | 18,10 | 378 | 2,84 | 5,15E-01 |
| fluoxetine,CB4856 - fluoxetine,JU2593 | 83,37 | 18,10 | 378 | 4,61 | 2,07E-03 |
| fluoxetine,CB4856 - fluoxetine,JU2829 | -38,29 | 18,10 | 378 | -2,12 | 9,55E-01 |
| fluoxetine,CB4856 - fluoxetine,JU3166 | -3,62 | 18,10 | 378 | -0,20 | 1,00E+00 |
| fluoxetine,CB4856 - fluoxetine,JU440 | -5,79 | 18,10 | 378 | -0,32 | 1,00E+00 |
| fluoxetine,CB4856 - fluoxetine,JU751 | 32,96 | 18,10 | 378 | 1,82 | 9,94E-01 |
| fluoxetine,CB4856 - fluoxetine,JU830 | 37,33 | 18,10 | 378 | 2,06 | 9,67E-01 |
| fluoxetine,CB4856 - fluoxetine,N2 | -69,31 | 15,67 | 378 | -4,42 | 4,52E-03 |
| fluoxetine,CB4856 - fluoxetine,NIC1786 | -10,00 | 18,10 | 378 | -0,55 | 1,00E+00 |
| fluoxetine,CB4856 - fluoxetine,NIC1832 | 63,37 | 18,10 | 378 | 3,50 | 1,17E-01 |
| fluoxetine,CB4856 - fluoxetine,QG2873 | 70,67 | 18,10 | 378 | 3,91 | 3,25E-02 |
| fluoxetine,CB4856 - fluoxetine,QX1430 | -41,17 | 15,67 | 378 | -2,63 | 6,86E-01 |
| fluoxetine,CB4856 - M9,CB4856 | 255,00 | 18,10 | 378 | 14,09 | <0,0001 |
| fluoxetine,CB4856 - M9,ED3005 | 216,00 | 18,10 | 378 | 11,94 | <0,0001 |
| fluoxetine,CB4856 - M9,JU1200 | 227,62 | 18,10 | 378 | 12,58 | <0,0001 |
| fluoxetine,CB4856 - M9,JU2587 | 227,25 | 18,10 | 378 | 12,56 | <0,0001 |
| fluoxetine,CB4856 - M9,JU2593 | 253,62 | 18,10 | 378 | 14,02 | <0,0001 |
| fluoxetine,CB4856 - M9,JU2829 | 136,62 | 18,10 | 378 | 7,55 | 1,32E-10 |
| fluoxetine,CB4856 - M9,JU3166 | 133,00 | 18,10 | 378 | 7,35 | 5,26E-10 |
| fluoxetine,CB4856 - M9,JU440 | 215,37 | 18,10 | 378 | 11,90 | <0,0001 |
| fluoxetine,CB4856 - M9,JU751 | 250,29 | 18,10 | 378 | 13,83 | <0,0001 |
| fluoxetine,CB4856 - M9,JU830 | 182,17 | 18,10 | 378 | 10,07 | <0,0001 |
| fluoxetine,CB4856 - M9,N2 | 175,48 | 15,67 | 378 | 11,20 | <0,0001 |
| fluoxetine,CB4856 - M9,NIC1786 | 270,67 | 18,10 | 378 | 14,96 | <0,0001 |
| fluoxetine,CB4856 - M9,NIC1832 | 247,54 | 18,10 | 378 | 13,68 | <0,0001 |
| fluoxetine,CB4856 - M9,QG2873 | 179,87 | 18,10 | 378 | 9,94 | <0,0001 |
| fluoxetine,CB4856 - M9,QX1430 | 182,04 | 15,67 | 378 | 11,62 | <0,0001 |
| fluoxetine,ED3005 - fluoxetine,JU1200 | -77,25 | 18,10 | 378 | -4,27 | 8,42E-03 |
| fluoxetine,ED3005 - fluoxetine,JU2587 | -15,33 | 18,10 | 378 | -0,85 | 1,00E+00 |
| fluoxetine,ED3005 - fluoxetine,JU2593 | 16,67 | 18,10 | 378 | 0,92 | 1,00E+00 |
| fluoxetine,ED3005 - fluoxetine,JU2829 | -105,00 | 18,10 | 378 | -5,80 | 5,86E-06 |
| fluoxetine,ED3005 - fluoxetine,JU3166 | -70,33 | 18,10 | 378 | -3,89 | 3,47E-02 |
| fluoxetine,ED3005 - fluoxetine,JU440 | -72,50 | 18,10 | 378 | -4,01 | 2,27E-02 |
| fluoxetine,ED3005 - fluoxetine,JU751 | -33,75 | 18,10 | 378 | -1,87 | 9,91E-01 |
| fluoxetine,ED3005 - fluoxetine,JU830 | -29,37 | 18,10 | 378 | -1,62 | 9,99E-01 |
| fluoxetine,ED3005 - fluoxetine,N2 | -136,02 | 15,67 | 378 | -8,68 | <0,0001 |
| fluoxetine,ED3005 - fluoxetine,NIC1786 | -76,71 | 18,10 | 378 | -4,24 | 9,47E-03 |

| | | | | | |
|---|---|---|---|---|---|
| fluoxetine,ED3005 - fluoxetine,NIC1832 | -3,33 | 18,10 | 378 | -0,18 | 1,00E+00 |
| fluoxetine,ED3005 - fluoxetine,QG2873 | 3,96 | 18,10 | 378 | 0,22 | 1,00E+00 |
| fluoxetine,ED3005 - fluoxetine,QX1430 | -107,88 | 15,67 | 378 | -6,88 | 1,05E-08 |
| fluoxetine,ED3005 - M9,CB4856 | 188,29 | 18,10 | 378 | 10,41 | <0,0001 |
| fluoxetine,ED3005 - M9,ED3005 | 149,29 | 18,10 | 378 | 8,25 | <0,0001 |
| fluoxetine,ED3005 - M9,JU1200 | 160,92 | 18,10 | 378 | 8,89 | <0,0001 |
| fluoxetine,ED3005 - M9,JU2587 | 160,54 | 18,10 | 378 | 8,87 | <0,0001 |
| fluoxetine,ED3005 - M9,JU2593 | 186,92 | 18,10 | 378 | 10,33 | <0,0001 |
| fluoxetine,ED3005 - M9,JU2829 | 69,92 | 18,10 | 378 | 3,86 | 3,75E-02 |
| fluoxetine,ED3005 - M9,JU3166 | 66,29 | 18,10 | 378 | 3,66 | 7,22E-02 |
| fluoxetine,ED3005 - M9,JU440 | 148,67 | 18,10 | 378 | 8,22 | <0,0001 |
| fluoxetine,ED3005 - M9,JU751 | 183,58 | 18,10 | 378 | 10,15 | <0,0001 |
| fluoxetine,ED3005 - M9,JU830 | 115,46 | 18,10 | 378 | 6,38 | 2,22E-07 |
| fluoxetine,ED3005 - M9,N2 | 108,77 | 15,67 | 378 | 6,94 | 7,37E-09 |
| fluoxetine,ED3005 - M9,NIC1786 | 203,96 | 18,10 | 378 | 11,27 | <0,0001 |
| fluoxetine,ED3005 - M9,NIC1832 | 180,83 | 18,10 | 378 | 9,99 | <0,0001 |
| fluoxetine,ED3005 - M9,QG2873 | 113,17 | 18,10 | 378 | 6,25 | 4,65E-07 |
| fluoxetine,ED3005 - M9,QX1430 | 115,33 | 15,67 | 378 | 7,36 | 4,93E-10 |
| fluoxetine,JU1200 - fluoxetine,JU2587 | 61,92 | 18,10 | 378 | 3,42 | 1,46E-01 |
| fluoxetine,JU1200 - fluoxetine,JU2593 | 93,92 | 18,10 | 378 | 5,19 | 1,40E-04 |
| fluoxetine,JU1200 - fluoxetine,JU2829 | -27,75 | 18,10 | 378 | -1,53 | 1,00E+00 |
| fluoxetine,JU1200 - fluoxetine,JU3166 | 6,92 | 18,10 | 378 | 0,38 | 1,00E+00 |
| fluoxetine,JU1200 - fluoxetine,JU440 | 4,75 | 18,10 | 378 | 0,26 | 1,00E+00 |
| fluoxetine,JU1200 - fluoxetine,JU751 | 43,50 | 18,10 | 378 | 2,40 | 8,38E-01 |
| fluoxetine,JU1200 - fluoxetine,JU830 | 47,88 | 18,10 | 378 | 2,65 | 6,71E-01 |
| fluoxetine,JU1200 - fluoxetine,N2 | -58,77 | 15,67 | 378 | -3,75 | 5,48E-02 |
| fluoxetine,JU1200 - fluoxetine,NIC1786 | 0,54 | 18,10 | 378 | 0,03 | 1,00E+00 |
| fluoxetine,JU1200 - fluoxetine,NIC1832 | 73,92 | 18,10 | 378 | 4,08 | 1,71E-02 |
| fluoxetine,JU1200 - fluoxetine,QG2873 | 81,21 | 18,10 | 378 | 4,49 | 3,45E-03 |
| fluoxetine,JU1200 - fluoxetine,QX1430 | -30,62 | 15,67 | 378 | -1,95 | 9,83E-01 |
| fluoxetine,JU1200 - M9,CB4856 | 265,54 | 18,10 | 378 | 14,67 | <0,0001 |
| fluoxetine,JU1200 - M9,ED3005 | 226,54 | 18,10 | 378 | 12,52 | <0,0001 |
| fluoxetine,JU1200 - M9,JU1200 | 238,17 | 18,10 | 378 | 13,16 | <0,0001 |
| fluoxetine,JU1200 - M9,JU2587 | 237,79 | 18,10 | 378 | 13,14 | <0,0001 |
| fluoxetine,JU1200 - M9,JU2593 | 264,17 | 18,10 | 378 | 14,60 | <0,0001 |
| fluoxetine,JU1200 - M9,JU2829 | 147,17 | 18,10 | 378 | 8,13 | <0,0001 |
| fluoxetine,JU1200 - M9,JU3166 | 143,54 | 18,10 | 378 | 7,93 | 5,48E-14 |
| fluoxetine,JU1200 - M9,JU440 | 225,92 | 18,10 | 378 | 12,48 | <0,0001 |
| fluoxetine,JU1200 - M9,JU751 | 260,83 | 18,10 | 378 | 14,41 | <0,0001 |
| fluoxetine,JU1200 - M9,JU830 | 192,71 | 18,10 | 378 | 10,65 | <0,0001 |
| fluoxetine,JU1200 - M9,N2 | 186,02 | 15,67 | 378 | 11,87 | <0,0001 |
| fluoxetine,JU1200 - M9,NIC1786 | 281,21 | 18,10 | 378 | 15,54 | <0,0001 |
| fluoxetine,JU1200 - M9,NIC1832 | 258,08 | 18,10 | 378 | 14,26 | <0,0001 |
| fluoxetine,JU1200 - M9,QG2873 | 190,42 | 18,10 | 378 | 10,52 | <0,0001 |
| fluoxetine,JU1200 - M9,QX1430 | 192,58 | 15,67 | 378 | 12,29 | <0,0001 |
| fluoxetine,JU2587 - fluoxetine,JU2593 | 32,00 | 18,10 | 378 | 1,77 | 9,96E-01 |
| fluoxetine,JU2587 - fluoxetine,JU2829 | -89,67 | 18,10 | 378 | -4,96 | 4,31E-04 |
| fluoxetine,JU2587 - fluoxetine,JU3166 | -55,00 | 18,10 | 378 | -3,04 | 3,61E-01 |
| fluoxetine,JU2587 - fluoxetine,JU440 | -57,17 | 18,10 | 378 | -3,16 | 2,80E-01 |
| fluoxetine,JU2587 - fluoxetine,JU751 | -18,42 | 18,10 | 378 | -1,02 | 1,00E+00 |

| | | | | | |
|---|---|---|---|---|---|
| fluoxetine,JU2587 - fluoxetine,JU830 | -14,04 | 18,10 | 378 | -0,78 | 1,00E+00 |
| fluoxetine,JU2587 - fluoxetine,N2 | -120,69 | 15,67 | 378 | -7,70 | 4,11E-11 |
| fluoxetine,JU2587 - fluoxetine,NIC1786 | -61,38 | 18,10 | 378 | -3,39 | 1,58E-01 |
| fluoxetine,JU2587 - fluoxetine,NIC1832 | 12,00 | 18,10 | 378 | 0,66 | 1,00E+00 |
| fluoxetine,JU2587 - fluoxetine,QG2873 | 19,29 | 18,10 | 378 | 1,07 | 1,00E+00 |
| fluoxetine,JU2587 - fluoxetine,QX1430 | -92,54 | 15,67 | 378 | -5,91 | 3,34E-06 |
| fluoxetine,JU2587 - M9,CB4856 | 203,62 | 18,10 | 378 | 11,25 | <0,0001 |
| fluoxetine,JU2587 - M9,ED3005 | 164,62 | 18,10 | 378 | 9,10 | <0,0001 |
| fluoxetine,JU2587 - M9,JU1200 | 176,25 | 18,10 | 378 | 9,74 | <0,0001 |
| fluoxetine,JU2587 - M9,JU2587 | 175,88 | 18,10 | 378 | 9,72 | <0,0001 |
| fluoxetine,JU2587 - M9,JU2593 | 202,25 | 18,10 | 378 | 11,18 | <0,0001 |
| fluoxetine,JU2587 - M9,JU2829 | 85,25 | 18,10 | 378 | 4,71 | 1,31E-03 |
| fluoxetine,JU2587 - M9,JU3166 | 81,62 | 18,10 | 378 | 4,51 | 3,13E-03 |
| fluoxetine,JU2587 - M9,JU440 | 164,00 | 18,10 | 378 | 9,06 | <0,0001 |
| fluoxetine,JU2587 - M9,JU751 | 198,92 | 18,10 | 378 | 10,99 | <0,0001 |
| fluoxetine,JU2587 - M9,JU830 | 130,79 | 18,10 | 378 | 7,23 | 1,18E-09 |
| fluoxetine,JU2587 - M9,N2 | 124,10 | 15,67 | 378 | 7,92 | 1,07E-12 |
| fluoxetine,JU2587 - M9,NIC1786 | 219,29 | 18,10 | 378 | 12,12 | <0,0001 |
| fluoxetine,JU2587 - M9,NIC1832 | 196,17 | 18,10 | 378 | 10,84 | <0,0001 |
| fluoxetine,JU2587 - M9,QG2873 | 128,50 | 18,10 | 378 | 7,10 | 2,67E-09 |
| fluoxetine,JU2587 - M9,QX1430 | 130,67 | 15,67 | 378 | 8,34 | <0,0001 |
| fluoxetine,JU2593 - fluoxetine,JU2829 | -121,67 | 18,10 | 378 | -6,72 | 2,84E-08 |
| fluoxetine,JU2593 - fluoxetine,JU3166 | -87,00 | 18,10 | 378 | -4,81 | 8,51E-04 |
| fluoxetine,JU2593 - fluoxetine,JU440 | -89,17 | 18,10 | 378 | -4,93 | 4,91E-04 |
| fluoxetine,JU2593 - fluoxetine,JU751 | -50,42 | 18,10 | 378 | -2,79 | 5,59E-01 |
| fluoxetine,JU2593 - fluoxetine,JU830 | -46,04 | 18,10 | 378 | -2,54 | 7,47E-01 |
| fluoxetine,JU2593 - fluoxetine,N2 | -152,69 | 15,67 | 378 | -9,74 | <0,0001 |
| fluoxetine,JU2593 - fluoxetine,NIC1786 | -93,38 | 18,10 | 378 | -5,16 | 1,62E-04 |
| fluoxetine,JU2593 - fluoxetine,NIC1832 | -20,00 | 18,10 | 378 | -1,11 | 1,00E+00 |
| fluoxetine,JU2593 - fluoxetine,QG2873 | -12,71 | 18,10 | 378 | -0,70 | 1,00E+00 |
| fluoxetine,JU2593 - fluoxetine,QX1430 | -124,54 | 15,67 | 378 | -7,95 | <0,0001 |
| fluoxetine,JU2593 - M9,CB4856 | 171,62 | 18,10 | 378 | 9,48 | <0,0001 |
| fluoxetine,JU2593 - M9,ED3005 | 132,62 | 18,10 | 378 | 7,33 | 6,04E-10 |
| fluoxetine,JU2593 - M9,JU1200 | 144,25 | 18,10 | 378 | 7,97 | <0,0001 |
| fluoxetine,JU2593 - M9,JU2587 | 143,88 | 18,10 | 378 | 7,95 | <0,0001 |
| fluoxetine,JU2593 - M9,JU2593 | 170,25 | 18,10 | 378 | 9,41 | <0,0001 |
| fluoxetine,JU2593 - M9,JU2829 | 53,25 | 18,10 | 378 | 2,94 | 4,33E-01 |
| fluoxetine,JU2593 - M9,JU3166 | 49,62 | 18,10 | 378 | 2,74 | 5,94E-01 |
| fluoxetine,JU2593 - M9,JU440 | 132,00 | 18,10 | 378 | 7,29 | 7,59E-10 |
| fluoxetine,JU2593 - M9,JU751 | 166,92 | 18,10 | 378 | 9,22 | <0,0001 |
| fluoxetine,JU2593 - M9,JU830 | 98,79 | 18,10 | 378 | 5,46 | 3,60E-05 |
| fluoxetine,JU2593 - M9,N2 | 92,10 | 15,67 | 378 | 5,88 | 3,89E-06 |
| fluoxetine,JU2593 - M9,NIC1786 | 187,29 | 18,10 | 378 | 10,35 | <0,0001 |
| fluoxetine,JU2593 - M9,NIC1832 | 164,17 | 18,10 | 378 | 9,07 | <0,0001 |
| fluoxetine,JU2593 - M9,QG2873 | 96,50 | 18,10 | 378 | 5,33 | 6,86E-05 |
| fluoxetine,JU2593 - M9,QX1430 | 98,67 | 15,67 | 378 | 6,30 | 3,64E-07 |
| fluoxetine,JU2829 - fluoxetine,JU3166 | 34,67 | 18,10 | 378 | 1,92 | 9,87E-01 |
| fluoxetine,JU2829 - fluoxetine,JU440 | 32,50 | 18,10 | 378 | 1,80 | 9,95E-01 |
| fluoxetine,JU2829 - fluoxetine,JU751 | 71,25 | 18,10 | 378 | 3,94 | 2,91E-02 |
| fluoxetine,JU2829 - fluoxetine,JU830 | 75,63 | 18,10 | 378 | 4,18 | 1,19E-02 |

| | | | | | |
|---|---|---|---|---|---|
| fluoxetine,JU2829 - fluoxetine,N2 | -31,02 | 15,67 | 378 | -1,98 | 9,80E-01 |
| fluoxetine,JU2829 - fluoxetine,NIC1786 | 28,29 | 18,10 | 378 | 1,56 | 9,99E-01 |
| fluoxetine,JU2829 - fluoxetine,NIC1832 | 101,67 | 18,10 | 378 | 5,62 | 1,57E-05 |
| fluoxetine,JU2829 - fluoxetine,QG2873 | 108,96 | 18,10 | 378 | 6,02 | 1,75E-06 |
| fluoxetine,JU2829 - fluoxetine,QX1430 | -2,87 | 15,67 | 378 | -0,18 | 1,00E+00 |
| fluoxetine,JU2829 - M9,CB4856 | 293,29 | 18,10 | 378 | 16,21 | <0,0001 |
| fluoxetine,JU2829 - M9,ED3005 | 254,29 | 18,10 | 378 | 14,05 | <0,0001 |
| fluoxetine,JU2829 - M9,JU1200 | 265,92 | 18,10 | 378 | 14,70 | <0,0001 |
| fluoxetine,JU2829 - M9,JU2587 | 265,54 | 18,10 | 378 | 14,67 | <0,0001 |
| fluoxetine,JU2829 - M9,JU2593 | 291,92 | 18,10 | 378 | 16,13 | <0,0001 |
| fluoxetine,JU2829 - M9,JU2829 | 174,92 | 18,10 | 378 | 9,67 | <0,0001 |
| fluoxetine,JU2829 - M9,JU3166 | 171,29 | 18,10 | 378 | 9,47 | <0,0001 |
| fluoxetine,JU2829 - M9,JU440 | 253,67 | 18,10 | 378 | 14,02 | <0,0001 |
| fluoxetine,JU2829 - M9,JU751 | 288,58 | 18,10 | 378 | 15,95 | <0,0001 |
| fluoxetine,JU2829 - M9,JU830 | 220,46 | 18,10 | 378 | 12,18 | <0,0001 |
| fluoxetine,JU2829 - M9,N2 | 213,77 | 15,67 | 378 | 13,64 | <0,0001 |
| fluoxetine,JU2829 - M9,NIC1786 | 308,96 | 18,10 | 378 | 17,07 | <0,0001 |
| fluoxetine,JU2829 - M9,NIC1832 | 285,83 | 18,10 | 378 | 15,80 | <0,0001 |
| fluoxetine,JU2829 - M9,QG2873 | 218,17 | 18,10 | 378 | 12,06 | <0,0001 |
| fluoxetine,JU2829 - M9,QX1430 | 220,33 | 15,67 | 378 | 14,06 | <0,0001 |
| fluoxetine,JU3166 - fluoxetine,JU440 | -2,17 | 18,10 | 378 | -0,12 | 1,00E+00 |
| fluoxetine,JU3166 - fluoxetine,JU751 | 36,58 | 18,10 | 378 | 2,02 | 9,74E-01 |
| fluoxetine,JU3166 - fluoxetine,JU830 | 40,96 | 18,10 | 378 | 2,26 | 9,07E-01 |
| fluoxetine,JU3166 - fluoxetine,N2 | -65,69 | 15,67 | 378 | -4,19 | 1,14E-02 |
| fluoxetine,JU3166 - fluoxetine,NIC1786 | -6,38 | 18,10 | 378 | -0,35 | 1,00E+00 |
| fluoxetine,JU3166 - fluoxetine,NIC1832 | 67,00 | 18,10 | 378 | 3,70 | 6,38E-02 |
| fluoxetine,JU3166 - fluoxetine,QG2873 | 74,29 | 18,10 | 378 | 4,11 | 1,58E-02 |
| fluoxetine,JU3166 - fluoxetine,QX1430 | -37,54 | 15,67 | 378 | -2,40 | 8,43E-01 |
| fluoxetine,JU3166 - M9,CB4856 | 258,62 | 18,10 | 378 | 14,29 | <0,0001 |
| fluoxetine,JU3166 - M9,ED3005 | 219,62 | 18,10 | 378 | 12,14 | <0,0001 |
| fluoxetine,JU3166 - M9,JU1200 | 231,25 | 18,10 | 378 | 12,78 | <0,0001 |
| fluoxetine,JU3166 - M9,JU2587 | 230,87 | 18,10 | 378 | 12,76 | <0,0001 |
| fluoxetine,JU3166 - M9,JU2593 | 257,25 | 18,10 | 378 | 14,22 | <0,0001 |
| fluoxetine,JU3166 - M9,JU2829 | 140,25 | 18,10 | 378 | 7,75 | 2,64E-11 |
| fluoxetine,JU3166 - M9,JU3166 | 136,62 | 18,10 | 378 | 7,55 | 1,32E-10 |
| fluoxetine,JU3166 - M9,JU440 | 219,00 | 18,10 | 378 | 12,10 | <0,0001 |
| fluoxetine,JU3166 - M9,JU751 | 253,92 | 18,10 | 378 | 14,03 | <0,0001 |
| fluoxetine,JU3166 - M9,JU830 | 185,79 | 18,10 | 378 | 10,27 | <0,0001 |
| fluoxetine,JU3166 - M9,N2 | 179,10 | 15,67 | 378 | 11,43 | <0,0001 |
| fluoxetine,JU3166 - M9,NIC1786 | 274,29 | 18,10 | 378 | 15,16 | <0,0001 |
| fluoxetine,JU3166 - M9,NIC1832 | 251,17 | 18,10 | 378 | 13,88 | <0,0001 |
| fluoxetine,JU3166 - M9,QG2873 | 183,50 | 18,10 | 378 | 10,14 | <0,0001 |
| fluoxetine,JU3166 - M9,QX1430 | 185,67 | 15,67 | 378 | 11,85 | <0,0001 |
| fluoxetine,JU440 - fluoxetine,JU751 | 38,75 | 18,10 | 378 | 2,14 | 9,48E-01 |
| fluoxetine,JU440 - fluoxetine,JU830 | 43,13 | 18,10 | 378 | 2,38 | 8,50E-01 |
| fluoxetine,JU440 - fluoxetine,N2 | -63,52 | 15,67 | 378 | -4,05 | 1,92E-02 |
| fluoxetine,JU440 - fluoxetine,NIC1786 | -4,21 | 18,10 | 378 | -0,23 | 1,00E+00 |
| fluoxetine,JU440 - fluoxetine,NIC1832 | 69,17 | 18,10 | 378 | 3,82 | 4,32E-02 |
| fluoxetine,JU440 - fluoxetine,QG2873 | 76,46 | 18,10 | 378 | 4,23 | 1,00E-02 |
| fluoxetine,JU440 - fluoxetine,QX1430 | -35,38 | 15,67 | 378 | -2,26 | 9,10E-01 |

| | | | | | |
|---|---|---|---|---|---|
| fluoxetine,JU440 - M9,CB4856 | 260,79 | 18,10 | 378 | 14,41 | <0,0001 |
| fluoxetine,JU440 - M9,ED3005 | 221,79 | 18,10 | 378 | 12,26 | <0,0001 |
| fluoxetine,JU440 - M9,JU1200 | 233,42 | 18,10 | 378 | 12,90 | <0,0001 |
| fluoxetine,JU440 - M9,JU2587 | 233,04 | 18,10 | 378 | 12,88 | <0,0001 |
| fluoxetine,JU440 - M9,JU2593 | 259,42 | 18,10 | 378 | 14,34 | <0,0001 |
| fluoxetine,JU440 - M9,JU2829 | 142,42 | 18,10 | 378 | 7,87 | 5,74E-12 |
| fluoxetine,JU440 - M9,JU3166 | 138,79 | 18,10 | 378 | 7,67 | 5,33E-11 |
| fluoxetine,JU440 - M9,JU440 | 221,17 | 18,10 | 378 | 12,22 | <0,0001 |
| fluoxetine,JU440 - M9,JU751 | 256,08 | 18,10 | 378 | 14,15 | <0,0001 |
| fluoxetine,JU440 - M9,JU830 | 187,96 | 18,10 | 378 | 10,39 | <0,0001 |
| fluoxetine,JU440 - M9,N2 | 181,27 | 15,67 | 378 | 11,57 | <0,0001 |
| fluoxetine,JU440 - M9,NIC1786 | 276,46 | 18,10 | 378 | 15,28 | <0,0001 |
| fluoxetine,JU440 - M9,NIC1832 | 253,33 | 18,10 | 378 | 14,00 | <0,0001 |
| fluoxetine,JU440 - M9,QG2873 | 185,67 | 18,10 | 378 | 10,26 | <0,0001 |
| fluoxetine,JU440 - M9,QX1430 | 187,83 | 15,67 | 378 | 11,99 | <0,0001 |
| fluoxetine,JU751 - fluoxetine,JU830 | 4,37 | 18,10 | 378 | 0,24 | 1,00E+00 |
| fluoxetine,JU751 - fluoxetine,N2 | -102,27 | 15,67 | 378 | -6,53 | 9,38E-08 |
| fluoxetine,JU751 - fluoxetine,NIC1786 | -42,96 | 18,10 | 378 | -2,37 | 8,55E-01 |
| fluoxetine,JU751 - fluoxetine,NIC1832 | 30,42 | 18,10 | 378 | 1,68 | 9,98E-01 |
| fluoxetine,JU751 - fluoxetine,QG2873 | 37,71 | 18,10 | 378 | 2,08 | 9,62E-01 |
| fluoxetine,JU751 - fluoxetine,QX1430 | -74,13 | 15,67 | 378 | -4,73 | 1,21E-03 |
| fluoxetine,JU751 - M9,CB4856 | 222,04 | 18,10 | 378 | 12,27 | <0,0001 |
| fluoxetine,JU751 - M9,ED3005 | 183,04 | 18,10 | 378 | 10,12 | <0,0001 |
| fluoxetine,JU751 - M9,JU1200 | 194,67 | 18,10 | 378 | 10,76 | <0,0001 |
| fluoxetine,JU751 - M9,JU2587 | 194,29 | 18,10 | 378 | 10,74 | <0,0001 |
| fluoxetine,JU751 - M9,JU2593 | 220,67 | 18,10 | 378 | 12,19 | <0,0001 |
| fluoxetine,JU751 - M9,JU2829 | 103,67 | 18,10 | 378 | 5,73 | 8,72E-06 |
| fluoxetine,JU751 - M9,JU3166 | 100,04 | 18,10 | 378 | 5,53 | 2,52E-05 |
| fluoxetine,JU751 - M9,JU440 | 182,42 | 18,10 | 378 | 10,08 | <0,0001 |
| fluoxetine,JU751 - M9,JU751 | 217,33 | 18,10 | 378 | 12,01 | <0,0001 |
| fluoxetine,JU751 - M9,JU830 | 149,21 | 18,10 | 378 | 8,25 | <0,0001 |
| fluoxetine,JU751 - M9,N2 | 142,52 | 15,67 | 378 | 9,09 | <0,0001 |
| fluoxetine,JU751 - M9,NIC1786 | 237,71 | 18,10 | 378 | 13,14 | <0,0001 |
| fluoxetine,JU751 - M9,NIC1832 | 214,58 | 18,10 | 378 | 11,86 | <0,0001 |
| fluoxetine,JU751 - M9,QG2873 | 146,92 | 18,10 | 378 | 8,12 | <0,0001 |
| fluoxetine,JU751 - M9,QX1430 | 149,08 | 15,67 | 378 | 9,51 | <0,0001 |
| fluoxetine,JU830 - FLUOXETINE,N2 | -106,65 | 15,67 | 378 | -6,81 | 1,72E-08 |
| fluoxetine,JU830 - fluoxetine,NIC1786 | -47,33 | 18,10 | 378 | -2,62 | 6,95E-01 |
| fluoxetine,JU830 - fluoxetine,NIC1832 | 26,04 | 18,10 | 378 | 1,44 | 1,00E+00 |
| fluoxetine,JU830 - fluoxetine,QG2873 | 33,33 | 18,10 | 378 | 1,84 | 9,93E-01 |
| fluoxetine,JU830 - fluoxetine,QX1430 | -78,50 | 15,67 | 378 | -5,01 | 3,34E-04 |
| fluoxetine,JU830 - M9,CB4856 | 217,67 | 18,10 | 378 | 12,03 | <0,0001 |
| fluoxetine,JU830 - M9,ED3005 | 178,67 | 18,10 | 378 | 9,87 | <0,0001 |
| fluoxetine,JU830 - M9,JU1200 | 190,29 | 18,10 | 378 | 10,52 | <0,0001 |
| fluoxetine,JU830 - M9,JU2587 | 189,92 | 18,10 | 378 | 10,50 | <0,0001 |
| fluoxetine,JU830 - M9,JU2593 | 216,29 | 18,10 | 378 | 11,95 | <0,0001 |
| fluoxetine,JU830 - M9,JU2829 | 99,29 | 18,10 | 378 | 5,49 | 3,12E-05 |
| fluoxetine,JU830 - M9,JU3166 | 95,67 | 18,10 | 378 | 5,29 | 8,65E-05 |
| fluoxetine,JU830 - M9,JU440 | 178,04 | 18,10 | 378 | 9,84 | <0,0001 |
| fluoxetine,JU830 - M9,JU751 | 212,96 | 18,10 | 378 | 11,77 | <0,0001 |

| | | | | | |
|---|---|---|---|---|---|
| fluoxetine,JU830 - M9,JU830 | 144,83 | 18,10 | 378 | 8,00 | <0,0001 |
| fluoxetine,JU830 - M9,N2 | 138,15 | 15,67 | 378 | 8,82 | <0,0001 |
| fluoxetine,JU830 - M9,NIC1786 | 233,33 | 18,10 | 378 | 12,89 | <0,0001 |
| fluoxetine,JU830 - M9,NIC1832 | 210,21 | 18,10 | 378 | 11,62 | <0,0001 |
| fluoxetine,JU830 - M9,QG2873 | 142,54 | 18,10 | 378 | 7,88 | 4,98E-12 |
| fluoxetine,JU830 - M9,QX1430 | 144,71 | 15,67 | 378 | 9,23 | <0,0001 |
| fluoxetine,N2 - fluoxetine,NIC1786 | 59,31 | 15,67 | 378 | 3,78 | 4,89E-02 |
| fluoxetine,N2 - fluoxetine,NIC1832 | 132,69 | 15,67 | 378 | 8,47 | <0,0001 |
| fluoxetine,N2 - fluoxetine,QG2873 | 139,98 | 15,67 | 378 | 8,93 | <0,0001 |
| fluoxetine,N2 - fluoxetine,QX1430 | 28,15 | 12,80 | 378 | 2,20 | 9,31E-01 |
| fluoxetine,N2 - M9,CB4856 | 324,31 | 15,67 | 378 | 20,70 | <0,0001 |
| fluoxetine,N2 - M9,ED3005 | 285,31 | 15,67 | 378 | 18,21 | <0,0001 |
| fluoxetine,N2 - M9,JU1200 | 296,94 | 15,67 | 378 | 18,95 | <0,0001 |
| fluoxetine,N2 - M9,JU2587 | 296,56 | 15,67 | 378 | 18,92 | <0,0001 |
| fluoxetine,N2 - M9,JU2593 | 322,94 | 15,67 | 378 | 20,61 | <0,0001 |
| fluoxetine,N2 - M9,JU2829 | 205,94 | 15,67 | 378 | 13,14 | <0,0001 |
| fluoxetine,N2 - M9,JU3166 | 202,31 | 15,67 | 378 | 12,91 | <0,0001 |
| fluoxetine,N2 - M9,JU440 | 284,69 | 15,67 | 378 | 18,17 | <0,0001 |
| fluoxetine,N2 - M9,JU751 | 319,60 | 15,67 | 378 | 20,39 | <0,0001 |
| fluoxetine,N2 - M9,JU830 | 251,48 | 15,67 | 378 | 16,05 | <0,0001 |
| fluoxetine,N2 - M9,N2 | 244,79 | 12,80 | 378 | 19,13 | <0,0001 |
| fluoxetine,N2 - M9,NIC1786 | 339,98 | 15,67 | 378 | 21,69 | <0,0001 |
| fluoxetine,N2 - M9,NIC1832 | 316,85 | 15,67 | 378 | 20,22 | <0,0001 |
| fluoxetine,N2 - M9,QG2873 | 249,19 | 15,67 | 378 | 15,90 | <0,0001 |
| fluoxetine,N2 - M9,QX1430 | 251,35 | 12,80 | 378 | 19,64 | <0,0001 |
| fluoxetine,NIC1786 - fluoxetine,NIC1832 | 73,37 | 18,10 | 378 | 4,05 | 1,91E-02 |
| fluoxetine,NIC1786 - fluoxetine,QG2873 | 80,67 | 18,10 | 378 | 4,46 | 3,91E-03 |
| fluoxetine,NIC1786 - fluoxetine,QX1430 | -31,17 | 15,67 | 378 | -1,99 | 9,79E-01 |
| fluoxetine,NIC1786 - M9,CB4856 | 265,00 | 18,10 | 378 | 14,64 | <0,0001 |
| fluoxetine,NIC1786 - M9,ED3005 | 226,00 | 18,10 | 378 | 12,49 | <0,0001 |
| fluoxetine,NIC1786 - M9,JU1200 | 237,63 | 18,10 | 378 | 13,13 | <0,0001 |
| fluoxetine,NIC1786 - M9,JU2587 | 237,25 | 18,10 | 378 | 13,11 | <0,0001 |
| fluoxetine,NIC1786 - M9,JU2593 | 263,63 | 18,10 | 378 | 14,57 | <0,0001 |
| fluoxetine,NIC1786 - M9,JU2829 | 146,63 | 18,10 | 378 | 8,10 | <0,0001 |
| fluoxetine,NIC1786 - M9,JU3166 | 143,00 | 18,10 | 378 | 7,90 | 2,51E-12 |
| fluoxetine,NIC1786 - M9,JU440 | 225,38 | 18,10 | 378 | 12,45 | <0,0001 |
| fluoxetine,NIC1786 - M9,JU751 | 260,29 | 18,10 | 378 | 14,38 | <0,0001 |
| fluoxetine,NIC1786 - M9,JU830 | 192,17 | 18,10 | 378 | 10,62 | <0,0001 |
| fluoxetine,NIC1786 - M9,N2 | 185,48 | 15,67 | 378 | 11,84 | <0,0001 |
| fluoxetine,NIC1786 - M9,NIC1786 | 280,67 | 18,10 | 378 | 15,51 | <0,0001 |
| fluoxetine,NIC1786 - M9,NIC1832 | 257,54 | 18,10 | 378 | 14,23 | <0,0001 |
| fluoxetine,NIC1786 - M9,QG2873 | 189,88 | 18,10 | 378 | 10,49 | <0,0001 |
| fluoxetine,NIC1786 - M9,QX1430 | 192,04 | 15,67 | 378 | 12,25 | <0,0001 |
| fluoxetine,NIC1832 - fluoxetine,QG2873 | 7,29 | 18,10 | 378 | 0,40 | 1,00E+00 |
| fluoxetine,NIC1832 - fluoxetine,QX1430 | -104,54 | 15,67 | 378 | -6,67 | 3,91E-08 |
| fluoxetine,NIC1832 - M9,CB4856 | 191,63 | 18,10 | 378 | 10,59 | <0,0001 |
| Fluoxetine,NIC1832 - M9,ED3005 | 152,63 | 18,10 | 378 | 8,43 | <0,0001 |
| Fluoxetine,NIC1832 - M9,JU1200 | 164,25 | 18,10 | 378 | 9,08 | <0,0001 |
| Fluoxetine,NIC1832 - M9,JU2587 | 163,88 | 18,10 | 378 | 9,06 | <0,0001 |
| Fluoxetine,NIC1832 - M9,JU2593 | 190,25 | 18,10 | 378 | 10,51 | <0,0001 |

| | | | | | |
|---|---|---|---|---|---|
| Fluoxetine,NIC1832 - M9,JU2829 | 73,25 | 18,10 | 378 | 4,05 | 1,95E-02 |
| fluoxetine,NIC1832 - M9,JU3166 | 69,63 | 18,10 | 378 | 3,85 | 3,96E-02 |
| fluoxetine,NIC1832 - M9,JU440 | 152,00 | 18,10 | 378 | 8,40 | <0,0001 |
| fluoxetine,NIC1832 - M9,JU751 | 186,92 | 18,10 | 378 | 10,33 | <0,0001 |
| fluoxetine,NIC1832 - M9,JU830 | 118,79 | 18,10 | 378 | 6,56 | 7,44E-08 |
| fluoxetine,NIC1832 - M9,N2 | 112,10 | 15,67 | 378 | 7,15 | 1,90E-09 |
| fluoxetine,NIC1832 - M9,NIC1786 | 207,29 | 18,10 | 378 | 11,46 | <0,0001 |
| fluoxetine,NIC1832 - M9,NIC1832 | 184,17 | 18,10 | 378 | 10,18 | <0,0001 |
| fluoxetine,NIC1832 - M9,QG2873 | 116,50 | 18,10 | 378 | 6,44 | 1,58E-07 |
| fluoxetine,NIC1832 - M9,QX1430 | 118,67 | 15,67 | 378 | 7,57 | 1,13E-10 |
| fluoxetine,QG2873 - fluoxetine,QX1430 | -111,83 | 15,67 | 378 | -7,14 | 2,13E-09 |
| fluoxetine,QG2873 - M9,CB4856 | 184,33 | 18,10 | 378 | 10,19 | <0,0001 |
| fluoxetine,QG2873 - M9,ED3005 | 145,33 | 18,10 | 378 | 8,03 | <0,0001 |
| fluoxetine,QG2873 - M9,JU1200 | 156,96 | 18,10 | 378 | 8,67 | <0,0001 |
| fluoxetine,QG2873 - M9,JU2587 | 156,58 | 18,10 | 378 | 8,65 | <0,0001 |
| fluoxetine,QG2873 - M9,JU2593 | 182,96 | 18,10 | 378 | 10,11 | <0,0001 |
| fluoxetine,QG2873 - M9,JU2829 | 65,96 | 18,10 | 378 | 3,65 | 7,65E-02 |
| fluoxetine,QG2873 - M9,JU3166 | 62,33 | 18,10 | 378 | 3,44 | 1,37E-01 |
| fluoxetine,QG2873 - M9,JU440 | 144,71 | 18,10 | 378 | 8,00 | <0,0001 |
| fluoxetine,QG2873 - M9,JU751 | 179,62 | 18,10 | 378 | 9,93 | <0,0001 |
| fluoxetine,QG2873 - M9,JU830 | 111,50 | 18,10 | 378 | 6,16 | 7,90E-07 |
| fluoxetine,QG2873 - M9,N2 | 104,81 | 15,67 | 378 | 6,69 | 3,52E-08 |
| fluoxetine,QG2873 - M9,NIC1786 | 200,00 | 18,10 | 378 | 11,05 | <0,0001 |
| fluoxetine,QG2873 - M9,NIC1832 | 176,87 | 18,10 | 378 | 9,77 | <0,0001 |
| fluoxetine,QG2873 - M9,QG2873 | 109,21 | 18,10 | 378 | 6,04 | 1,62E-06 |
| fluoxetine,QG2873 - M9,QX1430 | 111,37 | 15,67 | 378 | 7,11 | 2,57E-09 |
| fluoxetine,QX1430 - M9,CB4856 | 296,17 | 15,67 | 378 | 18,90 | <0,0001 |
| fluoxetine,QX1430 - M9,ED3005 | 257,17 | 15,67 | 378 | 16,41 | <0,0001 |
| fluoxetine,QX1430 - M9,JU1200 | 268,79 | 15,67 | 378 | 17,15 | <0,0001 |
| fluoxetine,QX1430 - M9,JU2587 | 268,42 | 15,67 | 378 | 17,13 | <0,0001 |
| fluoxetine,QX1430 - M9,JU2593 | 294,79 | 15,67 | 378 | 18,81 | <0,0001 |
| fluoxetine,QX1430 - M9,JU2829 | 177,79 | 15,67 | 378 | 11,35 | <0,0001 |
| fluoxetine,QX1430 - M9,JU3166 | 174,17 | 15,67 | 378 | 11,11 | <0,0001 |
| fluoxetine,QX1430 - M9,JU440 | 256,54 | 15,67 | 378 | 16,37 | <0,0001 |
| fluoxetine,QX1430 - M9,JU751 | 291,46 | 15,67 | 378 | 18,60 | <0,0001 |
| fluoxetine,QX1430 - M9,JU830 | 223,33 | 15,67 | 378 | 14,25 | <0,0001 |
| fluoxetine,QX1430 - M9,N2 | 216,65 | 12,80 | 378 | 16,93 | <0,0001 |
| fluoxetine,QX1430 - M9,NIC1786 | 311,83 | 15,67 | 378 | 19,90 | <0,0001 |
| fluoxetine,QX1430 - M9,NIC1832 | 288,71 | 15,67 | 378 | 18,42 | <0,0001 |
| fluoxetine,QX1430 - M9,QG2873 | 221,04 | 15,67 | 378 | 14,11 | <0,0001 |
| fluoxetine,QX1430 - M9,QX1430 | 223,21 | 12,80 | 378 | 17,44 | <0,0001 |
| M9,CB4856 - M9,ED3005 | -39,00 | 18,10 | 378 | -2,16 | 9,45E-01 |
| M9,CB4856 - M9,JU1200 | -27,38 | 18,10 | 378 | -1,51 | 1,00E+00 |
| M9,CB4856 - M9,JU2587 | -27,75 | 18,10 | 378 | -1,53 | 1,00E+00 |
| M9,CB4856 - M9,JU2593 | -1,37 | 18,10 | 378 | -0,08 | 1,00E+00 |
| M9,CB4856 - M9,JU2829 | -118,38 | 18,10 | 378 | -6,54 | 8,54E-08 |
| M9,CB4856 - M9,JU3166 | -122,00 | 18,10 | 378 | -6,74 | 2,53E-08 |
| M9,CB4856 - M9,JU440 | -39,62 | 18,10 | 378 | -2,19 | 9,34E-01 |
| M9,CB4856 - M9,JU751 | -4,71 | 18,10 | 378 | -0,26 | 1,00E+00 |
| M9,CB4856 - M9,JU830 | -72,83 | 18,10 | 378 | -4,02 | 2,13E-02 |

| | | | | | |
|---|---|---|---|---|---|
| M9,CB4856 - M9,N2 | -79,52 | 15,67 | 378 | -5,07 | 2,45E-04 |
| M9,CB4856 - M9,NIC1786 | 15,67 | 18,10 | 378 | 0,87 | 1,00E+00 |
| M9,CB4856 - M9,NIC1832 | -7,46 | 18,10 | 378 | -0,41 | 1,00E+00 |
| M9,CB4856 - M9,QG2873 | -75,12 | 18,10 | 378 | -4,15 | 1,33E-02 |
| M9,CB4856 - M9,QX1430 | -72,96 | 15,67 | 378 | -4,66 | 1,68E-03 |
| M9,ED3005 - M9,JU1200 | 11,62 | 18,10 | 378 | 0,64 | 1,00E+00 |
| M9,ED3005 - M9,JU2587 | 11,25 | 18,10 | 378 | 0,62 | 1,00E+00 |
| M9,ED3005 - M9,JU2593 | 37,63 | 18,10 | 378 | 2,08 | 9,63E-01 |
| M9,ED3005 - M9,JU2829 | -79,37 | 18,10 | 378 | -4,39 | 5,25E-03 |
| M9,ED3005 - M9,JU3166 | -83,00 | 18,10 | 378 | -4,59 | 2,27E-03 |
| M9,ED3005 - M9,JU440 | -0,62 | 18,10 | 378 | -0,03 | 1,00E+00 |
| M9,ED3005 - M9,JU751 | 34,29 | 18,10 | 378 | 1,90 | 9,89E-01 |
| M9,ED3005 - M9,JU830 | -33,83 | 18,10 | 378 | -1,87 | 9,91E-01 |
| M9,ED3005 - M9,N2 | -40,52 | 15,67 | 378 | -2,59 | 7,17E-01 |
| M9,ED3005 - M9,NIC1786 | 54,67 | 18,10 | 378 | 3,02 | 3,74E-01 |
| M9,ED3005 - M9,NIC1832 | 31,54 | 18,10 | 378 | 1,74 | 9,97E-01 |
| M9,ED3005 - M9,QG2873 | -36,12 | 18,10 | 378 | -2,00 | 9,78E-01 |
| M9,ED3005 - M9,QX1430 | -33,96 | 15,67 | 378 | -2,17 | 9,41E-01 |
| M9,JU1200 - M9,JU2587 | -0,37 | 18,10 | 378 | -0,02 | 1,00E+00 |
| M9,JU1200 - M9,JU2593 | 26,00 | 18,10 | 378 | 1,44 | 1,00E+00 |
| M9,JU1200 - M9,JU2829 | -91,00 | 18,10 | 378 | -5,03 | 3,04E-04 |
| M9,JU1200 - M9,JU3166 | -94,62 | 18,10 | 378 | -5,23 | 1,15E-04 |
| M9,JU1200 - M9,JU440 | -12,25 | 18,10 | 378 | -0,68 | 1,00E+00 |
| M9,JU1200 - M9,JU751 | 22,67 | 18,10 | 378 | 1,25 | 1,00E+00 |
| M9,JU1200 - M9,JU830 | -45,46 | 18,10 | 378 | -2,51 | 7,70E-01 |
| M9,JU1200 - M9,N2 | -52,15 | 15,67 | 378 | -3,33 | 1,87E-01 |
| M9,JU1200 - M9,NIC1786 | 43,04 | 18,10 | 378 | 2,38 | 8,52E-01 |
| M9,JU1200 - M9,NIC1832 | 19,92 | 18,10 | 378 | 1,10 | 1,00E+00 |
| M9,JU1200 - M9,QG2873 | -47,75 | 18,10 | 378 | -2,64 | 6,77E-01 |
| M9,JU1200 - M9,QX1430 | -45,58 | 15,67 | 378 | -2,91 | 4,60E-01 |
| M9,JU2587 - M9,JU2593 | 26,38 | 18,10 | 378 | 1,46 | 1,00E+00 |
| M9,JU2587 - M9,JU2829 | -90,63 | 18,10 | 378 | -5,01 | 3,36E-04 |
| M9,JU2587 - M9,JU3166 | -94,25 | 18,10 | 378 | -5,21 | 1,28E-04 |
| M9,JU2587 - M9,JU440 | -11,88 | 18,10 | 378 | -0,66 | 1,00E+00 |
| M9,JU2587 - M9,JU751 | 23,04 | 18,10 | 378 | 1,27 | 1,00E+00 |
| M9,JU2587 - M9,JU830 | -45,08 | 18,10 | 378 | -2,49 | 7,84E-01 |
| M9,JU2587 - M9,N2 | -51,77 | 15,67 | 378 | -3,30 | 1,99E-01 |
| M9,JU2587 - M9,NIC1786 | 43,42 | 18,10 | 378 | 2,40 | 8,41E-01 |
| M9,JU2587 - M9,NIC1832 | 20,29 | 18,10 | 378 | 1,12 | 1,00E+00 |
| M9,JU2587 - M9,QG2873 | -47,38 | 18,10 | 378 | -2,62 | 6,93E-01 |
| M9,JU2587 - M9,QX1430 | -45,21 | 15,67 | 378 | -2,88 | 4,79E-01 |
| M9,JU2593 - M9,JU2829 | -117,00 | 18,10 | 378 | -6,47 | 1,34E-07 |
| M9,JU2593 - M9,JU3166 | -120,63 | 18,10 | 378 | -6,67 | 4,03E-08 |
| M9,JU2593 - M9,JU440 | -38,25 | 18,10 | 378 | -2,11 | 9,55E-01 |
| M9,JU2593 - M9,JU751 | -3,33 | 18,10 | 378 | -0,18 | 1,00E+00 |
| M9,JU2593 - M9,JU830 | -71,46 | 18,10 | 378 | -3,95 | 2,79E-02 |
| M9,JU2593 - M9,N2 | -78,15 | 15,67 | 378 | -4,99 | 3,72E-04 |
| M9,JU2593 - M9,NIC1786 | 17,04 | 18,10 | 378 | 0,94 | 1,00E+00 |
| M9,JU2593 - M9,NIC1832 | -6,08 | 18,10 | 378 | -0,34 | 1,00E+00 |
| M9,JU2593 - M9,QG2873 | -73,75 | 18,10 | 378 | -4,08 | 1,77E-02 |

| | | | | | |
|---|---|---|---|---|---|
| M9,JU2593 - M9,QX1430 | -71,58 | 15,67 | 378 | -4,57 | 2,46E-03 |
| M9,JU2829 - M9,JU3166 | -3,62 | 18,10 | 378 | -0,20 | 1,00E+00 |
| M9,JU2829 - M9,JU440 | 78,75 | 18,10 | 378 | 4,35 | 6,05E-03 |
| M9,JU2829 - M9,JU751 | 113,67 | 18,10 | 378 | 6,28 | 3,97E-07 |
| M9,JU2829 - M9,JU830 | 45,54 | 18,10 | 378 | 2,52 | 7,67E-01 |
| M9,JU2829 - M9,N2 | 38,85 | 15,67 | 378 | 2,48 | 7,92E-01 |
| M9,JU2829 - M9,NIC1786 | 134,04 | 18,10 | 378 | 7,41 | 3,57E-10 |
| M9,JU2829 - M9,NIC1832 | 110,92 | 18,10 | 378 | 6,13 | 9,50E-07 |
| M9,JU2829 - M9,QG2873 | 43,25 | 18,10 | 378 | 2,39 | 8,46E-01 |
| M9,JU2829 - M9,QX1430 | 45,42 | 15,67 | 378 | 2,90 | 4,68E-01 |
| M9,JU3166 - M9,JU440 | 82,38 | 18,10 | 378 | 4,55 | 2,63E-03 |
| M9,JU3166 - M9,JU751 | 117,29 | 18,10 | 378 | 6,48 | 1,22E-07 |
| M9,JU3166 - M9,JU830 | 49,17 | 18,10 | 378 | 2,72 | 6,15E-01 |
| M9,JU3166 - M9,N2 | 42,48 | 15,67 | 378 | 2,71 | 6,20E-01 |
| M9,JU3166 - M9,NIC1786 | 137,67 | 18,10 | 378 | 7,61 | 8,66E-11 |
| M9,JU3166 - M9,NIC1832 | 114,54 | 18,10 | 378 | 6,33 | 2,99E-07 |
| M9,JU3166 - M9,QG2873 | 46,88 | 18,10 | 378 | 2,59 | 7,14E-01 |
| M9,JU3166 - M9,QX1430 | 49,04 | 15,67 | 378 | 3,13 | 2,99E-01 |
| M9,JU440 - M9,JU751 | 34,92 | 18,10 | 378 | 1,93 | 9,86E-01 |
| M9,JU440 - M9,JU830 | -33,21 | 18,10 | 378 | -1,84 | 9,93E-01 |
| M9,JU440 - M9,N2 | -39,90 | 15,67 | 378 | -2,55 | 7,46E-01 |
| M9,JU440 - M9,NIC1786 | 55,29 | 18,10 | 378 | 3,06 | 3,49E-01 |
| M9,JU440 - M9,NIC1832 | 32,17 | 18,10 | 378 | 1,78 | 9,96E-01 |
| M9,JU440 - M9,QG2873 | -35,50 | 18,10 | 378 | -1,96 | 9,82E-01 |
| M9,JU440 - M9,QX1430 | -33,33 | 15,67 | 378 | -2,13 | 9,52E-01 |
| M9,JU751 - M9,JU830 | -68,12 | 18,10 | 378 | -3,76 | 5,23E-02 |
| M9,JU751 - M9,N2 | -74,81 | 15,67 | 378 | -4,77 | 9,93E-04 |
| M9,JU751 - M9,NIC1786 | 20,38 | 18,10 | 378 | 1,13 | 1,00E+00 |
| M9,JU751 - M9,NIC1832 | -2,75 | 18,10 | 378 | -0,15 | 1,00E+00 |
| M9,JU751 - M9,QG2873 | -70,42 | 18,10 | 378 | -3,89 | 3,41E-02 |
| M9,JU751 - M9,QX1430 | -68,25 | 15,67 | 378 | -4,36 | 5,97E-03 |
| M9,JU830 - M9,N2 | -6,69 | 15,67 | 378 | -0,43 | 1,00E+00 |
| M9,JU830 - M9,NIC1786 | 88,50 | 18,10 | 378 | 4,89 | 5,82E-04 |
| M9,JU830 - M9,NIC1832 | 65,37 | 18,10 | 378 | 3,61 | 8,44E-02 |
| M9,JU830 - M9,QG2873 | -2,29 | 18,10 | 378 | -0,13 | 1,00E+00 |
| M9,JU830 - M9,QX1430 | -0,13 | 15,67 | 378 | -0,01 | 1,00E+00 |
| M9,N2 - M9,NIC1786 | 95,19 | 15,67 | 378 | 6,07 | 1,30E-06 |
| M9,N2 - M9,NIC1832 | 72,06 | 15,67 | 378 | 4,60 | 2,15E-03 |
| M9,N2 - M9,QG2873 | 4,40 | 15,67 | 378 | 0,28 | 1,00E+00 |
| M9,N2 - M9,QX1430 | 6,56 | 12,80 | 378 | 0,51 | 1,00E+00 |
| M9,NIC1786 - M9,NIC1832 | -23,13 | 18,10 | 378 | -1,28 | 1,00E+00 |
| M9,NIC1786 - M9,QG2873 | -90,79 | 18,10 | 378 | -5,02 | 3,22E-04 |
| M9,NIC1786 - M9,QX1430 | -88,63 | 15,67 | 378 | -5,66 | 1,29E-05 |
| M9,NIC1832 - M9,QG2873 | -67,67 | 18,10 | 378 | -3,74 | 5,67E-02 |
| M9,NIC1832 - M9,QX1430 | -65,50 | 15,67 | 378 | -4,18 | 1,19E-02 |
| M9,QG2873 - M9,QX1430 | 2,17 | 15,67 | 378 | 0,14 | 1,00E+00 |

**Table S5 (accompanies Fig. 5B).** Results for statistical analyses testing for the effects of and interactions between *State* (presence versus absence of *mod-5* (*lf*)) and *Background* (strain) on egg laying in M9 (Align Rank Transform ANOVA).

| Term | Df | Sum of squares | F value | Pr(>F) | Sig |
|---|---|---|---|---|---|
| State | 1 | 6268217,46 | 175,55 | 5,84E-36 | *** |
| Background | 9 | 6768751,82 | 20,98 | 1,46E-31 | *** |
| State x Background | 9 | 2037491,32 | 5,34 | 4,34E-07 | *** |

**Contrasts**

| Contrast | Estimate | Se | Df | T-ratio | P-value |
|---|---|---|---|---|---|
| mod-5,CB4856 - mod-5,ED3005 | -82,65 | 45,14 | 709 | -1,83 | 9,55E-01 |
| mod-5,CB4856 - mod-5,JU1200 | -208,00 | 49,45 | 709 | -4,21 | 4,67E-03 |
| mod-5,CB4856 - mod-5,JU2829 | -242,31 | 45,14 | 709 | -5,37 | 2,00E-05 |
| mod-5,CB4856 - mod-5,JU3166 | -49,40 | 49,45 | 709 | -1,00 | 1,00E+00 |
| mod-5,CB4856 - mod-5,JU751 | -77,25 | 45,96 | 709 | -1,68 | 9,81E-01 |
| mod-5,CB4856 - mod-5,JU830 | 74,53 | 42,98 | 709 | 1,73 | 9,74E-01 |
| mod-5,CB4856 - mod-5,N2 | -158,09 | 41,58 | 709 | -3,80 | 2,18E-02 |
| mod-5,CB4856 - mod-5,QG2873 | 56,92 | 49,45 | 709 | 1,15 | 1,00E+00 |
| mod-5,CB4856 - mod-5,QX1430 | -89,35 | 41,69 | 709 | -2,14 | 8,33E-01 |
| mod-5,CB4856 - WT,CB4856 | 292,10 | 49,45 | 709 | 5,91 | 1,01E-06 |
| mod-5,CB4856 - WT,ED3005 | 137,80 | 45,67 | 709 | 3,02 | 2,25E-01 |
| mod-5,CB4856 - WT,JU1200 | 109,98 | 49,45 | 709 | 2,22 | 7,85E-01 |
| mod-5,CB4856 - WT,JU2829 | -34,34 | 45,14 | 709 | -0,76 | 1,00E+00 |
| mod-5,CB4856 - WT,JU3166 | 93,88 | 49,45 | 709 | 1,90 | 9,38E-01 |
| mod-5,CB4856 - WT,N2 | -31,25 | 41,38 | 709 | -0,76 | 1,00E+00 |
| mod-5,ED3005 - mod-5,JU3166 | 33,25 | 45,14 | 709 | 0,74 | 1,00E+00 |
| mod-5,ED3005 - mod-5,JU830 | 157,18 | 37,94 | 709 | 4,14 | 6,03E-03 |
| mod-5,ED3005 - mod-5,N2 | -75,44 | 36,35 | 709 | -2,08 | 8,68E-01 |
| mod-5,ED3005 - mod-5,QX1430 | -6,70 | 36,47 | 709 | -0,18 | 1,00E+00 |
| mod-5,ED3005 - WT,CB4856 | 374,75 | 45,14 | 709 | 8,30 | <0,0001 |
| mod-5,ED3005 - WT,JU1200 | 192,63 | 45,14 | 709 | 4,27 | 3,64E-03 |
| mod-5,ED3005 - WT,QG2873 | 226,49 | 45,73 | 709 | 4,95 | 1,64E-04 |
| mod-5,JU1200 - mod-5,JU3166 | 158,60 | 49,45 | 709 | 3,21 | 1,40E-01 |
| mod-5,JU1200 - mod-5,N2 | 49,91 | 41,58 | 709 | 1,20 | 1,00E+00 |
| mod-5,JU1200 - mod-5,QX1430 | 118,65 | 41,69 | 709 | 2,85 | 3,27E-01 |
| mod-5,JU1200 - WT,JU2829 | 173,66 | 45,14 | 709 | 3,85 | 1,86E-02 |
| mod-5,JU1200 - WT,JU830 | 330,14 | 43,14 | 709 | 7,65 | <0,0001 |
| mod-5,JU1200 - WT,QG2873 | 351,85 | 49,99 | 709 | 7,04 | <0,0001 |
| mod-5,JU1200 - WT,QX1430 | 341,81 | 41,48 | 709 | 8,24 | <0,0001 |
| mod-5,JU2829 - mod-5,JU830 | 316,84 | 37,94 | 709 | 8,35 | <0,0001 |
| mod-5,JU2829 - mod-5,QG2873 | 299,23 | 45,14 | 709 | 6,63 | 9,57E-09 |
| mod-5,JU2829 - mod-5,QX1430 | 152,97 | 36,47 | 709 | 4,19 | 4,90E-03 |
| mod-5,JU2829 - WT,CB4856 | 534,42 | 45,14 | 709 | 11,84 | <0,0001 |
| mod-5,JU2829 - WT,ED3005 | 380,12 | 40,97 | 709 | 9,28 | <0,0001 |
| mod-5,JU2829 - WT,JU1200 | 352,29 | 45,14 | 709 | 7,80 | <0,0001 |
| mod-5,JU2829 - WT,JU2829 | 207,97 | 40,38 | 709 | 5,15 | 6,13E-05 |
| mod-5,JU2829 - WT,JU3166 | 336,19 | 45,14 | 709 | 7,45 | <0,0001 |
| mod-5,JU2829 - WT,JU751 | 269,03 | 40,38 | 709 | 6,66 | 7,06E-09 |
| mod-5,JU2829 - WT,JU830 | 364,45 | 38,12 | 709 | 9,56 | <0,0001 |

| | | | | | |
|---|---|---|---|---|---|
| MOD-5,JU2829 - WT,N2 | 211,07 | 36,12 | 709 | 5,84 | 1,45E-06 |
| mod-5,JU2829 - WT,QG2873 | 386,16 | 45,73 | 709 | 8,44 | <0,0001 |
| mod-5,JU2829 - WT,QX1430 | 376,12 | 36,23 | 709 | 10,38 | <0,0001 |
| mod-5,JU3166 - mod-5,JU751 | -27,85 | 45,96 | 709 | -0,61 | 1,00E+00 |
| mod-5,JU3166 - mod-5,JU830 | 123,93 | 42,98 | 709 | 2,88 | 3,03E-01 |
| mod-5,JU3166 - mod-5,N2 | -108,69 | 41,58 | 709 | -2,61 | 4,95E-01 |
| mod-5,JU3166 - mod-5,QG2873 | 106,31 | 49,45 | 709 | 2,15 | 8,29E-01 |
| mod-5,JU3166 - mod-5,QX1430 | -39,95 | 41,69 | 709 | -0,96 | 1,00E+00 |
| mod-5,JU3166 - WT,CB4856 | 341,50 | 49,45 | 709 | 6,91 | <0,0001 |
| mod-5,JU3166 - WT,ED3005 | 187,20 | 45,67 | 709 | 4,10 | 7,17E-03 |
| mod-5,JU3166 - WT,JU1200 | 159,38 | 49,45 | 709 | 3,22 | 1,34E-01 |
| mod-5,JU3166 - WT,JU2829 | 15,06 | 45,14 | 709 | 0,33 | 1,00E+00 |
| mod-5,JU3166 - WT,JU3166 | 143,27 | 49,45 | 709 | 2,90 | 2,94E-01 |
| mod-5,JU3166 - WT,JU751 | 76,11 | 45,14 | 709 | 1,69 | 9,81E-01 |
| mod-5,JU3166 - WT,JU830 | 171,54 | 43,14 | 709 | 3,98 | 1,15E-02 |
| mod-5,JU3166 - WT,N2 | 18,15 | 41,38 | 709 | 0,44 | 1,00E+00 |
| mod-5,JU3166 - WT,QG2873 | 193,24 | 49,99 | 709 | 3,87 | 1,74E-02 |
| mod-5,JU3166 - WT,QX1430 | 183,21 | 41,48 | 709 | 4,42 | 1,93E-03 |
| mod-5,JU751 - mod-5,JU830 | 151,78 | 38,91 | 709 | 3,90 | 1,52E-02 |
| mod-5,JU751 - mod-5,N2 | -80,84 | 37,35 | 709 | -2,16 | 8,21E-01 |
| mod-5,JU751 - mod-5,QG2873 | 134,16 | 45,96 | 709 | 2,92 | 2,81E-01 |
| mod-5,JU751 - mod-5,QX1430 | -12,10 | 37,47 | 709 | -0,32 | 1,00E+00 |
| mod-5,JU751 - WT,CB4856 | 369,35 | 45,96 | 709 | 8,04 | <0,0001 |
| mod-5,JU751 - WT,ED3005 | 215,05 | 41,86 | 709 | 5,14 | 6,57E-05 |
| mod-5,JU751 - WT,JU1200 | 187,23 | 45,96 | 709 | 4,07 | 7,91E-03 |
| mod-5,JU751 - WT,JU2829 | 42,91 | 41,29 | 709 | 1,04 | 1,00E+00 |
| mod-5,JU751 - WT,JU3166 | 171,12 | 45,96 | 709 | 3,72 | 2,87E-02 |
| mod-5,JU751 - WT,JU751 | 103,96 | 41,29 | 709 | 2,52 | 5,70E-01 |
| mod-5,JU751 - WT,JU830 | 199,39 | 39,08 | 709 | 5,10 | 7,84E-05 |
| mod-5,JU751 - WT,N2 | 46,00 | 37,13 | 709 | 1,24 | 1,00E+00 |
| mod-5,JU751 - WT,QG2873 | 221,10 | 46,53 | 709 | 4,75 | 4,29E-04 |
| mod-5,JU751 - WT,QX1430 | 211,06 | 37,24 | 709 | 5,67 | 3,93E-06 |
| mod-5,JU830 - mod-5,N2 | -232,62 | 33,62 | 709 | -6,92 | <0,0001 |
| mod-5,JU830 - mod-5,QG2873 | -17,62 | 42,98 | 709 | -0,41 | 1,00E+00 |
| mod-5,JU830 - mod-5,QX1430 | -163,88 | 33,75 | 709 | -4,86 | 2,63E-04 |
| mod-5,JU830 - WT,CB4856 | 217,57 | 42,98 | 709 | 5,06 | 9,56E-05 |
| mod-5,JU830 - WT,ED3005 | 63,27 | 38,57 | 709 | 1,64 | 9,86E-01 |
| mod-5,JU830 - WT,JU1200 | 35,45 | 42,98 | 709 | 0,82 | 1,00E+00 |
| mod-5,JU830 - WT,JU2829 | -108,87 | 37,94 | 709 | -2,87 | 3,12E-01 |
| mod-5,JU830 - WT,JU3166 | 19,34 | 42,98 | 709 | 0,45 | 1,00E+00 |
| mod-5,JU830 - WT,JU751 | -47,82 | 37,94 | 709 | -1,26 | 9,99E-01 |
| mod-5,JU830 - WT,JU830 | 47,61 | 35,53 | 709 | 1,34 | 9,99E-01 |
| mod-5,JU830 - WT,N2 | -105,78 | 33,37 | 709 | -3,17 | 1,55E-01 |
| mod-5,JU830 - WT,QG2873 | 69,32 | 43,59 | 709 | 1,59 | 9,90E-01 |
| mod-5,JU830 - WT,QX1430 | 59,28 | 33,49 | 709 | 1,77 | 9,68E-01 |
| mod-5,N2 - mod-5,QG2873 | 215,01 | 41,58 | 709 | 5,17 | 5,53E-05 |
| mod-5,N2 - mod-5,QX1430 | 68,74 | 31,95 | 709 | 2,15 | 8,28E-01 |
| mod-5,N2 - WT,CB4856 | 450,19 | 41,58 | 709 | 10,83 | <0,0001 |
| mod-5,N2 - WT,ED3005 | 295,89 | 37,00 | 709 | 8,00 | <0,0001 |
| mod-5,N2 - WT,JU1200 | 268,07 | 41,58 | 709 | 6,45 | 3,67E-08 |

| | | | | | |
|---|---|---|---|---|---|
| mod-5,N2 - WT,JU2829 | 123,75 | 36,35 | 709 | 3,40 | 8,03E-02 |
| mod-5,N2 - WT,JU3166 | 251,96 | 41,58 | 709 | 6,06 | 4,13E-07 |
| mod-5,N2 - WT,JU751 | 184,81 | 36,35 | 709 | 5,08 | 8,57E-05 |
| mod-5,N2 - WT,JU830 | 280,23 | 33,82 | 709 | 8,29 | <0,0001 |
| mod-5,N2 - WT,N2 | 126,84 | 31,55 | 709 | 4,02 | 9,71E-03 |
| mod-5,N2 - WT,QG2873 | 301,94 | 42,21 | 709 | 7,15 | <0,0001 |
| mod-5,N2 - WT,QX1430 | 291,90 | 31,68 | 709 | 9,22 | <0,0001 |
| mod-5,QG2873 - mod-5,QX1430 | -146,26 | 41,69 | 709 | -3,51 | 5,84E-02 |
| mod-5,QG2873 - WT,CB4856 | 235,19 | 49,45 | 709 | 4,76 | 4,21E-04 |
| mod-5,QG2873 - WT,ED3005 | 80,89 | 45,67 | 709 | 1,77 | 9,68E-01 |
| mod-5,QG2873 - WT,JU1200 | 53,06 | 49,45 | 709 | 1,07 | 1,00E+00 |
| mod-5,QG2873 - WT,JU2829 | -91,26 | 45,14 | 709 | -2,02 | 8,93E-01 |
| mod-5,QG2873 - WT,JU3166 | 36,96 | 49,45 | 709 | 0,75 | 1,00E+00 |
| mod-5,QG2873 - WT,JU751 | -30,20 | 45,14 | 709 | -0,67 | 1,00E+00 |
| mod-5,QG2873 - WT,JU830 | 65,23 | 43,14 | 709 | 1,51 | 9,94E-01 |
| mod-5,QG2873 - WT,N2 | -88,16 | 41,38 | 709 | -2,13 | 8,40E-01 |
| mod-5,QG2873 - WT,QG2873 | 86,93 | 49,99 | 709 | 1,74 | 9,73E-01 |
| mod-5,QG2873 - WT,QX1430 | 76,89 | 41,48 | 709 | 1,85 | 9,50E-01 |
| mod-5,QX1430 - WT,CB4856 | 381,45 | 41,69 | 709 | 9,15 | <0,0001 |
| mod-5,QX1430 - WT,ED3005 | 227,15 | 37,12 | 709 | 6,12 | 2,90E-07 |
| mod-5,QX1430 - WT,JU1200 | 199,32 | 41,69 | 709 | 4,78 | 3,72E-04 |
| mod-5,QX1430 - WT,JU2829 | 55,01 | 36,47 | 709 | 1,51 | 9,95E-01 |
| mod-5,QX1430 - WT,JU3166 | 183,22 | 41,69 | 709 | 4,40 | 2,12E-03 |
| mod-5,QX1430 - WT,JU751 | 116,06 | 36,47 | 709 | 3,18 | 1,50E-01 |
| mod-5,QX1430 - WT,JU830 | 211,49 | 33,95 | 709 | 6,23 | 1,49E-07 |
| mod-5,QX1430 - WT,N2 | 58,10 | 31,69 | 709 | 1,83 | 9,55E-01 |
| mod-5,QX1430 - WT,QG2873 | 233,19 | 42,32 | 709 | 5,51 | 9,30E-06 |
| mod-5,QX1430 - WT,QX1430 | 223,16 | 31,82 | 709 | 7,01 | <0,0001 |
| WT,CB4856 - WT,ED3005 | -154,30 | 45,67 | 709 | -3,38 | 8,67E-02 |
| WT,CB4856 - WT,JU1200 | -182,13 | 49,45 | 709 | -3,68 | 3,30E-02 |
| WT,CB4856 - WT,JU2829 | -326,44 | 45,14 | 709 | -7,23 | <0,0001 |
| WT,CB4856 - WT,JU3166 | -198,23 | 49,45 | 709 | -4,01 | 1,02E-02 |
| WT,CB4856 - WT,JU751 | -265,39 | 45,14 | 709 | -5,88 | 1,19E-06 |
| WT,CB4856 - WT,JU830 | -169,96 | 43,14 | 709 | -3,94 | 1,32E-02 |
| WT,CB4856 - WT,N2 | -323,35 | 41,38 | 709 | -7,82 | <0,0001 |
| WT,CB4856 - WT,QG2873 | -148,26 | 49,99 | 709 | -2,97 | 2,53E-01 |
| WT,CB4856 - WT,QX1430 | -158,29 | 41,48 | 709 | -3,82 | 2,07E-02 |
| WT,ED3005 - WT,JU1200 | -27,82 | 45,67 | 709 | -0,61 | 1,00E+00 |
| WT,ED3005 - WT,JU2829 | -172,14 | 40,97 | 709 | -4,20 | 4,75E-03 |
| WT,ED3005 - WT,JU3166 | -43,93 | 45,67 | 709 | -0,96 | 1,00E+00 |
| WT,ED3005 - WT,JU751 | -111,09 | 40,97 | 709 | -2,71 | 4,21E-01 |
| WT,ED3005 - WT,JU830 | -15,66 | 38,74 | 709 | -0,40 | 1,00E+00 |
| WT,ED3005 - WT,N2 | -169,05 | 36,77 | 709 | -4,60 | 8,72E-04 |
| WT,ED3005 - WT,QG2873 | 6,05 | 46,25 | 709 | 0,13 | 1,00E+00 |
| WT,ED3005 - WT,QX1430 | -3,99 | 36,89 | 709 | -0,11 | 1,00E+00 |
| WT,JU1200 - WT,JU2829 | -144,32 | 45,14 | 709 | -3,20 | 1,44E-01 |
| WT,JU1200 - WT,JU3166 | -16,10 | 49,45 | 709 | -0,33 | 1,00E+00 |
| WT,JU1200 - WT,JU751 | -83,26 | 45,14 | 709 | -1,84 | 9,52E-01 |
| WT,JU1200 - WT,JU830 | 12,16 | 43,14 | 709 | 0,28 | 1,00E+00 |
| WT,JU1200 - WT,N2 | -141,23 | 41,38 | 709 | -3,41 | 7,82E-02 |

| | | | | | |
|---|---|---|---|---|---|
| WT,JU1200 - WT,QG2873 | 33,87 | 49,99 | 709 | 0,68 | 1,00E+00 |
| WT,JU1200 - WT,QX1430 | 23,83 | 41,48 | 709 | 0,57 | 1,00E+00 |
| WT,JU2829 - WT,JU3166 | 128,22 | 45,14 | 709 | 2,84 | 3,31E-01 |
| WT,JU2829 - WT,JU751 | 61,06 | 40,38 | 709 | 1,51 | 9,94E-01 |
| WT,JU2829 - WT,JU830 | 156,48 | 38,12 | 709 | 4,10 | 7,00E-03 |
| WT,JU2829 - WT,N2 | 3,09 | 36,12 | 709 | 0,09 | 1,00E+00 |
| WT,JU2829 - WT,QG2873 | 178,19 | 45,73 | 709 | 3,90 | 1,55E-02 |
| WT,JU2829 - WT,QX1430 | 168,15 | 36,23 | 709 | 4,64 | 7,14E-04 |
| WT,JU3166 - WT,JU751 | -67,16 | 45,14 | 709 | -1,49 | 9,95E-01 |
| WT,JU3166 - WT,JU830 | 28,27 | 43,14 | 709 | 0,66 | 1,00E+00 |
| WT,JU3166 - WT,N2 | -125,12 | 41,38 | 709 | -3,02 | 2,22E-01 |
| WT,JU3166 - WT,QG2873 | 49,97 | 49,99 | 709 | 1,00 | 1,00E+00 |
| WT,JU3166 - WT,QX1430 | 39,93 | 41,48 | 709 | 0,96 | 1,00E+00 |
| WT,JU751 - WT,JU830 | 95,43 | 38,12 | 709 | 2,50 | 5,82E-01 |
| WT,JU751 - WT,N2 | -57,96 | 36,12 | 709 | -1,60 | 9,89E-01 |
| WT,JU751 - WT,QG2873 | 117,13 | 45,73 | 709 | 2,56 | 5,36E-01 |
| WT,JU751 - WT,QX1430 | 107,09 | 36,23 | 709 | 2,96 | 2,59E-01 |
| WT,JU830 - WT,N2 | -153,39 | 33,57 | 709 | -4,57 | 9,89E-04 |
| WT,JU830 - WT,QG2873 | 21,71 | 43,75 | 709 | 0,50 | 1,00E+00 |
| WT,JU830 - WT,QX1430 | 11,67 | 33,70 | 709 | 0,35 | 1,00E+00 |
| WT,N2 - WT,QG2873 | 175,09 | 42,01 | 709 | 4,17 | 5,45E-03 |
| WT,N2 - WT,QX1430 | 165,06 | 31,41 | 709 | 5,26 | 3,59E-05 |
| WT,QG2873 - WT,QX1430 | -10,04 | 42,11 | 709 | -0,24 | 1,00E+00 |

**Table S6 (accompanies Fig. S5B).** Results for statistical analyses testing for the effects of and interactions between *State* (presence versus absence of *mod-5* (*lf*)) and *Background* (strain) on egg laying when exposed to 25mM serotonin (Align Rank Transform ANOVA).

| Term | Df | Sum sq | F value | Pr(>F) | Sig |
|---|---|---|---|---|---|
| State | 1 | 8774,82 | 0,29 | 5,92E-01 | ns |
| Background | 7 | 13709361,51 | 277,73 | 6,93E-181 | *** |
| State x Background | 7 | 732093,79 | 3,57 | 8,99E-04 | *** |

**Contrasts**

| Contrast | Estimate | Se | Df | t-ratio | p-value |
|---|---|---|---|---|---|
| mod-5,ED3005 - mod-5,JU2829 | 431,32 | 19,69 | 582 | 21,90 | 5,06E-10 |
| mod-5,ED3005 - mod-5,JU3166 | 101,79 | 22,02 | 582 | 4,62 | 5,16E-04 |
| mod-5,ED3005 - mod-5,JU751 | 392,48 | 19,43 | 582 | 20,20 | 5,06E-10 |
| mod-5,ED3005 - mod-5,JU830 | 5,26 | 18,51 | 582 | 0,28 | 1,00E+00 |
| mod-5,ED3005 - mod-5,N2 | 226,21 | 19,43 | 582 | 11,64 | 5,06E-10 |
| mod-5,ED3005 - mod-5,QG2873 | 258,17 | 22,02 | 582 | 11,72 | 5,06E-10 |
| mod-5,ED3005 - mod-5,QX1430 | 193,76 | 17,67 | 582 | 10,96 | 5,06E-10 |
| mod-5,ED3005 - WT,ED3005 | 71,07 | 19,69 | 582 | 3,61 | 2,90E-02 |
| mod-5,ED3005 - WT,JU2829 | 452,31 | 19,69 | 582 | 22,97 | 5,06E-10 |
| mod-5,ED3005 - WT,JU3166 | 77,85 | 22,02 | 582 | 3,54 | 3,70E-02 |
| mod-5,ED3005 - WT,JU751 | 394,53 | 19,83 | 582 | 19,89 | 5,06E-10 |
| mod-5,ED3005 - WT,JU830 | -12,54 | 18,68 | 582 | -0,67 | 1,00E+00 |
| mod-5,ED3005 - WT,N2 | 169,87 | 19,69 | 582 | 8,63 | 5,06E-10 |
| mod-5,ED3005 - WT,QG2873 | 295,62 | 22,02 | 582 | 13,43 | 5,06E-10 |
| mod-5,ED3005 - WT,QX1430 | 195,98 | 17,61 | 582 | 11,13 | 5,06E-10 |
| mod-5,JU2829 - mod-5,JU3166 | -329,53 | 22,02 | 582 | -14,97 | 5,06E-10 |
| mod-5,JU2829 - mod-5,JU751 | -38,84 | 19,43 | 582 | -2,00 | 8,25E-01 |
| mod-5,JU2829 - mod-5,JU830 | -426,06 | 18,51 | 582 | -23,02 | 5,06E-10 |
| mod-5,JU2829 - mod-5,N2 | -205,11 | 19,43 | 582 | -10,55 | 5,06E-10 |
| mod-5,JU2829 - mod-5,QG2873 | -173,15 | 22,02 | 582 | -7,86 | 5,08E-10 |
| mod-5,JU2829 - mod-5,QX1430 | -237,56 | 17,67 | 582 | -13,44 | 5,06E-10 |
| mod-5,JU2829 - WT,ED3005 | -360,25 | 19,69 | 582 | -18,29 | 5,06E-10 |
| mod-5,JU2829 - WT,JU2829 | 20,99 | 19,69 | 582 | 1,07 | 1,00E+00 |
| mod-5,JU2829 - WT,JU3166 | -353,47 | 22,02 | 582 | -16,05 | 5,06E-10 |
| mod-5,JU2829 - WT,JU751 | -36,78 | 19,83 | 582 | -1,85 | 8,95E-01 |
| mod-5,JU2829 - WT,JU830 | -443,86 | 18,68 | 582 | -23,76 | 5,06E-10 |
| mod-5,JU2829 - WT,N2 | -261,44 | 19,69 | 582 | -13,28 | 5,06E-10 |
| mod-5,JU2829 - WT,QG2873 | -135,69 | 22,02 | 582 | -6,16 | 1,60E-07 |
| mod-5,JU2829 - WT,QX1430 | -235,34 | 17,61 | 582 | -13,36 | 5,06E-10 |
| mod-5,JU3166 - mod-5,JU751 | 290,68 | 21,79 | 582 | 13,34 | 5,06E-10 |
| mod-5,JU3166 - mod-5,JU830 | -96,53 | 20,96 | 582 | -4,60 | 5,60E-04 |
| mod-5,JU3166 - mod-5,N2 | 124,42 | 21,79 | 582 | 5,71 | 2,12E-06 |
| mod-5,JU3166 - mod-5,QG2873 | 156,38 | 24,12 | 582 | 6,48 | 2,34E-08 |
| mod-5,JU3166 - mod-5,QX1430 | 91,97 | 20,23 | 582 | 4,55 | 7,29E-04 |
| mod-5,JU3166 - WT,ED3005 | -30,72 | 22,02 | 582 | -1,40 | 9,91E-01 |
| mod-5,JU3166 - WT,JU2829 | 350,51 | 22,02 | 582 | 15,92 | 5,06E-10 |
| mod-5,JU3166 - WT,JU3166 | -23,94 | 24,12 | 582 | -0,99 | 1,00E+00 |
| mod-5,JU3166 - WT,JU751 | 292,74 | 22,14 | 582 | 13,22 | 5,06E-10 |
| mod-5,JU3166 - WT,JU830 | -114,33 | 21,12 | 582 | -5,41 | 1,06E-05 |

| | | | | | |
|---|---|---|---|---|---|
| mod-5,JU3166 - WT,N2 | 68,08 | 22,02 | 582 | 3,09 | 1,36E-01 |
| mod-5,JU3166 - WT,QG2873 | 193,83 | 24,12 | 582 | 8,04 | 5,07E-10 |
| mod-5,JU3166 - WT,QX1430 | 94,19 | 20,18 | 582 | 4,67 | 4,22E-04 |
| mod-5,JU751 - mod-5,JU830 | -387,22 | 18,23 | 582 | -21,24 | 5,06E-10 |
| mod-5,JU751 - mod-5,N2 | -166,26 | 19,17 | 582 | -8,67 | 5,06E-10 |
| mod-5,JU751 - mod-5,QG2873 | -134,31 | 21,79 | 582 | -6,17 | 1,58E-07 |
| mod-5,JU751 - mod-5,QX1430 | -198,72 | 17,38 | 582 | -11,43 | 5,06E-10 |
| mod-5,JU751 - WT,ED3005 | -321,41 | 19,43 | 582 | -16,54 | 5,06E-10 |
| mod-5,JU751 - WT,JU2829 | 59,83 | 19,43 | 582 | 3,08 | 1,41E-01 |
| mod-5,JU751 - WT,JU3166 | -314,62 | 21,79 | 582 | -14,44 | 5,06E-10 |
| mod-5,JU751 - WT,JU751 | 2,06 | 19,58 | 582 | 0,11 | 1,00E+00 |
| mod-5,JU751 - WT,JU830 | -405,02 | 18,41 | 582 | -22,00 | 5,06E-10 |
| mod-5,JU751 - WT,N2 | -222,60 | 19,43 | 582 | -11,45 | 5,06E-10 |
| mod-5,JU751 - WT,QG2873 | -96,85 | 21,79 | 582 | -4,45 | 1,14E-03 |
| mod-5,JU751 - WT,QX1430 | -196,49 | 17,32 | 582 | -11,34 | 5,06E-10 |
| mod-5,JU830 - mod-5,N2 | 220,95 | 18,23 | 582 | 12,12 | 5,06E-10 |
| mod-5,JU830 - mod-5,QG2873 | 252,91 | 20,96 | 582 | 12,06 | 5,06E-10 |
| mod-5,JU830 - mod-5,QX1430 | 188,50 | 16,34 | 582 | 11,54 | 5,06E-10 |
| mod-5,JU830 - WT,ED3005 | 65,81 | 18,51 | 582 | 3,56 | 3,46E-02 |
| mod-5,JU830 - WT,JU2829 | 447,05 | 18,51 | 582 | 24,16 | 5,06E-10 |
| mod-5,JU830 - WT,JU3166 | 72,59 | 20,96 | 582 | 3,46 | 4,67E-02 |
| mod-5,JU830 - WT,JU751 | 389,27 | 18,65 | 582 | 20,87 | 5,06E-10 |
| mod-5,JU830 - WT,JU830 | -17,80 | 17,43 | 582 | -1,02 | 1,00E+00 |
| mod-5,JU830 - WT,N2 | 164,62 | 18,51 | 582 | 8,90 | 5,06E-10 |
| mod-5,JU830 - WT,QG2873 | 290,37 | 20,96 | 582 | 13,85 | 5,06E-10 |
| mod-5,JU830 - WT,QX1430 | 190,72 | 16,28 | 582 | 11,72 | 5,06E-10 |
| mod-5,N2 - mod-5,QG2873 | 31,95 | 21,79 | 582 | 1,47 | 9,86E-01 |
| mod-5,N2 - mod-5,QX1430 | -32,45 | 17,38 | 582 | -1,87 | 8,90E-01 |
| mod-5,N2 - WT,ED3005 | -155,14 | 19,43 | 582 | -7,98 | 5,07E-10 |
| mod-5,N2 - WT,JU2829 | 226,09 | 19,43 | 582 | 11,63 | 5,06E-10 |
| mod-5,N2 - WT,JU3166 | -148,36 | 21,79 | 582 | -6,81 | 3,43E-09 |
| mod-5,N2 - WT,JU751 | 168,32 | 19,58 | 582 | 8,60 | 5,06E-10 |
| mod-5,N2 - WT,JU830 | -238,75 | 18,41 | 582 | -12,97 | 5,06E-10 |
| mod-5,N2 - WT,N2 | -56,34 | 19,43 | 582 | -2,90 | 2,19E-01 |
| mod-5,N2 - WT,QG2873 | 69,41 | 21,79 | 582 | 3,19 | 1,06E-01 |
| mod-5,N2 - WT,QX1430 | -30,23 | 17,32 | 582 | -1,75 | 9,34E-01 |
| mod-5,QG2873 - mod-5,QX1430 | -64,41 | 20,23 | 582 | -3,18 | 1,07E-01 |
| mod-5,QG2873 - WT,ED3005 | -187,10 | 22,02 | 582 | -8,50 | 5,06E-10 |
| mod-5,QG2873 - WT,JU2829 | 194,14 | 22,02 | 582 | 8,82 | 5,06E-10 |
| mod-5,QG2873 - WT,JU3166 | -180,31 | 24,12 | 582 | -7,48 | 5,40E-10 |
| mod-5,QG2873 - WT,JU751 | 136,37 | 22,14 | 582 | 6,16 | 1,64E-07 |
| mod-5,QG2873 - WT,JU830 | -270,71 | 21,12 | 582 | -12,82 | 5,06E-10 |
| mod-5,QG2873 - WT,N2 | -88,29 | 22,02 | 582 | -4,01 | 6,81E-03 |
| mod-5,QG2873 - WT,QG2873 | 37,46 | 24,12 | 582 | 1,55 | 9,76E-01 |
| mod-5,QG2873 - WT,QX1430 | -62,18 | 20,18 | 582 | -3,08 | 1,40E-01 |
| mod-5,QX1430 - WT,ED3005 | -122,69 | 17,67 | 582 | -6,94 | 1,74E-09 |
| mod-5,QX1430 - WT,JU2829 | 258,55 | 17,67 | 582 | 14,63 | 5,06E-10 |
| mod-5,QX1430 - WT,JU3166 | -115,90 | 20,23 | 582 | -5,73 | 1,91E-06 |
| mod-5,QX1430 - WT,JU751 | 200,78 | 17,83 | 582 | 11,26 | 5,06E-10 |
| mod-5,QX1430 - WT,JU830 | -206,30 | 16,54 | 582 | -12,48 | 5,06E-10 |

| | | | | | |
|---|---|---|---|---|---|
| mod-5,QX1430 - WT,N2 | -23,88 | 17,67 | 582 | -1,35 | 9,94E-01 |
| mod-5,QX1430 - WT,QG2873 | 101,87 | 20,23 | 582 | 5,04 | 7,32E-05 |
| mod-5,QX1430 - WT,QX1430 | 2,23 | 15,32 | 582 | 0,15 | 1,00E+00 |
| WT,ED3005 - WT,JU2829 | 381,24 | 19,69 | 582 | 19,36 | 5,06E-10 |
| WT,ED3005 - WT,JU3166 | 6,78 | 22,02 | 582 | 0,31 | 1,00E+00 |
| WT,ED3005 - WT,JU751 | 323,47 | 19,83 | 582 | 16,31 | 5,06E-10 |
| WT,ED3005 - WT,JU830 | -83,61 | 18,68 | 582 | -4,48 | 9,98E-04 |
| WT,ED3005 - WT,N2 | 98,81 | 19,69 | 582 | 5,02 | 8,02E-05 |
| WT,ED3005 - WT,QG2873 | 224,56 | 22,02 | 582 | 10,20 | 5,06E-10 |
| WT,ED3005 - WT,QX1430 | 124,91 | 17,61 | 582 | 7,09 | 9,69E-10 |
| WT,JU2829 - WT,JU3166 | -374,45 | 22,02 | 582 | -17,01 | 5,06E-10 |
| WT,JU2829 - WT,JU751 | -57,77 | 19,83 | 582 | -2,91 | 2,12E-01 |
| WT,JU2829 - WT,JU830 | -464,85 | 18,68 | 582 | -24,88 | 5,06E-10 |
| WT,JU2829 - WT,N2 | -282,43 | 19,69 | 582 | -14,34 | 5,06E-10 |
| WT,JU2829 - WT,QG2873 | -156,68 | 22,02 | 582 | -7,12 | 9,00E-10 |
| WT,JU2829 - WT,QX1430 | -256,32 | 17,61 | 582 | -14,55 | 5,06E-10 |
| WT,JU3166 - WT,JU751 | 316,68 | 22,14 | 582 | 14,30 | 5,06E-10 |
| WT,JU3166 - WT,JU830 | -90,40 | 21,12 | 582 | -4,28 | 2,30E-03 |
| WT,JU3166 - WT,N2 | 92,02 | 22,02 | 582 | 4,18 | 3,48E-03 |
| WT,JU3166 - WT,QG2873 | 217,77 | 24,12 | 582 | 9,03 | 5,06E-10 |
| WT,JU3166 - WT,QX1430 | 118,13 | 20,18 | 582 | 5,85 | 9,55E-07 |
| WT,JU751 - WT,JU830 | -407,08 | 18,83 | 582 | -21,62 | 5,06E-10 |
| WT,JU751 - WT,N2 | -224,66 | 19,83 | 582 | -11,33 | 5,06E-10 |
| WT,JU751 - WT,QG2873 | -98,91 | 22,14 | 582 | -4,47 | 1,04E-03 |
| WT,JU751 - WT,QX1430 | -198,55 | 17,77 | 582 | -11,17 | 5,06E-10 |
| WT,JU830 - WT,N2 | 182,42 | 18,68 | 582 | 9,76 | 5,06E-10 |
| WT,JU830 - WT,QG2873 | 308,17 | 21,12 | 582 | 14,59 | 5,06E-10 |
| WT,JU830 - WT,QX1430 | 208,52 | 16,48 | 582 | 12,66 | 5,06E-10 |
| WT,N2 - WT,QG2873 | 125,75 | 22,02 | 582 | 5,71 | 2,12E-06 |
| WT,N2 - WT,QX1430 | 26,11 | 17,61 | 582 | 1,48 | 9,84E-01 |
| WT,QG2873 - WT,QX1430 | -99,64 | 20,18 | 582 | -4,94 | 1,18E-04 |

**Table S7.** List of reagents used.

| | |
|---|---|
| crRNA and single strand DNA oligo used for *mod-5* (n822) loss-of-function | |
| crRNA crNB040 | GAAGUUCCGUGGGCGUCAUG |
| oNB322 | TCTAGATATTGTCACGTTGAGGTCATCTGAGCATCTCGGaGTgTTCCAgGGgTTtgCtCATGACGCCCACGG |
| Primers used for *mod-5* (n822) loss-of-function PCR screen | |
| oNB325 | GCGTCATGaGcaAAcCCc |
| oNB327 | ATCATCGCTCAAGCCGTCTA |
| oNB328 | CAGACGACTGTGGACCCTTC |

#### Additional supporting information

**Description of separate file Data S1 (raw data in Excel format). Numbers below refer to numbering of worksheets in the Excel file and corresponding figure numbers.**

1. Number of offspring *in utero* in hermaphrodites (L4 + 48h) in 316 *C. elegans* wild strains (Fig. 1A).

2. Geographic distribution of 316 C. elegans wild strains (Fig. 1B).

3. Single-marker based GWA mapping region for mean number of eggs *in utero.* (N = 316) (Fig. 1D).

4. Single-marker based GWA mapping region for the coefficient of variation (*CV*) (mean number of eggs *in utero*). N = 316) (Fig. 1E).

5. Egg retention at L4 + 30h in a subset of 15 strains with divergent egg retention, divided into the three phenotypic classes. Class I *weak*: < 10 eggs *in utero* (N = 34), Class II *canonical*: 10-25 eggs *in utero* (N = 230), Class III *strong*: > 25 eggs *in utero* (N = 14). Class III strains were further distinguished depending on the absence (Class IIIA) or presence (Class IIIB) of the KNCL-1 V530L variant explaining strong egg retention (Vigne et al., 2021). Embryonic stages were divided into five age groups: 1-2 cell stage, 4-26 cell stage, 44-cell to gastrula stage, bean to two-fold stage, three-fold stage, L1 larva. (Fig. 2A, 2D and S3A).

6. Geographic distribution of the 15 focal strains with divergent egg retention (Fig. 2B).

7. Temporal dynamics of offspring number in utero in the 15 focal strains. Number of eggs and larvae in utero at three stages covering the reproductive span of self-fertilizing hermaphrodites. (Fig. 2C and 2E).

8. Temporal dynamics of egg-laying activity in the 15 focal strains. Number of eggs laid (within a 2-hour window) at five time points across the reproductive span of self-fertilizing hermaphrodites (Fig. 3A).

9. Natural variation in temporal patterns of egg-laying behaviour. Temporal tracking of egg-laying events in individual hermaphrodites (at L4 + 30h) of the 15 focal strains during a three-hour interval (Fig. 3C and S4A).

10. Natural variation in temporal patterns of egg-laying behaviour. Temporal tracking of egg-laying events in individual hermaphrodites (at L4 + 30h) of the 15 focal strains during a three-hour interval (Fig. 3D-F and S4B).

11. Time between first fertilization and first egg-laying event and number of eggs in utero at first egg-laying event (Fig. 3G and 3H).

12. Natural variation in egg-laying activity in response to exogenous serotonin (25mM) compared to M9 buffer without food (control) (Fig. 4B).

13. Natural variation in egg-laying activity in response to exogenous imipramine (2.36mM) compared to M9 buffer without food (control) (Fig. 4B).

14. Natural variation in egg-laying activity in response to exogenous fluoxetine (1.5mM) compared to M9 buffer without food (control) (Fig. 4B).

15. Effects of exogenous serotonin, fluoxetine and imipramine on egg-laying activity in strains with strongly divergent egg retention due to variation in a single amino acid residue of KCNL-1 (Fig. 4C and S5A).

16. Natural variation in the effect of *mod-5(n822)* on egg laying in M9 (control) (Fig. 5B) and serotonin (25mM) (Fig. S5B).

17. Natural variation in egg laying in response to different serotonin concentrations in wild type and *mod-5(n822)* animals (Fig. 5C and S5C).

18. Lifetime production in self-fertilizing hermaphrodites of the 15 focal strains with divergent egg retention (Fig. 6A and 6B).

19. Correlations between mean egg retention and mean lifetime offspring production (Fig. 6C) and mean percentage of survival (day 5) across the 15 focal strains (Fig. 6E).

20. Hermaphrodite survival in the 15 focal strains (Fig. 6D).

21. Temporal progression of internal (matricidal) hatching during the survival assay (Fig. 6D) in the 15 focal strains, measured as the cumulative percentage of dead mothers containing one or more internally hatched larva (Fig. 6F).

22. Hermaphrodite self-sperm number and number of internally hatched larvae in select strains (Fig. 6G).

23. Age distribution of embryos contained within eggs laid by hermaphrodites (L4+40h) of the 15 focal strains (Fig. 7A).

24. Variation in the time until hatching of laid eggs among the 15 focal strains with divergent egg retention. The fraction of hatched eggs was scored every hour until all eggs had hatched (Fig. 7B).

25. Variation in the egg-adult developmental time among the 15 focal strains with divergent egg retention. Eggs were allowed to hatch and after 45 hours of development, populations were surveyed every 1-2 hours to count the fraction of reproductively mature adults, i.e. adults with one or two eggs in utero. (Fig. 7C).

26. Short-term competition of $JU1200_{WT}$ and $JU1200_{KCNL\text{-}1V530}$ against a GFP-tester strain (myo-2::gfp) with genotype starting frequencies of 50:50 (Fig. 7D).

27. Short- term competition of JU751 against each of five wild strains isolated from the same locality (Fig. 7E).

28. Differences in survival of eggs developing ex utero versus in utero when exposed to a high concentration of ethanol (96%, 10 minute-exposure) (Fig. 8A and S7A).

29. Differences in survival of eggs developing ex utero versus in utero when exposed to a high concentration of acetic acid (10M, 15 minute-exposure) (Fig. 8B and S7B).

30. Differences in survival of eggs developing ex utero versus in utero when exposed to a high concentration of acetic acid (10M, 15 minute-exposure). Comparison of the strain JU751 (Class III) to four strains (Class II) isolated from the same locality (Fig. 8C and Fig. S7C).

31. Differences in survival of eggs developing ex utero versus in utero when exposed to a high concentration of acetic acid (10M, 15 minute-exposure). Comparison between the Class II strain JU1200 and the CRISPR-engineered JU1200KCNL-1(V530L) (Fig. 8D).

32. Differences in survival of larvae derived from *ex utero* versus *in utero* eggs exposed to acetic acid (strain JU751) (Fig. 8E).

33. Effects of osmotic stress on survival of embryos from eggs developing ex utero in the strain JU751. Embryonic survival was estimated by counting the fraction of live larvae 24 hours later (Fig. 8F).

34. Differences in self-fertility of JU751 animals derived from surviving eggs developing ex utero versus in utero when exposed to mild osmotic stress (0.3M NaCl) for 15 hours (Fig. 8G).

35. Differences in developmental time of JU751 animals derived from surviving eggs ex utero versus in utero when exposed to a low concentration of acetic acid (1M) for 15 minutes; each stage was assigned a score of development as follows: L3 = 1; early mid-L4 = 2; midL4 = 3; lateL4 = 4; Adult = 5. (Fig. 8H).

36. Differences in total lifetime offspring production of selfing JU751 animals derived from eggs *ex utero* versus eggs and L1 larvae *in utero* when exposed to a low concentration of acetic acid (1M) for 15 minutes; in parallel, *ex utero* eggs (dissected from adults at L4 + 36h) were exposed to a control treatment (M9 buffer) (Fig. 8I).

37. Sampling details of the 316 examined *C. elegans* strains (geographic distribution, substrate category and haplotype information, and mean egg retention (Fig. S1 and S2). Sampling information were obtained from CeNDR (https://www.elegansvariation.org/) (Cook et al., 2017).

38. Egg size measurements of the 15 focal strains (Fig. S3B).

39. Body size measurements of the 15 focal strains; selfing hermaphrodites at first fertilization (Fig. S3C).

40. Correlations between size measurements (body volume, egg volume) and mean egg number *in utero* across the 15 focal strains (Fig. S3D to S3G).

41. Total self-sperm number and lifetime offspring production of the Class IIIB strain JU2593 (Fig. S6A).

42. Number of internally hatched larvae and gonad/germline damage in the Class IIIB strain JU2593 (Fig.S6B).

43. Number of self-sperm and unfertilized oocytes *in utero* in JU830 and NIC1832 strains uterus (Fig. S6C and S6D).
